# A synthetic RNA-based biosensor for fructose-1,6-bisphosphate that reports glycolytic flux

**DOI:** 10.1101/2020.10.11.335109

**Authors:** Alvaro D. Ortega, Vakil Takhaveev, Silke Bonsing-Vedelaar, Yi Long, Neus Mestre-Farràs, Danny Incarnato, Franziska Ersoy, Lars Folke Olsen, Günter Mayer, Matthias Heinemann

## Abstract

Metabolic heterogeneity, the occurrence of different metabolic phenotypes among cells, represents a key challenge in health and biotechnology. To unravel its molecular basis, tools probing metabolism of single cells are needed. While RNA devices harbor huge potential for the development of such tools, until today, it is challenging to create *in vivo*-functional sensors for any given metabolite. Here, we developed from scratch an RNA-based sensor for fructose-1,6-bisphosphate (FBP), a doubly phosphorylated intermediate of glycolysis. Starting from *in vitro* selection of an RNA aptamer and its structural analyses, we developed libraries of RNA-based regulatory devices with this aptamer and the hammerhead ribozyme as an actuator. Through FACS-seq-based high-throughput screening in yeast, we identified *in vivo-*functional FBP-sensing devices that generate fluorescent readout dependent on intracellular FBP concentration. As FBP reports the flux through glycolysis, the developed RNA device can be used to sense the glycolytic rate in single cells, offering unprecedented possibilities to investigate the causes of metabolic heterogeneity.

## INTRODUCTION

To cope with changing environments and intracellular demands, cellular metabolism can be dynamically reshaped. Recent research has shown that this so-called metabolic plasticity can lead to coexistence of different metabolic phenotypes within the same population (1–5). Such metabolic heterogeneity presents significant challenges for biotechnology (6) and biomedicine, especially in cancer research (7). To investigate the molecular basis of metabolic heterogeneity, tools need to be developed that can probe metabolism on the single-cell level (8).

Of particular interest would be single-cell-level tools that could sense the flux through glycolysis – a pathway with high relevance in biotechnology and cancer research. While ‘metabolic fluxes’ cannot be directly measured, certain intracellular metabolites function as physiological flux signals. That is, the concentration of such flux-signaling metabolites correlates with the flux through the respective pathway (9, 10). The glycolytic intermediate fructose-1,6-bisphosphate (FBP) is a glycolytic flux-signaling metabolite in organisms across kingdoms. FBP levels, on the one hand, correlate with the flux through glycolysis and, on the other, regulate enzyme activity and gene expression in response to changes in glycolytic flux (10). In this regard, FBP allosterically regulates the CggR and CcpA transcription factors in *Bacillus subtilis* (11, 12), it activates RAS-signaling in yeast (13), and in mammalian cells, it can stimulate the activation of AMPK through an AMP/ADP independent mechanism (14). Given its conserved role as a natural sensor of glycolytic flux, FBP thus poses a suitable indicator of glycolytic flux to unveil metabolic heterogeneity among cells.

RNA-based gene regulatory devices, or synthetic riboswitches, have been put forward as molecular tools for monitoring intracellular concentrations of particular metabolites (15). Such RNA-based tools are modular: they contain an aptamer as a sensing domain where metabolite binding induces a conformational change that is transmitted to an actuator domain (*e*.*g*. a ribozyme or an expression platform) which in turn triggers a specific output (16). In these ligand-dependent regulatory systems, actuators rely on different post-transcriptional mechanisms that control the expression of a (reporter) gene (17–23). A different concept for RNA-based metabolite sensing exploits fluorogenic aptamers, such as *Spinach*, which light up as a result of ligand binding (24–26). However, while the discovery of natural riboswitches in bacteria and pioneering work on modular allosteric ribozymes raised huge expectations for the engineering of RNA devices that could respond to any effector of interest (27), to date such RNA devices have only been accomplished for a very limited set of metabolites.

Most likely, there are several reasons for this situation. First, while aptamers can in principle be selected *in vitro* against any target molecule, obtaining aptamers that are functional in cells is a major challenge (28). In fact, to our knowledge, only a handful of *in vitro*-selected aptamers for small molecules has been functionally validated as sensing domains in cells: (i) three antibiotic-binding aptamers obtained through a combination of *in vitro* selection and screening in yeast (17, 18, 29); (ii) two aptamers that were raised against aromatic amino acids using RNA pools based on structural scaffolds derived from natural riboswitches (30); and (iii) the aptamers for theophylline and ATP obtained by conventional SELEX and shown to be directly functional in cells (24, 31). Besides, high-throughput screening of RNA-device libraries containing aptamers previously found to be functional in cells has led to sensors with improved sensitivity and/or dynamic response (32–37). However, such *in vivo* screening has not been utilized for *de novo* selection of aptamers, likely because the enormous size of the search space cannot be comprehensively screened due to the limited transformation efficiency. The second challenge for the development of RNA-based sensors for cellular metabolites stems from the fact that the intracellular concentration of a target metabolite is set physiologically. Hence, only aptamers with an affinity within the physiological concentration range would be suitable for RNA devices detecting metabolite changes in the cell. Moreover, binary screening conditions *in vivo* can be challenging because intracellular concentrations of cell-intrinsic metabolites might not be easily controlled externally. The third challenge is that concentrations of intracellular metabolites are typically in the micro to millimolar range (38) while at the same time chemically similar metabolites are also present, which requires the sensing domain to respond to a highly concentrated ligand in a yet selective manner.

Here, tackling these three challenges, we accomplished the development of an RNA-based device to sense the intracellular metabolite fructose-1,6-bisphosphate (FBP). Specifically, we selected *in vitro* an aptamer that binds FBP under physiological conditions, designed RNA-based regulatory device libraries with a hammerhead ribozyme as an actuator, and used a high-throughput screening in budding yeast to identify RNA devices that generate an FBP concentration-dependent ratiometric fluorescent readout. In single-cell microscopic analyses, we demonstrate that the identified FBP sensor can differentiate cells with different glycolytic fluxes within clonal yeast populations. The potential of the developed sensor to monitor glycolytic flux *in vivo* opens exciting applications for fundamental research into metabolic heterogeneity, as well as novel applications in biotechnology such as activity-based screening.

## RESULTS

### Selection of an FBP-binding RNA aptamer

To generate an FBP-sensing domain for our envisioned RNA device that reports intracellular FBP levels, and thus glycolytic flux, we first selected an FBP-binding RNA aptamer through systematic evolution of ligands by exponential enrichment (SELEX). After immobilizing FBP on epoxide-sepharose, we used an RNA library with the randomized region of 40 nucleotides flanked by constant regions of 15 and 25 nucleotides, respectively. To mimic the physiological conditions of the cytoplasm, we employed a phosphate buffer rich in K^+^ and with a high concentration of glucose (to account for the presence of cytosolic metabolites) and added to it 5 mM Mg^2+^ to facilitate the interaction of the doubly phosphorylated FBP with the anionic core of RNA (39). After 13 selection cycles, cloning and sequencing, we obtained ten different sequences, all of which showed a shortened variable region compared to the original library (Supplementary Table S1), consistent with agarose gel electrophoresis analysis of the pools revealing a significantly reduced dsDNA length from selection round 9 onward (40). Sequence analysis demonstrated highly similar clones, with an enriched motif, WUCCU, at the 5’ end of the variable region followed by a short nucleotide stretch (up to 9 nucleotides), which allowed us to categorize the sequences into three sets (Supplementary Table S1).

To assess the affinity and specificity of the selected clones to FBP, fluorescently labeled (CAL Fluor Red 610) RNA clones 34, 45 and 58 (*i*.*e*. representatives of each of the three sets of clones, Supplementary Table S1) were incubated with sepharose columns with immobilized FBP, and then eluted with solutions of different FBP concentrations. We found that all clones bound FBP within the low mM concentration range (Fig. 1A and Supplementary Fig. S1A-B), which is the physiological concentration range of FBP in yeast (9). A mutant in the constant region that disrupted a stem structure predicted for these clones by Mfold energy minimization (nucleotide stretch C13 to A21, Supplementary Fig. S2A-D) required a 10-fold higher FBP concentration for elution (Fig. 1A). Other phosphorylated intracellular metabolites (glucose-6-phosphate, fructose-6-phosphate, and ATP) failed to elute the RNA from the FBP column (Fig. 1B and Supplementary Fig. S1C-D). These results suggested that our selected clones are aptamers that selectively bind FBP in the physiological mM range. The fact that the clones C34, C45 and C58 displayed a similar affinity and selectivity (Fig. 1A-B and Supplementary Fig. S1) as well as they all had a predicted structural stem-loop motif according to Mfold energy minimization (Supplementary Fig. S2A-C) suggested that the nucleotide sequence downstream the WUCCU motif (Supplementary Table S1) is not essential for binding, indicating that the three clones are functionally equivalent. Thus, we continued our work with one clone, C45.

**Figure 1.**
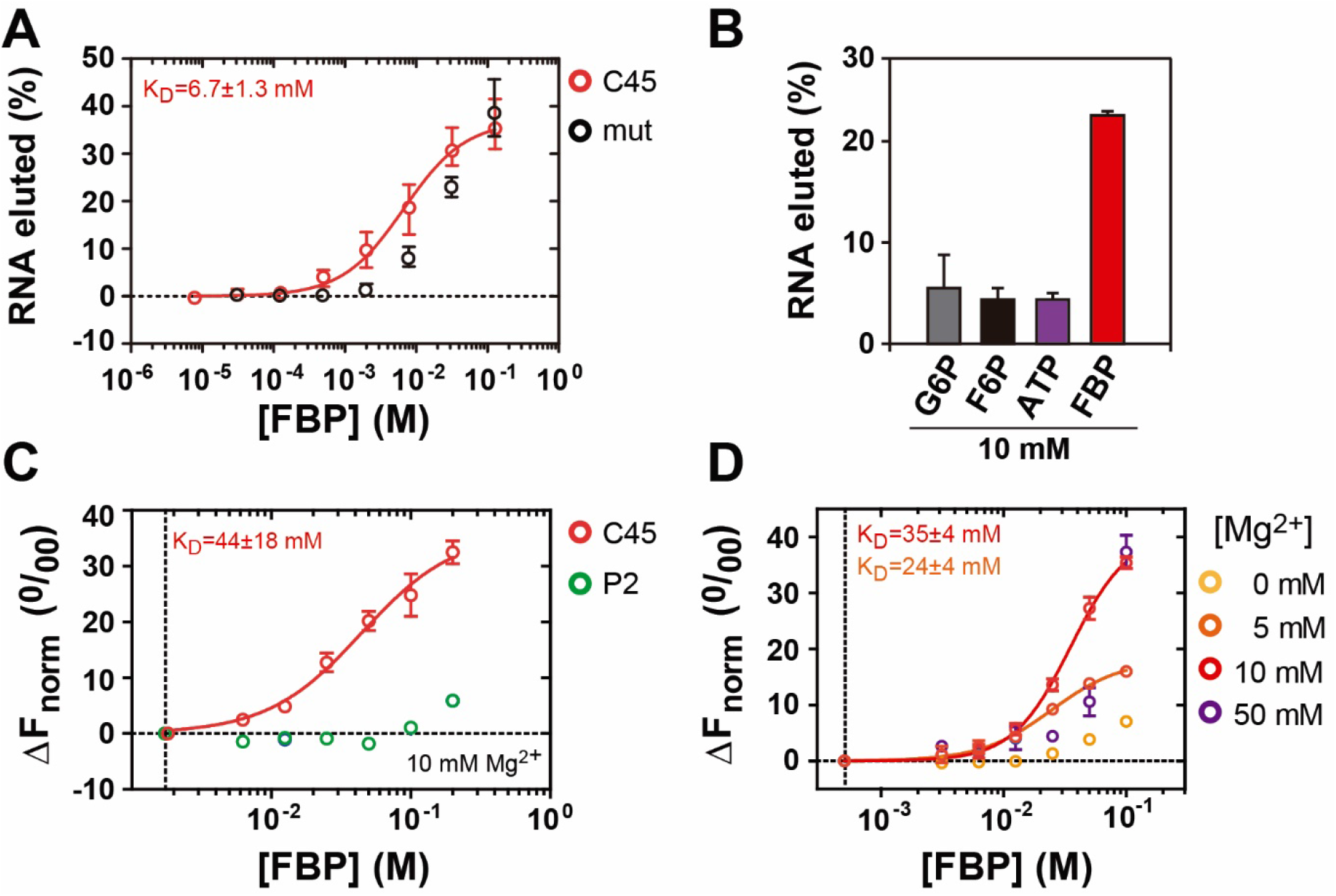
The aptamer C45 binds FBP under physiological conditions. **A**. Binding assays of an RNA clone (C45) retrieved by SELEX and a mutant variant (mut), in which the stem structure predicted to exist in most selected clones is disrupted (Supplementary Fig. S2). Fluorescently labeled (CAL Fluor Red 610) RNAs were incubated with FBP immobilized to sepharose, and eluted with different concentrations of free FBP. **B**. Assay done as in A, but where 10 mM solutions of glucose-6-phosphate (G6P), fructose-6-phosphate (F6P), adenosine-triphosphate (ATP), or fructose-1,6-bisphosphate (FBP) were used to release RNA from a sepharose column with immobilized FBP. **C**. Microscale thermophoresis (MST) analysis of free FBP binding to Cy5-labeled C45 aptamer and a mutant lacking the 5’and 3’-end tails and the four pair of nucleotides that make up the base of the conserved stem (P2) (Supplementary Fig. S2). Binding assays were carried out in cytoplasmic buffer CB (See Methods) with 10 mM MgCl_2_. **D**. MST analysis of FBP-binding to C45 in solution under different MgCl_2_ concentrations. Apparent dissociation constants (K_D_) were estimated using a non-linear regression model assuming one binding-site and specific binding with Hill slope. In D, K_D_ values correspond to binding assays carried out with 10 and 5 mM MgCl_2_, respectively. Results in A-D represent the mean ± one standard deviation from three replicates. Dashed lines in C-D indicate 0 in x and y axes.

So far, we tested FBP binding to the aptamer when this compound was attached to sepharose. To delimit the concentration range of free FBP required for binding to the aptamer, we used microscale thermophoresis (MST). Thermophoresis, the directed movement of a molecule in a temperature gradient, is altered in response to binding-induced changes in size, charge or hydration shell of a labeled molecule, and allows the use of complex buffers and high concentrations of the ligand (41). We tested FBP binding to Cy5-labeled C45 aptamer in the cytosolic buffer used for aptamer selection and found that C45 also bound free FBP in the mM concentration range. In contrast, a trimmed mutant lacking the 5’ and 3’-end tails and four nucleotide-pairs that make up the base of the conserved stem (P2) (Supplementary Fig. 2D) did not show any binding (Fig. 1C). FBP binding was strongly dependent on the presence of Mg^2+^ (Fig. 1D), suggesting that Mg^2+^ is necessary for proper C45 folding and/or for stabilizing the interaction with FBP. Together, these data show that we have identified an aptamer that selectively binds FBP in a Mg^2+^-dependent fashion and within the physiological FBP concentration range.

**Figure 2.**
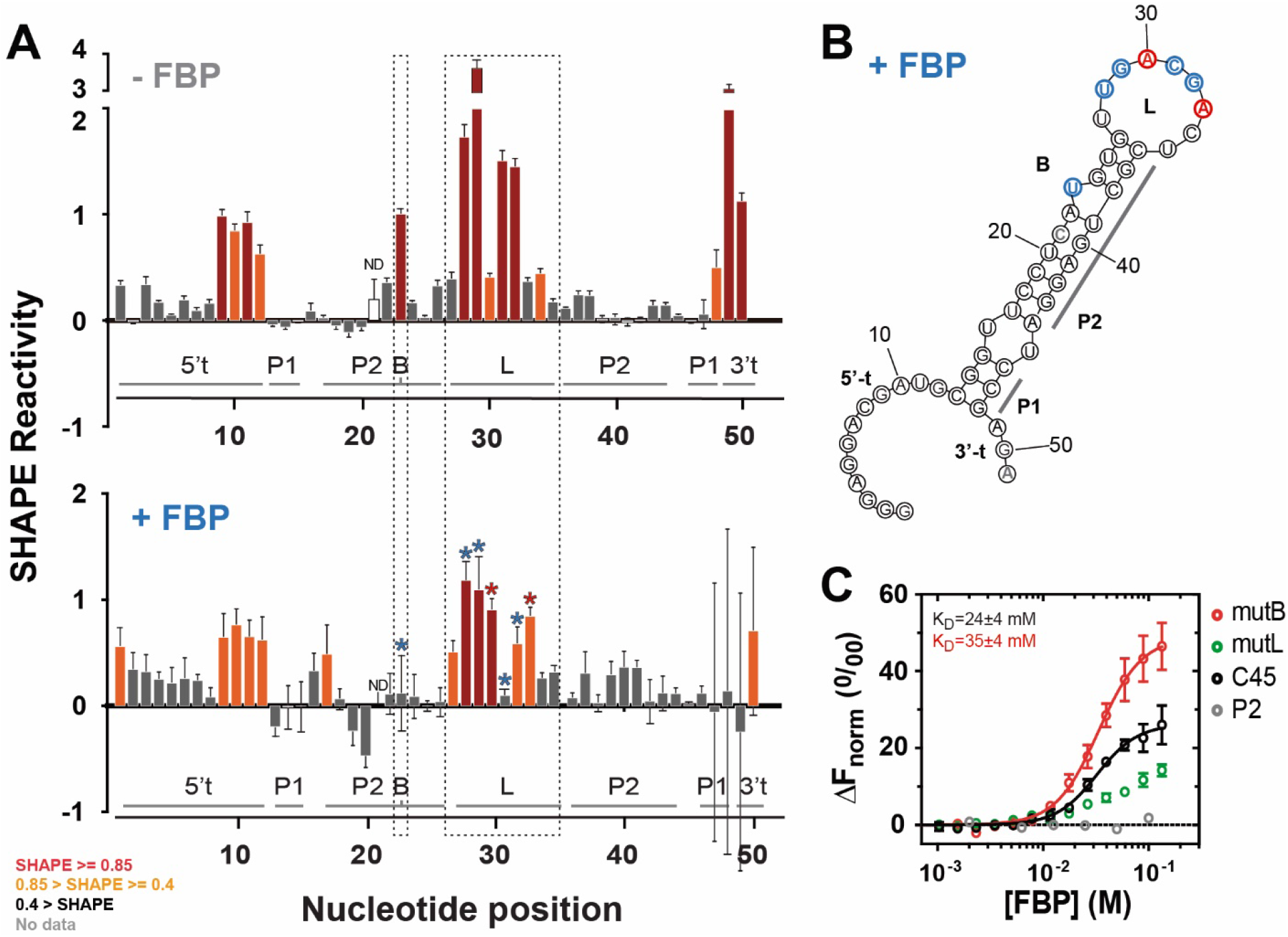
C45 aptamer adopts a stem-loop secondary structure and, upon FBP binding, changes its tertiary structure, involving the loop. **A**. 1-methyl-6-nitroisatoic anhydride (1M6)-SHAPE reactivity profiles of C45 aptamer in the presence (+FBP, bottom panel) or absence (-FBP, top panel) of FBP (86 mM). Bars represent the average of SHAPE reactivities of each nucleotide from five independent experiments; error bars represent the standard error of the mean. Bar colors are used to categorize SHAPE reactivity in three levels: dark grey for low, orange for intermediate and red for high reactivity. SHAPE reactivity values ranges are indicated in the legend (bottom left). Asterisks indicate significant differences, where differences in reactivity were higher than 0.3 and p-values from two-sided student’s t-test lower than 0.05; blue and red asterisks indicate a significant decrease or increase of SHAPE reactivity in the presence of FBP, respectively. Boxes with dotted lines delimit bulge (B) and loop (L) regions in C45 aptamer, where clustered differences in SHAPE reactivity occur. Different parts of the aptamer are indicated below the bars (5’-t and 3’t: tails in 5’ and 3’ ends, respectively; P1, P2: stems 1 and 2; B: bulge; L: loop) **B**. Schematic representation of predicted C45 RNA secondary structure obtained with *RNAstructure* software, which considers energy minimization coupled to SHAPE reactivity. As in A, blue and red indicate nucleotides showing a significant decrease or increase of SHAPE reactivity in the presence of FBP **C**. Microscale thermophoresis (MST) analysis of FBP binding to mutants targeting the regions that altered SHAPE reactivity upon FBP binding: the U-bulge (mutB: U23 deletion) and C45 loop (mutL: U27 to U34 substitution by its complementary sequence). C45 and mutant P2 (deletion of the base of the conserved stem: G1-U16 and U45-A51) show the extremes of the affinity to FBP. Points represent the mean ± S.D. of three independent experiments.

### FBP-binding alters the tertiary structure of C45 RNA aptamer

Towards unraveling the structural changes occurring in the C45 aptamer upon FBP binding, we performed 2’-hydroxyl acylation analyzed by primer extension (SHAPE) (42) analyses, which, coupled with the *RNAstructure* energy minimization algorithm (43), allowed us to generate an evidence-based model of the C45 secondary structure. The SHAPE-reactivity patterns of the C45 nucleotides revealed a highly reactive core flanked by non-reactive regions predicted to fold as a stem-loop (Fig. 2A, top panel), which matched the Mfold prediction solely based on energy minimization (Supplementary Fig. S2A). Specifically, the stem has twelve base pairs (C13-G26 and C36-G48) interrupted by a single mismatch (U16, U45) and a base bulge (U23); and the distal loop (L) contains nine nucleotides (from U27 to U35). The stem-loop structure is flanked by two single stranded unstructured tails (Fig. 2A, top panel).

To determine the structural changes upon FBP binding, we performed SHAPE also in the presence of a saturating concentration of FBP (86 mM). We found significant changes in the SHAPE reactivity in six of the nine residues in the distal loop (U28-A33) and in the bulge of the stem (U23) (Fig. 2A, bottom panel). As the *RNAstructure* model of the secondary structure remained the same as without FBP (Fig. 2B), these FBP-induced changes in SHAPE reactivity most likely result from an altered tertiary structure of the aptamer.

To study the role of the distal loop and the bulge of C45 aptamer in FBP binding, we tested FBP binding to the aptamer variants in which these regions were mutated. In particular, we generated a mutant devoid of the bulge (U23, mutB) and a mutant where we substituted eight out of the nine nucleotides of the loop by their complementary counterparts (U27 to U34, mutL). These mutations did not impair the C45 structure as, according to Mfold energy minimization, mutB and mutL overall maintained the stem-loop secondary structure (Supplementary Fig. S2A, F-G), which we had found characteristic for the majority of the selected aptamer clones (Supplementary Fig. S2A-C and results not shown). In MST analyses, we observed that mutL had an impaired binding of FBP (Fig. 2C), whereas the bulge deletion (mutB) only slightly decreased C45 affinity for FBP (Fig. 2C), and preserved selectivity (Supplementary Fig. S3), indicating that the bulge plays a less relevant role in FBP binding. Collectively, these results show that C45 folds as a stem-loop where FBP binding likely triggers tertiary-structure rearrangements of the aptamer involving the loop.

### An *in vivo* system to screen for RNA devices responding to intracellular FBP levels

To develop this *in vitro*-selected aptamer into an RNA device suitable for FBP sensing in yeast cells, we engineered a reporter system that allowed us to screen large RNA-device libraries *in vivo*. To this aim, we had to couple the C45 aptamer with an actuator domain suitable to relay the FBP-triggered changes in the tertiary structure of the aptamer into a quantitative readout. Here, we employed the hammerhead ribozyme (HHRz) as an actuator domain because it has been shown to allow engineering RNA devices with ligand-responsive ribozyme tertiary interactions (35). By integrating a C45-HHRz device into the 3’-UTR of a GFP-encoding mRNA, FBP binding to C45 would allosterically control HHRz self-cleaving activity, and thus GFP mRNA stability and protein expression (Fig. 3A, Supplementary Fig. S4A).

**Figure 3.**
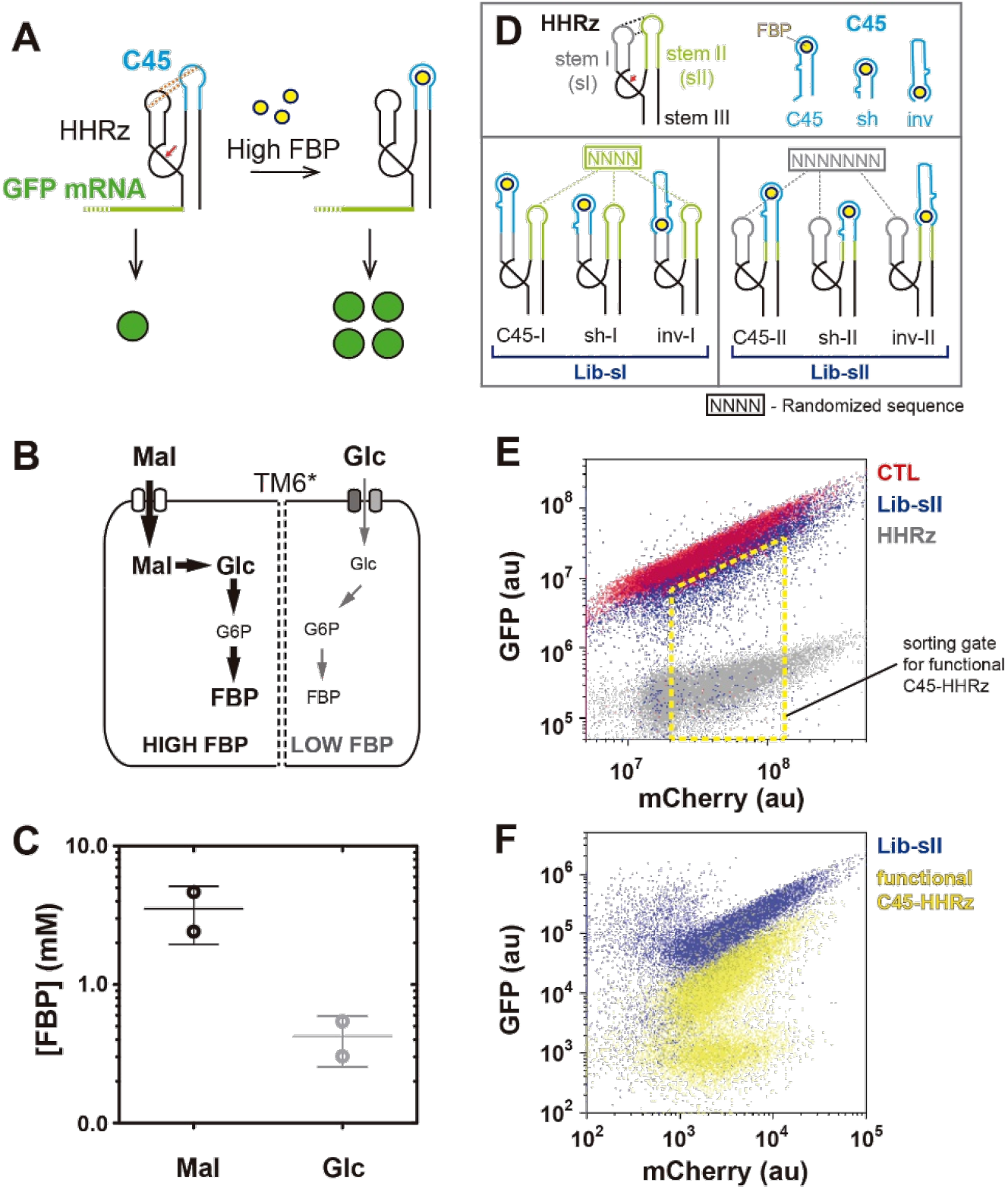
An *in vivo* system to screen for RNA devices that respond to altered intracellular FBP levels. **A**. A reporter system that transduces intracellular FBP concentration into a fluorescent output. C45, the FBP-binding aptamer adopting a stem-loop secondary structure (blue), is grafted into the hammerhead ribozyme (HHRz) from the satellite RNA of tobacco ringspot virus at one of its stem-loops. Long-distance tertiary structure interactions (dotted lines) are necessary to sustain self-cleaving activity of the ribozyme (red arrow). FBP binding at its high intracellular concentration distorts the tertiary structure interactions, and hence inhibits HHRz activity. The C45-HHRz RNA device is cloned in the 3’ UTR of GFP. HHRz-mediated cleavage with low FBP levels leads to a short and unstable GFP mRNA, and thus to low expression of the protein (green circles). In contrast, when FBP levels are high, longer and more stable transcripts lead to higher GFP protein expression. The opposite mechanism of action can be also envisioned when FBP binding activates ribozyme activivty. **B**. A system to modulate intracellular FBP concentrations to set screening conditions. *S. cerevisiae TM6\** strain expresses only a chimeric hexose transporter that sustains a low glucose-uptake rate, and thus a low glycolytic flux (right) and low FBP levels, when glucose is provided (44). Maltose is taken up normally, leading to a high glycolytic flux and thus high FBP level. **C**. Metabolomic analysis of intracellular FBP concentration in *TM6\** yeast cells cultured in minimal medium with either maltose or glucose as the sole carbon source. FBP abundance was obtained from two independent experiments and two determinations per experiment. Cell volume was determined for at least 70 cells per sample. Lines represent the mean of the two experiments ± S.D. **D**. Design of the C45-HHRz RNA device library. C45 aptamer was grafted into either of the two stem-loops of HHRz (stem I and II, grey and green, respectively) using three different strategies: (i) the loop of the receiving stem in HHRz is removed and C45 placed next to the HHRz stem in a direct (C45-I, C45-II) or (ii) inverted orientation (invI, invII), and (iii) the HHRz loop is removed and a shortened version of C45 merged with the HHRz stem (sh-I, sh-II). The sequence of the opposite HHRz loop (that is, the loop I when C45 is grafted into stem II, and vice versa) is randomized (grey and green N-stretches) to generate a library. Out of it, we select those clones that establish interactions between C45 and the opposite HHRz stem loop, and therefore sustain the HHRz activity. Lib-sI and Lib-sII are the libraries made by equimolar pooling of the libraries C45-, sh- and inv, -I and -II. **E**. Sorting of RNA devices with residual HHRz activity in the Lib-sII C45-HHRz library. GFP-mCherry scatter plot where Lib-sII library (blue), along with RNA device controls containing fully active (HHRz, grey) and inactive HHRz (CTL, Red) are overlaid. The sorting gate (yellow dotted polygon) used to collect C45-HHRz RNA devices with functional HHRz is indicated. **F**. Comparison of the initial cellular Lib-sII library and the sorted population enriched in C45-HHRz devices with functional ribozyme. In E and F, 20,000 cells from each population are plotted. Identifiers of the populations plotted are indicated on the right side of each plot. Fluorescence arbitrary units shown in E and F are different because sorting (E) and ulterior analysis after re-growth of the sorted population (F) were done in different devices. For simplicity, sorting and post-growth analysis of sorted population are shown only for Lib-sII.

To correct for cell-to-cell variation in protein expression activity and for other extrinsic factors that may alter the readout of the reporter system, we used a centromeric plasmid, from which we not only expressed the HHRz-GFP construct, but also mCherry without HHRz as an unregulated control (Supplementary Fig. S4B), as done recently (35). Together, this system can report the activity of the HHRz via the mCherry-normalized GFP readout (GFP/mCherry ratio) in single yeast cells. To investigate the reporter system’s dynamic range and resolution, we analyzed the readout of a set of mutants of HHRz (without the FBP aptamer included yet), which were recently shown to deliver different GFP/mCherry values (35) (Supplementary Note 1). We found that GFP/mCherry readout spanned over two logs, which enabled us to separate eight yeast clonal populations with different GFP/mCherry ratios without significant overlap (Supplementary Fig. S4B-D), indicating a broad dynamic range of the reporter system. Thus, we have implemented a reporter system in yeast that quantitatively relays HHRz activity to a GFP/mCherry readout, which is suitable for screening RNA devices with HHRz tertiary structure-dependent activities.

For the screening, we next needed a strategy to modulate intracellular FBP concentrations. To this end, we used the engineered *Saccharomyces cerevisiae* strain *TM6\**, which has only one chimeric hexose transporter (instead of the 17 of the wild type) and thus displays a low glucose uptake rate and a low glycolytic flux (45). In contrast, when grown on maltose, which is transported into the cell with a different transporter, this strain has a high glycolytic flux, just like the wild type (44) (Fig. 3B). Since glycolytic flux generally correlates with FBP levels (10, 46, 47), we hypothesized that growing *TM6\** cells in these two carbon sources could be utilized as the two extreme FBP conditions for screening. Quantitative targeted metabolomics demonstrated that FBP levels are on average about 9-fold lower in *TM6\** grown on glucose compared to maltose (Fig. 3C), reflecting a spread which nearly covers the physiological dynamic range of FBP (9, 46). Thus, by growing the *TM6\** strain on two different carbon sources, we could control FBP levels within its physiological range, and therefore we could use this system to screen *in vivo* for RNA devices responding to physiological FBP concentration changes.

### Design and pre-selection of C45-HHRz RNA devices with a functional ribozyme

To identify RNA devices with an FBP-dependent ribozyme activity, we next constructed the libraries of HHRz-C45 RNA devices where we grafted the C45 aptamer into one of the two HHRz stem-loops (I or II) involved in tertiary interactions required for self-cleavage (35). FBP-binding would then disrupt tertiary interactions between the C45 aptamer’s loop and the opposite stem-loop of the ribozyme. Here, either the HHRz stem-loop I could receive the aptamer to interact with the stem-loop II, or *vice versa* (Fig. 3A). However, grafting the aptamer into an HHRz stem-loop itself may distort HHRz structure, thereby rendering the ribozyme inactive irrespective of FBP binding.

To increase the chances of maintaining residual ribozyme activity after aptamer integration and to mitigate a potential structural bias of using a single grafting strategy, we introduced the FBP aptamer into either the stem-loop I (Supplementary Fig. S5A-C) or II (Supplementary Fig. S5D-F), each with three different strategies (Fig. 3D). In the first strategy, we removed the HHRz loop and appended C45 to the remainder of the stem (the receiving stem) in the same orientation (*C45*, Supplementary Fig. S5A, D). In the second strategy, we did the same but just inserted C45 in the inverted orientation (*inv*, Supplementary Fig. S5C, F). In the third strategy, we replaced the whole HHRz stem-loop by a version of the FBP aptamer containing a shortened stem (*sh*, Supplementary Fig. S5B, E). With these three integration strategies, we generated combinatorial libraries of the RNA devices in which the HHRz loop sequence of the stem-loop opposite to the receiving stem was randomized: four nucleotides of loop II when the receiving stem was stem I, and seven nucleotides of loop I when the receiving was stem II (Fig. 3D, Supplementary Fig. S5). The libraries generated with these three different C45-integration strategies in each of the two stems I and II were pooled and transformed into yeast, resulting in two C45-HHRz RNA device libraries (Lib-sI and Lib-sII, Fig. 3D) with theoretical sizes of 768 and 49,152, respectively.

From these two C45-HHRz libraries we next selected RNA devices in which the ribozyme was still functional after C45 insertion as only those with residual ribozyme activity are suitable for later screening for FBP-dependent regulation of that activity. To this aim, we used FACS to collect those clones in the libraries with low GFP expression at the low FBP condition (*i*.*e*. in glucose-grown *TM6\** cells) (Fig. 3E) because then RNA devices should be mainly in their FBP-unbound state, and thus with unperturbed basal HHRz activity. We defined the reference points for non-self-cleaving activity and maximal self-cleaving activity with two of the HHRz-GFP constructs used for the calibration of the reporter system (without C45): the inactive HHRz (CTL, Supplementary Fig. S4B-D) and wild-type ribozyme (HHRz, Supplementary Fig. S4B-D). Here, we found that more than 95% of the clones in the C45-HHRz libraries had a non-functional HHRz, with GFP/mCherry ratio overlapping with that of the CTL inactive mutant (Fig. 3E). Then, we sorted the cells that had GFP/mCherry ratio lower than in the CTL mutant, thus collecting RNA devices with a functional ribozyme (Fig. 3E). By preselecting C45-HHRz RNA devices with a residual ribozyme activity we enriched the library in clones suitable for screening, thereby enhancing the chances to identify RNA devices with FBP-dependent regulation.

Next, we wondered which of the three aptamer grafting strategies and which sequence features in the randomized loop would preserve HHRz activity. Through next-generation sequencing (NGS) of the pre-selected populations, we found that active C45-HHRz devices predominantly contained the inverted (*inv*) aptamer in either of the HHRz stems, and the shortened aptamer replacing the whole HHRz stem-loop I (*sh*) (Supplementary Fig. S6D). The prevalence of *sh* variants in stem I within the active devices could be connected with the fact that stem I is naturally longer compared to stem II and therefore, its replacement by the aptamer having a comparable length might have less impact on the ribozyme structure (compare Supplementary Fig. S5B and E). Considering also that in *inv* variants the length of the receiving stem is not altered (Supplementary Fig. S5C, F), these results suggest that the length of the stem into which the aptamer is grafted is a critical factor for ribozyme functionality.

Finally, the analysis of the sequences of the HHRz loop opposite to the aptamer-receiving stem in the active HHRz variants revealed an enrichment of A and G in stem-loop II, and a high conservation of A in the fourth position when the shortened C45 was inserted in place of stem-loop I (*sh-I*) (Supplementary Fig. S6D). On the contrary, when C45 was inserted in place of stem-loop II, we found an overrepresentation of G and T in the randomized region of the stem-loop I. Interestingly, C was observed underrepresented in the randomized region of either stem-loop (Supplementary Fig. S6D). These features preserving ribozyme activity could be utilized in the generation of new, doped C45-HHRz libraries with a higher fraction of functional ribozymes to increase screening effectiveness.

At this point, we have generated an RNA device library with multiple FBP aptamer integration strategies from which we collected the variants with residual ribozyme activity, which is a requirement for the next step, *i*.*e*. to identify the variants where FBP levels influence this activity.

### High-throughput *in vivo* screening of RNA devices that report intracellular FBP

To identify C45-HHRz clones with an FBP-dependent readout, we implemented a high-throughput approach that examines the mCherry-normalized GFP expression of thousands of C45-HHRz variants simultaneously in a single experiment, previously referred to as FACS-Seq (35). Globally, our approach entailed four steps. First, we pooled the two libraries containing the functional C45-HHRz devices (c.f. Fig. 3F) into a single population (*input* from now on), which was then grown in either the high or low intracellular FBP condition, *i*.*e. TM6\** cells cultivated in either maltose or glucose, respectively (Fig. 4A, *growth*). Second, we sorted the cells of the input library into six sub-populations according to the cells’ GFP/mCherry expression ratios (Fig. 4A, *cell sorting*). Third, after identifying all clones in each sorted sub-population by NGS and inspecting each clone’s frequencies, we assigned a single GFP/mCherry ratio value (*µ*_*i*_ hereafter) to every C45-HHRz variant when grown in either condition (Fig. 4A, *NGS*). Finally, analyzing the distributions of *µ*_*i*_, we identified the clones that respond to the difference between the high and low intracellular FBP conditions.

**Figure 4.**
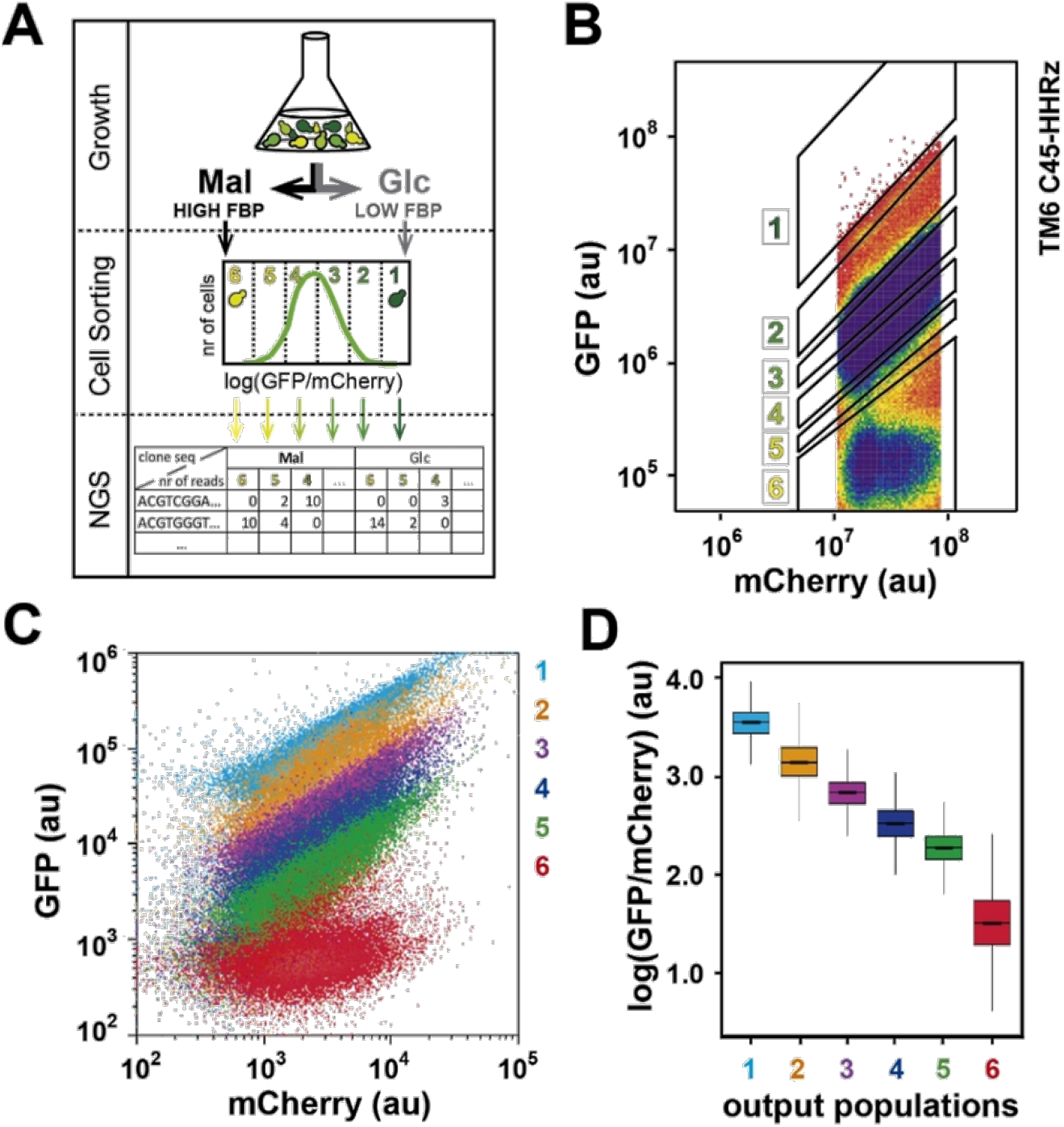
Sorting of C45-HHRz RNA device library yields six sub-populations with distinct GFP/mCherry ratios. **A**. Experimental pipeline of the high throughput *in vivo* screening of C45-HHRz RNA devices that respond to FBP. *TM6\** cells with the C45-HHRz library are grown in medium with either glucose or maltose (top panel). Cells are sorted into six sub-populations according to their GFP/mCherry ratios (middle panel). Sorted cells sub-populations are subjected to next-generation sequencing to identify the distribution of each clone throughout the population. **B**. Sorting of C45-HHRz RNA device library. A representative example of the gating of a FACS-seq sorting experiment is shown (C45-HHRz library grown in glucose; 378,309 cells plotted). Cells with an intermediate 1-log-wide mCherry expression were gated. We used this gating because we found that the GFP signal was less robust in cells with low mCherry expression in clonal populations (*c*.*f*. GFP-mCherry signal overlap among different HHRz versions for mCherry values between 5×10^5^-10^7^ in Supplementary Fig. S7A), while in cells with intermediate mCherry levels dispersion of GFP values remained constant across the bins (Supplementary Fig. S7B). Sorting gates were established according to GFP/mCherry ratio, ranging from the signal of fully active (gate 6) to that of inactive ribozymes (gate 1). The width of the gates 2-5 was set according to the width of a cell population from a single clone in GFP/mCherry plot. **C**. Overlay of GFP-mCherry scatter plots obtained from regrown, sorted sub-populations. Sorted sub-populations were grown for several generations prior to analysis. 20,000 cells from each sub-population are plotted. **D**. Single-cell GFP/mCherry ratio distribution in sorted sub-populations from C45-HHRz library calculated from experiment shown in C. The boxes show the interquartile range with the median in the middle, and the whiskers spanning from the 1^st^ to 99^th^ percentile.

To sort the C45-HHRz variants, we defined six bins on the basis of GFP/mCherry ratio. To this end, we first set the upper and lower limits of GFP/mCherry ratio to the readouts of the CTL inactive mutant (maximal GFP/mCherry ratio) and of the HHRz wild-type sequence (minimal ratio) (Supplementary Fig. S7A), thereby defining the boundaries of the readout’s dynamic range. Then, we split this range into six non-overlapping bins whose width was set to the size of the dispersion of GFP and mCherry signals (or intrinsic noise) in clonal populations (Fig. 4B, Supplementary Fig. S7A). With these settings, for both the low and the high FBP conditions, the sorting yielded six sub-populations with barely overlapping GFP/mCherry expression profiles that were stably maintained over generations (Fig. 4C). The single-cell GFP/mCherry distribution of all sub-populations evenly covered the readout’s dynamic range (*i*.*e*. without gaps with under-represented values or with peaks of over-represented values) spanning more than three logs (Fig. 4D). These results demonstrated that we could sort the clones of our input library into sub-populations with distinct and stable GFP/mCherry values.

Next, we used NGS to identify the C45-HHRz variants and to determine their respective frequencies in each sub-population. We utilized the variants’ frequencies in every sub-population together with the fraction of the input population that was sorted to each bin in the sorting experiment to reconstruct each variant’s distribution across the bins (Supplementary Table S2). The sum of these reconstructed absolute frequencies across the bins for every variant significantly correlated with the variants’ frequencies in the unsorted input population, indicating that our sampling method, based on GFP/mCherry-ratio-guided population binning, produced a fair representation of the entire population (Supplementary Fig. S8A). To determine µ_i_ of each variant, the medians of cells’ GFP/mCherry ratio in the bins (determined in each sorting experiment) were weighted by the distribution of the variant across the bins (Supplementary Table S2). The robustness of determining µ_i_ values was demonstrated by a strong correlation between two replicate experiments (Supplementary Fig. S8B). Through this procedure, we could assign a µ_i_ value for each intracellular FBP condition to 15,031 unique C45-HHRz variants (covered by four or more reads in two independent sorting experiments, Supplementary Notes 2-3).

To identify clones with significantly different *µ*_*i*_ values between the conditions of low and high intracellular FBP, we defined the ‘response’ of the sensor, *D*_*i*_, a new variable that corresponds to the logarithm of the quotient between the *µ*_*i*_ of a clone in the two FBP condition: 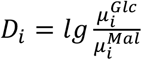 (Supplementary Table S2). The analysis of the distribution of this response variable revealed 63 clones (0.42% of all screened clones) that consistently showed an extremely high or low *D*_*i*_ value (*i*.*e*. outliers) in two replicate experiments (Fig. 5A, Supplementary Note 3). Thus, these 63 clones showed a condition-dependent change in *µ*_*i*_ indicative of an FBP-dependent response.

**Figure 5.**
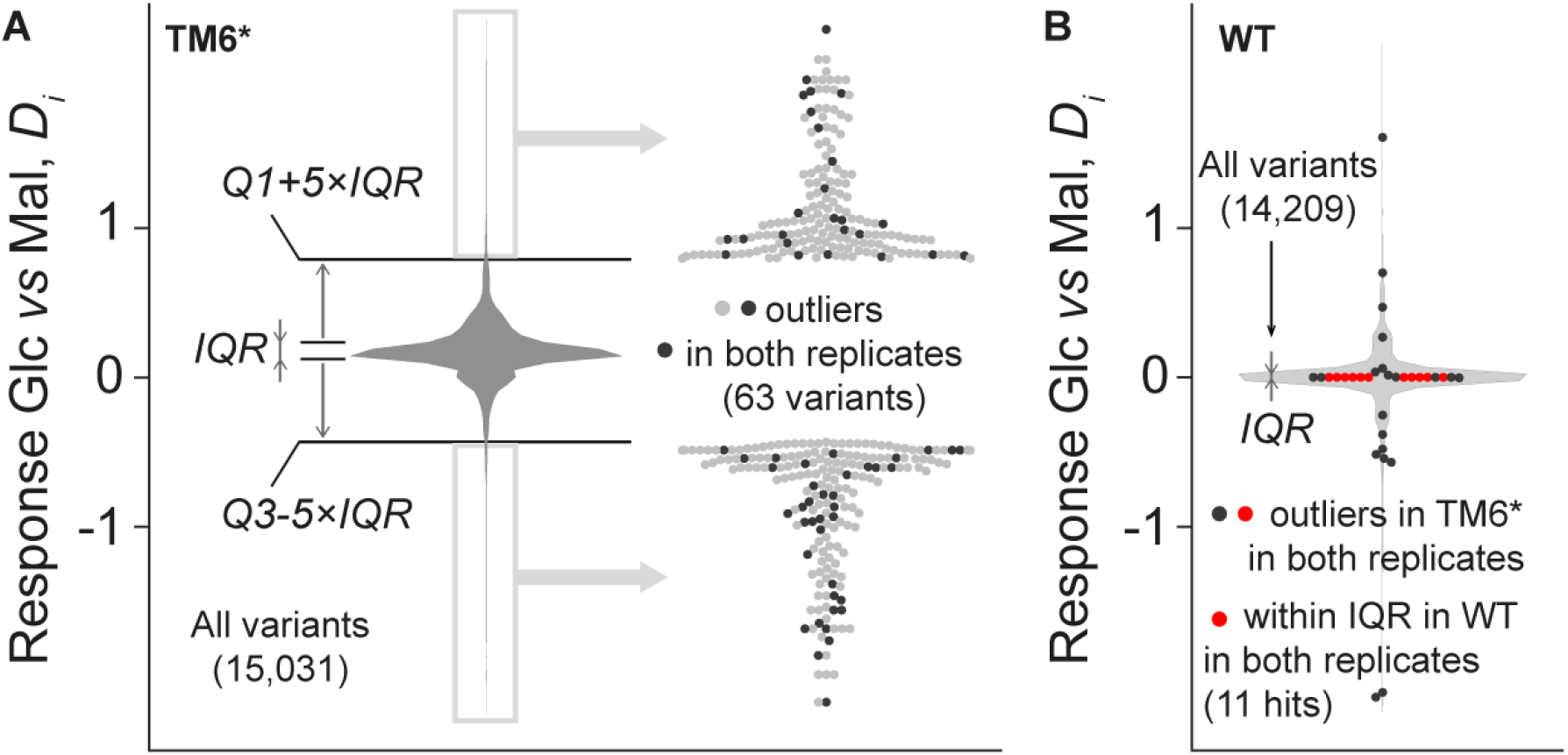
Hit selection in high-throughput *in vivo* screening of C45-HHRz variants that respond to differences of intracellular FBP. **A**. Identification of variants in C45-HHRz library expressed in the *TM6\** strain that have different response while growing on glucose versus growing on maltose, associated with low and high intracellular FBP, respectively. To compare a variant’s response to the change in intracellular FBP, we calculate the measure Response Glc *vs* Mal, *Di* (y-axis; see Table S2 for details), a transformed measure of the change in the GFP/mCherry ratio 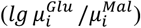, and check if its value deviates from those of other variants. The distribution of this response *Di* of all the variants is depicted in the violin plot. We select variants with the extreme response values that are located beyond the segment [*Q*1 – 5 · *IQR, Q*3 + 5 · *IQR*] (grey boxes on the left, grey and black dots on the right), where *Q*1 and *Q*3 are the lower and upper quartiles, and *IQR* is the interquartile range. To exclude variants that end up in the selection due to technical noise, we repeated the sorting experiment and looked for clones with consistently deviating response (63 clones, black dots on the right). All the clones in this analysis are covered by 4 sequencing reads or more in each of the four sorting experiments (2 carbon sources × 2 independent experiments). **B**. Exclusion of variants showing an intracellular FBP-independent response. Wild-type (WT) cells display a similar FBP level when cultured in glucose or in maltose, and thus, hits should not show a response between these carbon sources in C45-HHRz library expressed in wild-type background. 11 variants are located within the *IQR* of the response in C45-HHRz library in the wild type in two independent sorting experiments, these variants are thus considered the hits from the screening. The presented distribution and its *IQR* used for selection is calculated from the data of variants supported by 4 sequencing reads or more in each of the four sorting experiments; the values corresponding to the dots were obtained using the data supported by at least 2 reads in each of the four sorting experiments. For simplicity of the representation, in both **A** and **B**, we present the data corresponding to one replicate experiment (R1). See Supplementary Note 3 describing the hit selection in more detail.

In the screening experiments, we perturbed the intracellular FBP levels by changing the carbon source on which *TM6\** cells grew (low FBP on glucose and high FBP on maltose, Fig. 3C). To sort out clones that might respond to unknown carbon-source-related changes aside from FBP level differences, we next screened the C45-HHRz library also in wild-type cells. In contrast with *TM6\**, wild-type cells grown in glucose and maltose have nearly identical glycolytic rates, and thus FBP levels (9). Therefore, an ideal FBP-responding candidate should have different µ_i_ values in glucose and maltose in *TM6\**, but identical values in the wild type (where on both carbon sources FBP levels are high). Here, we found that out of the 63 strongly responding clones identified with *TM6\** (Fig. 5A) eleven had the response *D*_*i*_ around 0 in the wild type (Fig. 5B). Thus, our *in vivo* screening led to the discovery of eleven hits showing FBP-dependent changes in the GFP/mCherry ratio readout.

### *In vivo* FBP binding is essential for sensor functionality

To validate the hits as intracellular FBP sensors, we expressed them in the *TM6\** and wild-type strains, and analyzed their GFP/mCherry ratios in clonal populations grown on either maltose or glucose (which lead to different intracellular FBP concentrations in *TM6\** but not in the wild type). We found that nine out of ten tested hits had a different readout between cultivations on the two carbon sources in *TM6\** while they did not in the wild type (Fig. 6A), indicating that these hits exhibit an FBP-dependent response. A random variant not identified as a hit in the high-throughput screening, carried along here as a control, responded neither in *TM6*\* nor in the wild type (Control 1_10, Fig. 6A). These experiments validated the FBP-dependent response of nine of the hits identified in the screening.

**Figure 6.**
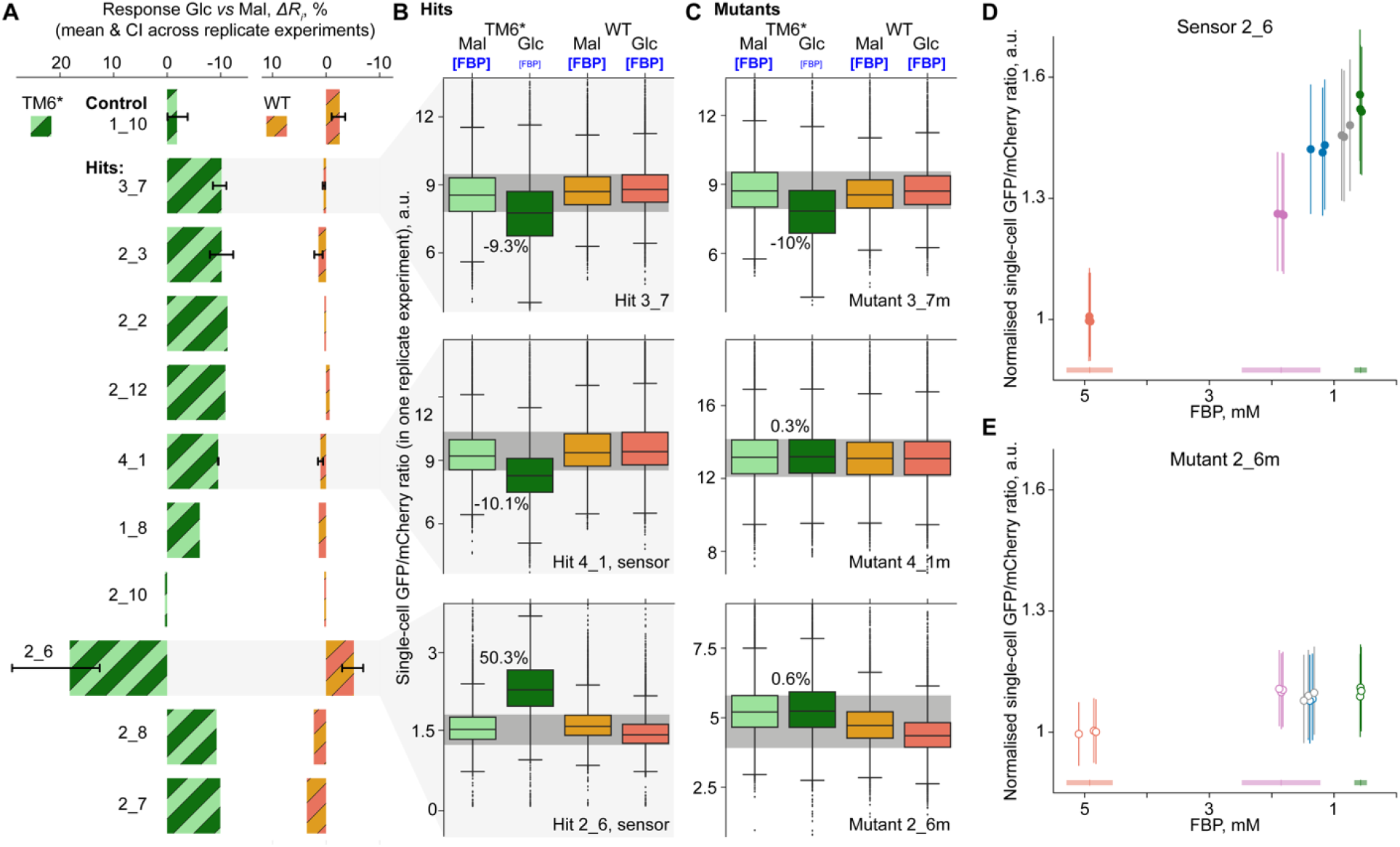
Validation of individual hits as FBP sensor candidates. **A**. Response of hits in cells cultivated in glucose as compared to cells cultivated in maltose (*Response Glc vs Mal, ΔRi*). Intracellular FBP concentration differs in *TM6\** strain and stays the same in the wild-type (WT) strain, when cells are grown on these two carbon sources (Glc, glucose; Mal, maltose). *Response Glc vs Mal, ΔRi* is calculated as the percentage of the change in the GFP/mCherry ratio between the cells cultivated in glucose and maltose as compared to the cells cultivated in maltose: 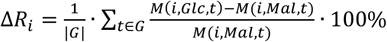, where *i* is a hit, *M*(*i, Glc,t*) and *M*(*i, Mal, t*) are the median GFP/mCherry ratios in the population of cells grown on glucose and maltose, respectively; *G* is the set of three (|*G*|) time points *t* in the exponential part of a culture’s growth curve. The bar and whisker lengths denote the mean value of Δ*Ri* and its 95%-confidence interval calculated for the replicate cultivation experiments. No whiskers are shown in case of single cultivation experiment. The following numbers of data points were used to calculate the mean and the confidence interval in both TM6* and WT: 6 for 1_10 in both strains, 3 for 3_7, 2 for 2_3, 3 for 4_1 and 3 for 2_6. **B-C**. The single-cell distribution of the GFP/mCherry ratio in the four populations (2 strains × 2 carbon sources) at one time point *t* in the growth curves of selected hits (B) and their mutants (C). The box shows the interquartile range *IQR*, the line in the center of it denotes the median *M*(*i, Mal, t*) or *M*(*i, Glc, t*), the whiskers spread until *Q*1 – 1.5 × *IQR* or *Q*3 + 1.5 × *IQR*, where *Q*1 and *Q*3 are the lower and upper quartiles. The percentage between the distributions of maltose- and glucose-grown TM6* populations is the value of the ratio 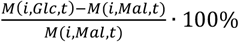, which contributed to the calculation of the response variable *ΔR*_*i*_ in case of hits (A). The dark-gray horizontal band denotes the central tendency of the values of the GFP/mCherry ratio associated with high intracellular FBP, with the upper and lower boundaries being the maximal upper quartile and the minimal lower quartile across the populations of TM6* on maltose, WT on maltose and glucose. To build each of the boxplots, from 15,786 to 18,729 single-cell values were used. **D-E**. The dependence of the GFP/mCherry ratio of the sensor 2_6 (D) and its mutant 2_6m with impaired FBP binding (E) on the intracellular FBP concentration varying across strains with different glucose uptake rates and glycolytic fluxes(45). Five strains shown in different colors (from left to right: wild type, *HXT7, TM3, TM4* and *TM6\**) were cultivated on glucose in three replicate batch cultures. For every replicate culture, the median and interquartile range (IQR) of single-cell GFP/mCherry ratios are presented as a circular marker and thin vertical line, respectively. To normalize the sensor and mutant readouts, we divided single-cell GFP/mCherry ratios by the average of the medians in three replicate wild-type cultures (1.44 for the sensor and 4.54 for the mutant). To calculate the medians and IQRs, we used from 23,474 to 86,554 cells analyzed in flow cytometry. The intracellular FBP concentrations of the wild-type, *HXT7* and *TM6*\* strains as previously measured in (48) are summarized via the mean ± standard deviation of replicate cultivations and shown as tick horizontal lines at the bottom of the plots. The estimation of the FBP concentration in the replicates and strains yielding GFP/mCherry ratios is described in the *Methods* section.

To confirm the specificity in the observed FBP-dependent response, we mutated the C45 aptamer in some validated hits. Drawing on our finding that FBP binding is impaired *in vitro* when the aptamer’s loop is mutated (Fig. 2C, *mutL*), we generated *mutL* versions of C45 in three validated hits. For hit 3_7, we found that the mutated and non-mutated versions showed a similar response between the carbon sources in *TM6\** (Fig. 6B-C), disproving that FBP binding triggers the observed change in the GFP/mCherry ratio. However, the *mutL* versions of the other tested hits (4_1m and 2_6m) lost the FBP-dependent change of the readout (Fig. 6B-C), demonstrating that in 4_1 and 2_6 FBP binding to C45 is essential for the observed response, confirming that these are FBP sensors.

Focusing on the developed sensor 2_6 with the biggest amplitude of the readout (Fig. 6A-B, 2_6), we investigated the sensor’s response to different intracellular FBP concentrations between the extreme levels employed in the high-throughput screening. For this, we exploited the *HXT7, TM3* and *TM4* strains which were previously shown to have different glucose uptake rates and glycolytic fluxes located between the respective values of the wild-type and *TM6\** strains used so far (45). Here, we found that, across the five strains, the GFP/mCherry ratios of the sensor were markedly different and negatively correlated with intracellular FBP concentrations (Figure 6D). On the contrary, the GFP/mCherry ratios from the mutated sensor with impaired FBP binding did not markedly change with different intracellular FBP concentrations (Figure 6E). Thus, the sensor 2_6 displays a dynamic range that covers various FBP concentrations observed *in vivo*.

We next wondered about the RNA structural elements relevant for sensor functionality *in vivo*. To this end, we performed *in vivo* RNA structural probing in which cells expressing the sensor 2_6 were treated with dimethyl sulfate (DMS) followed by targeted DMS-MaPseq analysis (49). Here, we found that the DMS-reactivity pattern is consistent with the expected formation of stem-loop I and a long stem with an internal loop, which is a product of merging HHRz stem II and the aptamer C45 (Figure 7A-B; Supplementary Fig. S5F). Zooming in on the aptamer region in the RNA device, we further found that, although some nucleotides involved in the loop *in vivo* (Figure 7B, U39-C42, U64-U66) are *in vitro* related to the stem (Figure 2B, G37-C38, U23-U25), the biggest part of the grafted aptamer (Figure 7B, U43-A63) has the same stem structure *in vivo* as determined *in vitro* (Figure 2B, C13-A22, U39-G48). In particular, we observed that the P1 sequence and the WUCCU motif (Figure 7B, U57-U61) constitute the stem also *in vivo*, identically to most *in vitro*-selected FBP-binding aptamer clones (Supplementary Fig. S2A-C; results not shown). Supporting the importance of C45 aptamer folding into this stem observed both *in vitro* and *in vivo*, DMS-structural probing of the FBP-unresponsive mutant 2_6m (Figure 6C) revealed a strand displacement that dramatically disrupts the folding of the merged stem II and the aptamer (Figure 7A-B). Thus, the structural elements identified *in vitro* to be critical for FBP binding by C45 aptamer are also essential for the FBP-dependent response of the sensor *in vivo*.

**Figure 7.**
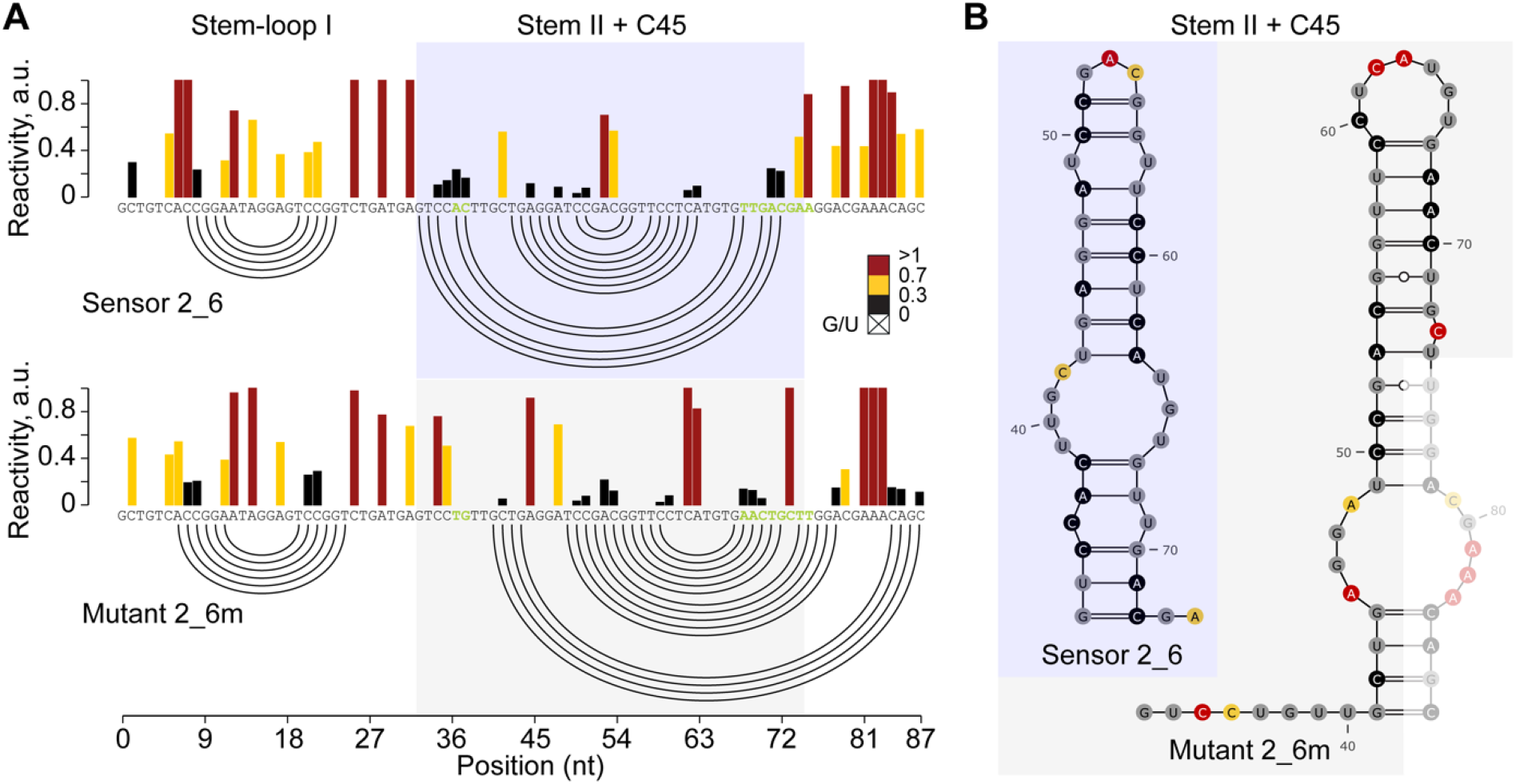
The intracellular-FBP sensor *in vivo* maintains the structural elements of the C45 aptamer that are critical for FBP binding *in vitro*. **A**. *In vivo* DMS-MaPseq analysis of the C45-HHRz RNA device in the sensor 2_6 and its mutant 2_6m. Bases differing between the two sequences are highlighted in green. Bars are color-coded according to base reactivity (low: black, medium: yellow, high: red). Mutational data has been normalized by box-plot normalization. Reactivities were capped at 1 for representation purposes. Experimentally-constrained secondary structures are depicted as arc plots. **B**. Secondary structure models of Stem II merged with the C45 aptamer (Stem II + C45) in the sensor 2_6, and of the corresponding region in the mutant 2_6m. Bases are colored according to reactivities shown in A.

Finally, we wanted to get evidence that the FBP-dependent response of the sensor relies on its riboswitch activity. We hypothesized that if FBP binding results in a conformational change of the RNA device, our sensor should occur *in vivo* as an equilibrium of two conformations: FBP-free and FBP-bound. To test if there are indeed two conformations of the sensor *in vivo*, we further analyzed DMS-MaPseq data. Compared to traditional DMS probing, DMS-MaPseq has also the advantage of recording multiple bases that were simultaneously single-stranded as mutations within the same cDNA molecule (50) (Morandi *et al*., manuscript in preparation). By means of spectral clustering, this information can be exploited to identify multiple co-existing RNA conformations (50) (Morandi *et al*., manuscript in preparation). Here, spectral clustering analysis showed that the sensor 2_6 exists in two conformations within the cell (Supplementary Fig. S9, 2_6), likely corresponding to FBP-free and FBP-bound states. Consistent with the premise that these two conformations correspond to the FBP-free and bound forms of the sensor, for the mutant 2_6m, which lacks FBP binding, we only observed one conformation *in vivo* (Supplementary Fig. S9, 2_6m). These results support the notion that an interaction of FBP with the sensor 2_6 triggers a conformational change necessary for the FBP-dependent response of the sensor.

### A developed FBP sensor functions as a glycolytic flux biosensor

Next, we wanted to apply our sensor to study metabolic heterogeneity. Since intracellular FBP concentrations were shown to correlate with the flux through glycolysis (10, 46, 47), we hypothesized that our developed FBP sensor would be suitable for reporting glycolytic flux in single cells. Non-dividing cells, which can be present in normal yeast cultures, were shown to exhibit a reduced glucose uptake rate, and thus glycolytic flux, compared to the co-existing dividing cells (44). To examine if the 2_6 FBP sensor can identify the different metabolic fluxes of dividing and non-dividing cells within the same yeast population, we cultured *TM6\** cells with the sensor on glucose in a microfluidic device (51, 52) and monitored the GFP/mCherry ratio in both subpopulations. Here, we found a markedly higher GFP/mCherry ratio in the non-dividing cells compared to the dividing cells (Fig. 8A), in line with the low glycolytic flux in the non-dividing cells (44). The same strain grown on maltose showed a ratio lower than that of cells dividing in glucose (Fig. 8A), consistent with the higher FBP levels on this carbon source (Fig. 3C), and the increased glycolytic flux (44). Across these three groups of cells, we could draw a notable correlation between the single-cell growth rate and the GFP/mCherry ratio (Fig. 8A). In contrast, the *mutL* version of the sensor, which has impaired FBP binding (Fig. 2C and 6C, mutant 2_6m), did not show any pronounced difference in the readout among the three groups of cells (Fig. 8B).

**Figure 8.**
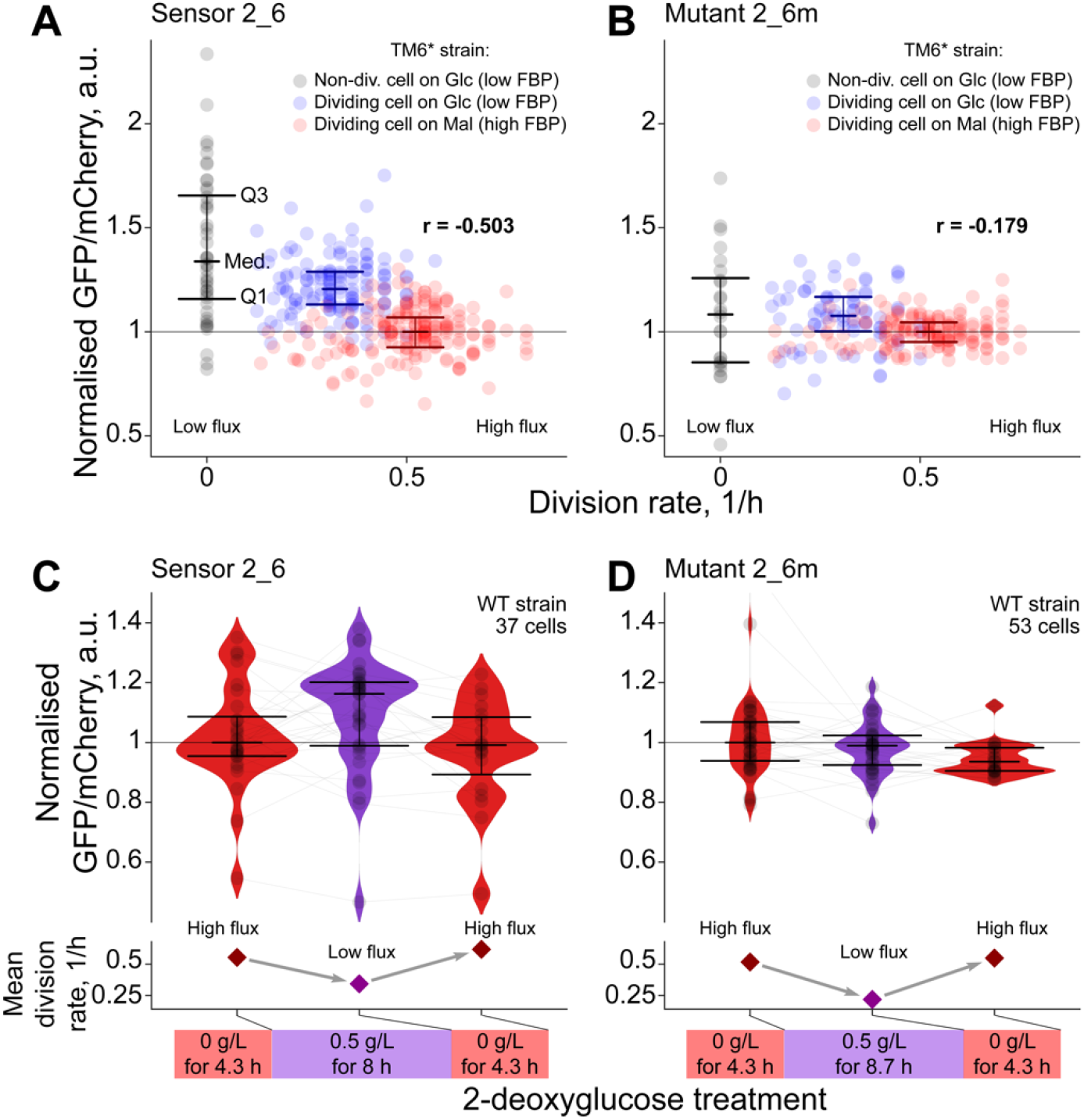
The FBP sensor functions as a glycolytic flux biosensor. **A-B**. The GFP/mCherry ratio of the sensor 2_6 (A) or its mutant 2_6m (B) in three categories of cells, namely, non-dividing TM6* cells consuming glucose (44 in 2_6 and 24 in 2_6m), dividing TM6* cells consuming glucose (138 in 2_6 and 77 in 2_6m) and dividing TM6* cells consuming maltose (183 in 2_6 and 151 in 2_6m). In case of the dividing cells, a data unit is the mean GFP/mCherry ratio throughout one cell cycle (from budding to budding). For the category of the non-dividing cells, we calculate the mean GFP/mCherry ratio in a single-cell trajectory between 15 and 20 hours after the last division. Q1, Q3 and Med. denote the lower and upper quartiles, and the median. The sensor’s and mutant’s readouts are normalized by the respective median GFP/mCherry ratio values of dividing TM6* cells consuming maltose (horizontal line).The division rate is the reciprocal for the cell-cycle duration in case of the dividing cells and zero in case of the non-dividing cells. We assess the correlation using the Pearson’s coefficient *r*. **C-D**. The GFP/mCherry ratio of the sensor 2_6 (C) or its mutant 2_6m (D) in single wild-type (WT) cells cultivated on 2 % (20 g/L) glucose and dynamically exposed to 0.5 g/L 2-deoxyglucose for several hours. The black markers denote the single-cell GFP/mCherry-ratio values averaged over 100 minutes before the indicated moments of the dynamic experiment. We present the readout of a cell after a particular phase of the treatment (indicated on the X-axis) only if this cell was present in the microfluidic device during the whole phase. Cells present in the microfluidic device in multiple phases of the treatment are indicated with gray lines connecting black markers. Specifically, in both C and D, 14 and 31 cells were present in three and two consecutive phases respectively, whereas the rest of the cells were present only during one full phase. The distributions of the readout are summarized via the violin plots (kernel bandwidth 0.2) as well as with the help of the lower and upper quartiles, and the median (like in A). The sensor’s and mutant’s readouts are normalized by the respective median GFP/mCherry-ratio values of the cells in the first phase of the treatment, that is without the inhibitor (horizontal line). The mean division rate is calculated as 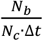, where *N*_*c*_ and *N*_*b*_ are the total numbers of cells and buddings observed during the time Δ*t* hours of a particular phase of the treatment. Cellular-autofluorescence correction and smoothing of the single-cell traces are described in the *Methods* section.

To further test that our FBP sensor responds to changes in glycolytic flux dynamically, we perturbed the metabolism of wild-type cells with 2-deoxyglucose (2-DG), an inhibitor of glycolysis. Specifically, we monitored the readout of the sensor 2_6 in wild-type cells cultivated in the microfluidic device in the presence of glucose that were temporarily exposed to a sub-lethal dose of 2-DG. Here, observing that the drug decreased the cell division rate due to the reduction of the glycolytic flux, we found that the sensor’s GFP/mCherry ratio markedly increased, which indicates a lower FBP level and glycolytic flux (Fig. 8C). Remarkably, the glycolytic-flux-dependent modulation of the sensor’s GFP/mCherry ratio proved to be reversible as the readout recovered to the original level when the 2-DG was removed from the medium (Fig. 8C). Using the sensor’s mutant 2_6m as a negative control, we did not find pronounced changes in the readout in response to 2-DG treatment (Fig. 8D). Overall, these results demonstrate that the developed FBP sensor can distinguish cells with different glycolytic fluxes in clonal yeast cell populations, thus unveiling metabolic heterogeneity.

## DISCUSSION

In this work, we have developed an RNA-based sensor for the intracellular metabolite fructose-1,6-bisphosphate (FBP), which reports the flux through glycolysis. Specifically, we selected *in vitro* an RNA aptamer that binds FBP – a doubly phosphorylated and non-aromatic compound – selectively with millimolar affinity. Based on structural probing of this aptamer, we rationally designed a library of the aptamer-containing RNA devices that control GFP expression in yeast cells. Through FACS-Seq, combining cell sorting with next-generation sequencing, we identified an RNA device that responds to changes in the physiological range of intracellular FBP levels. With DMS-MaPseq, a new method for *in vivo* structural probing, we demonstrated that *in vivo* FBP binding is essential for the functionality of the sensor. Because the intracellular FBP level correlates with the flux through glycolysis (10, 46, 47), our developed RNA-based sensor can be used as reporter for glycolytic flux in single cells.

An important element of our work was the selection of an aptamer that is suitable for FBP binding in living cells. While the quality of aptamers is traditionally judged on the basis of on their K_D_ value (*i*.*e*. the lower, the better), here, we needed an aptamer with an affinity in the high micro- to low millimolar concentration range so that the physiological concentration changes of FBP could be transduced into differential sensor readouts. Still, the aptamer required high selectivity for FBP since many other phosphorylated sugars are simultaneously present in the cytosol, also at micro- to millimolar concentrations (47, 53). While the notion of a highly selective but low-affinity aptamer seems counterintuitive, interestingly, nature itself has evolved RNA-based regulatory systems, which are selectively controlled by highly abundant metabolites (54). For instance, the *glnA* riboswitch of cyanobacteria *Synechococcus elongatus* loosely binds L-glutamine in an open pocket; however, hydrogen bonds between the RNA and the heteroatoms of glutamine make the interaction highly specific (55). Similarly, since we found that neither fructose, fructose-1-phosphate nor fructose-6-phosphate could compete with FBP for binding to the selected aptamer (Fig. 1B, Supplementary Fig. S3), we conjecture that the two phosphate groups of FBP might play a crucial role in the observed selectivity.

While high-throughput methods make it possible to screen large sensor libraries, the transformation efficiency of yeast is a severe bottleneck in such efforts, limiting the library sizes that can be screened to around 10^5^ clones. To increase the effectiveness of our screening for *in vivo*-functional RNA devices, we implemented several measures for a more targeted search. First, as the *in vitro*-selected aptamer is a stem-loop and alters its tertiary structure upon FBP binding (Fig. 2B), we chose the hammerhead ribozyme (HHRz) as actuator domain. The ribozyme’s activity depends on tertiary-structure-stabilizing interactions between stem-loops I and II (35), and as such we reasoned that integration of the FBP aptamer into one of these stem-loops would modulate these interactions, and thus HHRz activity, by FBP binding. Second, we increased the diversity of the RNA devices by generating six different libraries, where in each of them we fused the aptamer to the actuator’s stem-loops in a different manner. By doing so, we mitigated potential biases in the library design that could have resulted in unspecific alterations of HHRz activity or in FBP binding. Third, the depletion of non-functional RNA devices from the library allowed us to concentrate our screening capacity on variants that can display FBP-dependent activity. Given the challenges offered by the restricted search possibilities, these measures allowed us to ultimately identify a sensor with FBP-dependent activity, suggesting that a clever combination of rational design and limited screening efforts here led to the successful development of the sensor.

In contrast to protein-based metabolite sensors, e.g. those relying on Förster resonance energy transfer, RNA-based sensors are in theory scalable (21), meaning that one should – in principle – be able (i) to *in vitro* identify an aptamer for any given metabolite and (ii) to engineer such aptamer into a sensor. However, to our knowledge in the last 10 years, this has been accomplished only in a few instances (24, 29, 30), indicating that there is not yet an obvious way to turn *in vitro*-selected aptamers into metabolite sensors that are functional in cells. It has been argued that this is due to limitations in screening throughput and efficiency (28). Yet, even with labor-intensive manual screening, Suess and colleagues have developed three (out of the eight identified thus far) *in vitro*-selected aptamers into *in vivo*-functional sensors (17, 18, 29), which suggests that throughput might not be the most critical bottleneck. In our work, we managed to develop an *in vivo*-functional metabolite sensor by exploiting FACS-Seq (35), which allowed us to test thousands of clones simultaneously and in an automated way. We believe that an essential factor for the efficient development of new RNA-based intracellular metabolite sensors is to optimize screening effectiveness by exploiting unique properties of the respective aptamer. We did this by choosing a suitable actuator, and fusing the aptamer in a manner that exploits its structural elements. Besides, the way the initial *in vitro* aptamer selection is done also seems to be important for successful development of sensors from *in vitro*-selected aptamers. This includes SELEX under *in vivo*-like conditions, the use of capture SELEX and selection monitoring by NGS (29), and the use of “naturalized” RNA pools, in which the variable regions are embedded in a scaffold derived from natural riboswitches (30).

The developed RNA-based biosensor and the one we recently created based on a transcription factor (TF) (56), both gauging FBP and glycolytic flux, now offer a toolbox of alternative methods that have different properties and can thus be chosen to suit particular applications. First, due to different mechanisms of action, the readouts of the two biosensors respond to changes of intracellular FBP levels in opposite directions: while the RNA-based biosensor decreases the expression of the fluorescent reporter, the TF-based one increases it as a response to elevated FBP. Second, although the RNA-based biosensor has a smaller maximal fold change of the readout compared to the TF-based one, the RNA-based biosensor exhibits the same resolution across the intracellular FBP levels employed to test both biosensors, or even a higher resolution if we also consider different FBP levels employed to test each biosensor (Supplementary Note 4). The developed RNA-based biosensor has a broader range of potential applications compared to the previously developed TF-based biosensor. First, ribozyme-based systems work in all kingdoms, from bacteria to human cells (35, 37, 57, 58), which makes the engineered biosensor applicable in other organisms. Second, the RNA-based biosensor, which controls reporter expression post-transcriptionally, responds to FBP changes faster than TF-based biosensors, allowing for more dynamic measurements. Third, RNA-based sensors are modular, meaning that the sensing aptamer can be transferred to different RNA-based scaffolds for other purposes (23–25, 59), and the fluorescent reporter can be exchanged by a different gene that, for instance, provides a selective advantage (36, 60) or that codes for a post-transcriptional regulator such as an sRNA (61, 62). Fourth, in synthetic biology, RNA-based systems with different specificities, including our system specific for FBP, can be assembled together and function as logic gates to relay different molecular inputs into a single output (23, 63, 64). Finally, the RNA- and TF-based biosensors could be implemented orthogonally or combined in the same system (36). In this regard, it would be interesting to adjust the sensors so that they responded to different FBP levels and created cells where a single input (glycolytic flux) could lead to different outputs (for instance, a toxin and a selective advantage) depending on the intracellular concentration of FBP.

Overall, with this work, we further pushed the RNA-based sensor development from the proof-of-concept stage to the demonstration that it is now possible to develop RNA-based sensors from scratch for a physiologically highly relevant, and even doubly phosphorylated, metabolite. The developed FBP sensor is remarkable because FBP is not just a normal metabolite, but in organisms across kingdoms it reports the flux through glycolysis (10, 46, 47). Reporting on metabolic activity, the developed sensor produces a highly relevant functional readout. As such, the sensor can now be employed for measurement of glycolytic flux in single cells, for instance, to address new research questions on metabolic heterogeneity. Also, it can be used in FACS-based screening approaches, *i*.*e*. to identify highly productive cell factories in a biotech context. Beyond, novel applications in synthetic biology can be envisioned, such as the engineering of genetic circuits where glycolytic flux serves as an input.

## METHODS

### *In vitro* selection of FBP aptamer

C45 was selected according to a previously described method (40). In brief, 100 μL of D-fructose-1,6-bisphosphate (FBP) immobilized to epoxy-activated sepharose by 1,4-butanediol-diglycidyl-ether equilibrated with cytosolic buffer *CB* (6 mM KH_2_PO_4_, 14 mM K_2_HPO_4_, 140 mM KCl, 10 mM NaCl, 5 mM MgCl_2_, 5.5% glucose, pH 6.88) was mixed with 1 nmol of the RNA pool in a final volume of 400 μL and let to settle by gravity at 25°C. In rounds 1 to 3, prior to the incubation with FBP-sepharose, the RNA pool was pre-cleared with the same volume of sepharose. After incubation, the column was washed twice with 600 μL of CB and the RNA bound eluted. Elution buffer was 5 mM EDTA in rounds 1 to 3, and 5mM FBP in CB in rounds 4 to 13. RNA was precipitated from the eluates with 30 mM sodium acetate in ethanolic solution and 10 μg of glycogen, retrotranscribed with Superscript II (Invitrogen; Carlsbad, CA) at 54°C for 10 min and the resulting cDNA amplified with GoTaq Flexi DNA polymerase (Promega; Madison, WI) for 8 PCR cycles in the same reaction with a final volume of 100 μL. 10 μL of the amplification product was used as template to synthesize the RNA pool for the next round by *in vitro* transcription with T7 RNA polymerase at 37°C for 20 minutes. RT-PCR products obtained from round 13 were cloned with TOPO TA Kit (Invitrogen; Carlsbad, CA) according to the manufacturer’s indications, and plasmids extracted from individual bacterial clones sent for Sanger sequencing using the T3 primer.

### Binding assays with immobilized FBP

CAL Fluor Red 610-labelled RNA sequences were purchased from LGC Biosearch Technologies (Risskov, Denmark). 560 µl of FBP-sepharose were mixed with 8.4 mL of 20 nM CAL Fluor Red 610-labelled RNAs in a Econo-Column Chromatography Column (BioRad, Hercules,CA), and the column washed with CB until no fluorescence change was detected in the flow-through. The resin was then resuspended in 8.4 mL of CB and evenly distributed among seven empty polypropylene spin columns, from which the RNA was eluted using different concentrations of FBP (1.2 mL; 0.03, 0.13, 0.50, 2.0, 8.0, 32, 128 mM). The percentage of eluted RNA was calculated by measuring the fluorescence intensity in the eluted fractions and the one remaining in their respective columns with an Edinburgh FS920 spectrofluorometer (Edinburgh Instruments, Edinburgh, U.K.) using an excitation wavelength of 590 nm (slit of 5 nm) and an emission of 615 nm (slit of 20 nm) in a final volume of 1.2 mL for all samples in a 10mm x 10mm quartz cuvette. Specificity assays were performed as described but using 200 µl of FBP-sepharose and 1.2 mL of 20 nM CAL Fluor Red 610-labelled RNA and eluted with 1.2 mL 10 mM of glucose-6-phosphate, fructose-6-phosphate, adenosine-triphosphate and FBP (all from Sigma; St. Louis, MO). All measurements were performed in triplicate. Data were analyzed using GraphPad Prism 5.0, where a non-linear regression (assumptions: one binding-site – specific binding) with the least squares fitting method was used to model binding behavior and the K_D_.

### Preparation of metabolites

To remove the Na^+^ present in commercially-available salts of phosphorylated metabolites (fructose-1-phosphate, fructose-1,6-bisphosphate) these compounds were dialyzed prior to binding analyses. The compounds were dissolved in 2x CB, and mQ H_2_O was then added to reach a final concentration around 1.5M in 1x CB. This stock solution was dialyzed in a Spectrum Micro Float-A-Lyzer with a MW cut-off 0.1-0.5 kDa (Thermo Fisher Sci.; Waltham, MA) against 1L of CB with additional 2M glucose to prevent osmosis. Dialysis was let to proceed for 24 hours with continuous stirring at 4°C including two changes of the outer buffer. After dialysis, the pH of FBP solution was checked and adjusted to 7.0 if necessary, and the concentration determined by HPLC. In brief, a series of dilutions of the dialyzed stock were run through a HiPLEX column (Agilent, Santa Clara, CA) for 20 minutes in 0.005 M H_2_SO_4_. A unique peak at 6.4 min was integrated and the values interpolated in a standard curve run alongside.

### *In vitro* transcription

DNA templates for *in vitro* transcription of RNA aptamers were generated by hybridization two complementary single stranded DNA oligonucleotides (ADO-01 to ADO-12; Supp. Table S7). 150 μL of each oligo at 200 μM in TE buffer (10 mM Tris-HCl, 1 mM EDTA, pH = 8) were mixed, and the mix heated at 95°C for 5 minutes and then cooled down to 70°C with a −1°C/min slope. Hybridization was monitored by poly-acrylamide gel electrophoresis (PAGE) in a 6% PA – 1x TBE (89 mM Tris, 89 mM boric acid, 2 mM EDTA) gel using 0.5x TBE as running buffer for 20 min at 300V. The gel was stained with a 1:10,000 solution of Invitrogen SYBR safe (Thermo Fisher Sci.; Waltham, MA) in 1x TBE for 5 min and visualized in a UV transilluminator.

0.1 µM of template DNA was transcribed *in vitro* for 4 hours at 37°C with 1 U/µl T7 polymerase. His-tagged T7 RNA polymerase (kindly provided by Beatrix Suess, TU Darmstadt) was expressed in *E. coli* and purified with Ni^2+^-sepharose column. The transcription mix also included, 1x transcription buffer, NTPs 2.5 mM each, 0.2 U/µL RNase inhibitor, 0.1 mU/µL inorganic pyrophosphatase (all from Thermo Fisher Sci.; Waltham, MA), 2 mM spermidine (Sigma, St. Louis, MO) in a final volume of 0.3 mL. To generate 5’-thio-modified RNA for RNA labeling, the reaction mix also included 10 mM guanosine-5’-thiophosphate (GMPS; Genaxxon bioscience, Ulm, Germany). Nucleic acids were precipitated with 0.3 M sodium acetate in ethanolic solution, and then template DNA digested using Turbo DNA-free kit (Thermo Fisher Sci.) for 1h at 37°C following manufacturer’s instructions. DNA-free RNA was precipitated with sodium acetate, and purified by preparative denaturing urea-PAGE. RNA pellets from 5’-thio-modified RNA transcription were re-dissolved in coupling buffer (8M urea, 2 mM Tris-HCl, 20 mM EDTA pH = 7.0), and DTT was then added to a final concentration of 60 mM. After a 1 hour-incubation, un-incorporated GMPS and DTT were removed by gel filtration using two Zeba Spin desalting columns (Thermo Fisher Sci.) sequentially.

### RNA labeling

50 nmol of Cy5-maleimide (GE Healthcare, Chicago, IL) dissolved in anhydrous DMSO (Thermo Fisher) were mixed with 150 nmol of RNA (from 5’-thio-modification procedure) in coupling buffer and incubated for 2 h at 37°C in the dark. Cy5-labeled RNA was purified by preparative denaturing urea-PAGE. The integrity and purity of labeled RNA was monitored by denaturing PAGE, being the gels stained with SYBR safe and scanned with a Typhoon 9410 (two channels: 488/520 for SYBR; and 640/674 nm for Cy5) and analyzed with ImageQuant TL1 Software (GE Healthcare). Cy5-RNA concentration was determined in a Spark 10M multimode reader using a nanoquant plate (Tecan, Männedorf, Switzerland).

### Preparative denaturing urea-polyacrylamide gel electrophoresis

DNA-free RNA from in vitro transcription and Cy5-labeled RNA from coupling reactions precipitated with sodium acetate were re-dissolved in Bartel/Lau loading buffer (8M urea, 2 mM Tris-HCl, 20 mM EDTA pH = 8.0), heated 2.5 min at 90°C and snap-chilled in an ice-water bath. The sample was loaded on a 16×16 cm 8M urea – 10% PA – 1xTBE gel, previously pre-run at 250V for 40 min, and run in 1x TBE for 2.7 h at 300V in the dark. Gels were then shaded with an UV lamp to identify the bands corresponding either to labeled or unlabeled RNA. These bands were sliced from the gel, freeze-thawed once and crushed with a tip, and the RNA eluted in 500 µL of 0.3M sodium acetate pH = 5.8 for 2h at 50°C in the dark. Eluted RNA was precipitated, re-dissolved in RNA storage buffer (*RSB*: 3mM sodium acetate, 0.25 mM EDTA, pH = 6), heated for 2.5 min at 90 °C and snap-chilled, aliquoted and stored at −80°C.

### Binding assays with microscale thermophoresis

A frozen aliquot of 0.2 µM Cy5-labeled RNA in RSB was thawed in ice, heated 2.5 min at 90°C, snap-chilled and kept at 0°C for 3 min. This denatured RNA was immediately let to fold for 1h at 30°C in 1x CB with 0.5 mg mL^−1^ BSA at final RNA concentration of 15 nM. The ligand was diluted in 1x CB and then used to prepare serial dilutions with the same buffer, except for the lowest concentration point that contained only the buffer (control). To assemble the binding reactions 8 µL of the folded labeled RNA was mixed with the same volume of the ligand at 25°C and promptly used to fill up the capillaries for MST measurements. MST measurements were performed using a Monolith NT.115 Green Red MST instrument (Nanotemper technologies, München, Germany) with MST grade standard treated NT.115 capillaries. The MST settings were: temperature of the instrument: 22°C; LED power: 90%, Red – Ex: 625 nm, Em: 680 nm; IR/MST laser power: 60%; time before heating: 5 sec; time IR/MST on: 30 sec; time after heating: 3 sec. The normalized fluorescence (F_norm_) corresponds to the ratio between the fluorescence measured after the 30 sec period where IR/MST-Laser has been on (F_hot_) divided by the fluorescence measured before the IR/MST-Laser starts heating (F_initial_), i.e. F_norm_ = F_hot_/ F_initial_. F_norm_ measurements were obtained with NTA Analysis software (NanoTemper Technologies) and normalized (ΔF_norm_) by subtracting the value of the lowest concentration point (i.e. the only-buffer control). Data were analyzed using GraphPad Prism 5.0, where a non-linear regression (assumptions: one binding-site – specific binding with Hill slope) with the least squares fitting method was used to model binding behavior and the K_D_.

### Selective 2’-hydroxyl acylation analyzed by primer extension

DNA templates for *in vitro* transcription in SHAPE assays included the T7 promoter, two DNA cassettes to prevent signal loss from C45 (65), the C45 aptamer sequence, and the first 218 nucleotides of GFP coding sequence. C45 aptamer sequence was first cloned into pWHE601 immediately upstream of ATG initiation codon of GFP (17). To this aim, two DNA fragments corresponding to C45 and the linearized plasmid were produced by PCR with primers ADO13 and ADO14, and ADO15 and ADO16, respectively, and assembled using the Gibson Assembly mix (New England Biolabs, Ipswich, MA). pWHE601-C45 was then used as template to generate an amplicon that includes one cassette and C45 using ADO17 and ADO18 primers. In a second PCR, the T7 promoter and the second cassette were included using primers ADO19 and ADO20, and this amplicon was then used as template to generate the final template using ADO21 and ADO20 primers. All PCR reaction were carried out using the Q5® High-Fidelity DNA Polymerase (New England Biolabs) following manufacturer’s recommendations. RNA was transcribed *in vitro* and purified as described above.

For RNA folding, a frozen aliquot of unlabeled RNA (diluted in RSB) was thawed in ice, denatured as described above, and then diluted in 1x CB either with or without 86 mM FBP. RNA was let to fold for 1h at 37°C. 5 pmol of folded RNA was treated with 5 mM 1-methyl-6-nitroisatoic anhydride (1M6, dissolved in anhydrous DMSO) for 3 min at 37°C, and the reaction quenched with cold 0.3M sodium acetate in ethanol and precipitated. Control samples containing no reagent were prepared in parallel with an equivalent volume of DMSO. For primer extension, RNA pellets (1M6-modified or untreated) were dissolved in 10 µL of RNAse-free H_2_O and mixed with 2 µL of a 1.25 µM NED-labeled ADO-22 primer at 5’, 4.5 µL of 5x first strand buffer (Invitrogen), 1 µL of 100 mM DTT, 1 µL of dNTPs mix 10 mM, 0.2 µL of RNAse inhibitor and 0.3 µL of SuperScript III reverse transcriptase (Invitrogen). The reaction was carried out for 20 min at 52°C and quenched at 85°C for 5 min. Sequencing reactions included 5 pmol of un-modified RNA, 2 µL of 1.25 µM 6-FAM-labeled ADO-22 primer and 1 µl of 100 mM ddTTP (USB-Affymetrix, Cleveland, OH). Fluorescently-labeled cDNA fragments were analyzed by capillary electrophoresis (CE). A sequencing reaction and a sample under study (RNA treated with 1M6 or only DMSO) were loaded in each capillary and analyzed in a 3130xl Genetic Analyzer (Thermo Fisher Sci.) following a prior optimization of the ratio between experimental and sequencing reactions to avoid signal saturation.

To calculate the reactivity values for each nucleotide the electropherograms from CE were processed using QuShape software (66). In specific, a region of interest was first delimited excluding the final product to prevent signal saturation. Individual peaks were obtained by smoothing the signal and adjusting the baseline in this region. Signal decay correction was omitted. After aligning the peaks from the experimental and control samples, the sequence (provided as input) was aligned to these peaks. The normalized reactivity at each nucleotide position was calculated by dividing each value by the average intensity of the 10% most reactive residues (excluding outliers) (66). To calculate ligand-induced structural changes (differential shape) the reactivity value at each nucleotide position was subtracted from that in the RNA folded in the presence of FBP. Differences in the reactivity were considered when it was greater than 0.3 and the p-value (t-test, two tails, non-paired) smaller than 0.05.

For the secondary structure predictions, the reactivity values were send to RNA structure software (43), which coupled SHAPE reactivity with energy minimization.

### Yeast strains and cultivation

We used the following *S. cerevisiae* strains: KOY wild type (WT), KOY-VW100 *HXT7* (*HXT7*), KOY-VW100 *TM3* (*TM3*), KOY-VW100 *TM4* (*TM4*) and KOY-VW100 *TM6\**(45) (*TM6\**), all with *ura3-52* auxotrophy, plus WT and *TM6\** without *ura3-52* auxotrophy for microscopic analyses. Unless specifically indicated, yeast were cultivated in 500 mL-flasks containing 50 mL of Verduyn’s minimal medium (67) buffered at pH 5 with 10 mM KH-phthalate, and inoculated with exponentially growing cells. Starting cell density (0.75-2.0x 10^6^ cells mL^−1^) was adjusted to the foreseen time of cultivation and each strain’s growth rate so that the culture never achieved densities higher than 2.5-3.0x 10^7^ cells mL^−1^. To obtain a homogeneous culture of cells adapted to the carbon source and with a metabolic steady state, cells were allowed to grow at least for 20 generations through two pre-culturing steps prior to the main culture. The inoculum was prepared in the identical minimal medium. All cultivations were performed at 30 °C, and cultures were continuously shaken at 300 rpm. To make competent cells, we used Yeast Peptone broth (20 gL^−1^ peptone, 10 g L^−1^ yeast extract). To recover cells after sorting, we used Yeast Nitrogen Base (YNB) supplemented with Complete Supplement Mixture (CSM) dropout without uracil and 400 μg mL^−1^ of G418. All nutrients and supplements to make media were purchased to Foremedium™ (London, UK). All culture media were supplemented with either 20 g L^−1^ of D-glucose or maltose (both from Merck Millipore).

For microfluidic cultivation, several days before a microscopy experiment, we recovered necessary strains from their glycerol stocks stored at −80°C, by growing them for two days on 2%-glucose agar plates. For each strain, a single colony was inoculated in 10 mL of Verduyn’s minimal medium in a 100-mL Erlenmeyer flask, which initiated a pre-culture that was cultivated overnight at 30°C with shaking at the speed of 300 rpm. Cells still growing exponentially in the pre-culture were diluted at the OD_600_=0.025-0.05 in the fresh medium and cultivated for several hours under the same conditions until the OD_600_=0.1-0.2. Afterwards, cells were loaded in the microfluidic device as described previously(51, 52). In the microfluidic device, cells were constantly provided with the fresh medium at the flow rate of 4-5 µL/min via a syringe pump (for Figures 8A, B, D) or an air-pressurized pumping system (OB1, Elveflow) assisted by a flow sensor (MFS2, Elveflow) (for Figure 8C). During cultivation in the microfluidic device, the temperature was maintained at 30°C with the help of a microscope incubator. Cell cultivation in the two consecutive flask cultures and the microfluidic device was done on the same carbon source, namely, 2% (20 g/L) glucose or 2% maltose. For Figures 8A-B, we cultivated different strains simultaneously in two microfluidic devices attached to one cover glass: in one device, we loaded *TM6\** cells with the sensor 2_6, and *TM6\** cells without fluorescent proteins and *ura3-52* auxotrophy (for further cell-autofluorescence correction); in the other device, we loaded *TM6\** cells with the mutant 2_6m, and also *TM6\** cells without fluorescent proteins and *ura3-52* auxotrophy. For Figures 8C-D, we cultivated WT cells with the sensor 2_6 and the mutant 2_6m in microfluidic devices on different days. In both cases, we also loaded into the microfluidic device WT cells without fluorescent proteins and *ura3-52* auxotrophy for further cell autofluorescence correction.

For *in vivo* RNA structural mapping, *TM6\** cells with the sensor 2_6 and the mutant 2_6m were cultivated in two consecutive exponential cultures in 10 mL of Verduyn’s minimal medium with 2% glucose and 2% maltose in 100-mL Erlenmeyer flasks.

### Quantification of intracellular concentration of fructose-1,6-bisphosphate

3x 10^7^ *TM6\** cells from a exponentially-growing culture in minimal medium plus 2.0 g L^−1^ glucose or 2.0 g L^−1^ maltose were quenched in methanol at −80°C, harvested by centrifugation at 2,500x g for 10 min at – 10°C and the pellet washed with methanol and stored at −80°C. Extraction of cellular metabolites was done in a solution of methanol, acetonitrile and water (4:4:2 by volume) with 0.1 M formic acid, pre-cooled to – 20°C and with U^13^C internal standards spiked in. The pellet was extracted twice with 0.9 mL of the solution for 10 minutes at −20°C with shaking (final volume 1.8 mL). Organic solvents were removed from the supernatant in a vacuum concentrator (2 h, RT), and the remaining water by freeze-drying overnight. The precipitates were resuspended 0.2 mL of water and re-dissolved by vortexing and ultrasonication. Finally, the extract was centrifuged (2 min; 21,000x g,4°C), and 10 µL of the supernatant analyzed using a UHPLC-MS/MS. Quantification was based on the ^12^C/^13^C peak area ratios related to a calibration curve made of pure ^12^C-standards with a global U^13^C-internal standard. A detailed description of the UHPLC-MS/MS method and equipment used for LC-MS/MS and preparation of the standards has been previously described (68). FBP concentration in each of the 2 biological replicates, which corresponded to the average of independent FBP determinations from two samples of the same culture, was then used to calculate the average FBP-amount per cell.

To determine cell volume we used BudJ (69), a plugin for ImageJ. In brief, the algorithm determines cell boundaries by detecting two consecutive maximum and minimum pixel values in a densitometric profile along a radial axis from a seed point within the cell, and then fits an ellipse to the boundary pixel array. The measured major and minor axes from the ellipse are utilized to calculate the volume assuming that the cell has the shape of a prolate ellipsoid. At least 70 cells were selected from each biological replicate to calculate the median cell volume. Intracellular FBP concentration for each experiment was calculated by dividing the average FBP-amount per cell by the median of cell volume measurements.

For Figure 6D-E, to match the intracellular FBP concentrations with the GFP/mCherry ratios in the wildtype, *HXT7* and *TM6\** strains, we implemented the following conversion: 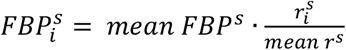, where 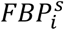 is the inferred FBP concentration of the strain *s*’s replicate culture *i* whose GFP/mCherry ratios are shown on the y-axis, *mean FBI*^*s*^ is the mean FBP concertation of the strain *s* reported in (48), 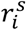 is the growth rate in the strain *s*’s replicate culture *i*, and *mean r*^*s*^ is the mean growth rate across three replicate of the strain *s*. To infer the intracellular FBP concentration in the *TM3* and *TM4* strains, we used the knowledge that their glucose uptake rates, glycolytic fluxes and growth rates are between the respective values of the *HXT7* and *TM6\** strains (45) and, building on the flux-signaling property of FBP, implemented the conversion: 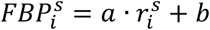, where *a* and *b* are the slope and intercept in the linear regression between the mean FBP concentrations and mean growth rates of the *HXT7* and *TM6\** strains.

### Determination of GFP/mCherry ratio by flow cytometry

GFP and mCherry fluorescence intensity was measured by flow cytometry (BD Accuri™ C6, Becton Dickinson) at three different time points on the hour scale where cells displayed their normal growth rate. Optical configuration to measure GFP and mCherry signals was 488 nm for excitation and 510/15 BP and 610/20 BP filters, respectively, for detection. Data were analyzed using Kaluza (Beckman Coulter) or Python applied in the same manner. Live cells were first gated by FCS and SSC height parameters, and duplets discarded using SSC width and FSC height parameters. Channels were compensated to remove GFP fluorescence spillover into mCherry signal using the double negative and single-color controls described for cell sorting (see above). GFP/mCherry plots were generated with 20,000 or a specified number of cells of this gate of singlets and live cells, and overlaid with other samples for comparison. To determine the ratio, we exported the compensated raw data from the cells in the main cloud (excluding cells with low mCherry) and calculated single-cell GFP/mCherry values.

### Preparation of C45-HHRz library and HHRz versions for the *in vivo* reporter system

To generate C45-HHRz libraries, and the different HHRz versions used as controls for the calibration of the *in vivo* reporter system, the inserts were amplified by PCR, digested with AvrII and XhoI and ligated to pCS1748 plasmid (34), which was previously digested with the same enzymes and de-phosphorylated with Antarctic phosphatase (New England Biolabs). The templates for the inserts of libraries C45-I and II, inv-I and II, and the wild-type satellite RNA of tobacco ringspot virus (sTRSV) HHRz control were generated after five amplification cycles of a SOE PCR, using 10 fmol of each F and R 3’-overlapping denatured primers. The optimal T_m_ was determined experimentally for each primer pair and a fifth of the volume of this reaction used as template for the amplification of the insert. For libraries sh-I and II and all HHRz-mutant versions we used 0.1 fmol of a unique ssDNA primer as template (ADO41-42 and ADO25-32, respectively). Insert amplification was run for 20 cycles, using 0.5 µM of each primer ADO043 and ADO044 and a T_m_ of 65°C. All amplifications were carried out using Q5 polymerase (New England Biolabs) being the PCR protocol set according to manufacturer’s recommendation. 100 ng of digested and de-phosphorylated pCS1748 were ligated to a 3-fold molar excess of each digested insert overnight at 16°C. Six ligation reactions were then pooled and precipitated with 10 µg of glycogen, re-dissolved in 5 µL of ddH_2_O, transformed into ultracompetent DH5α cells (70) by thermal shock, and the recombinants selected in LB with 100 µg mL^−1^ carbecillin. In these conditions we obtained around 2×10^5^ clones per library.

To transform the libraries and HHRz versions into the KOY wild-type and *TM6\** yeast strains we adapted the LiAc Gietz’s transformation protocol (71) so as to obtain at least as many as 4-fold the maximum theoretical number of clones for each library. In brief, to transform library LibII (containing 7 positions with randomized sequence and a theoretical maximum of 49,152 clones) into *TM6\** (displaying a 10-fold lower transformation efficiency than the wild type) we incubated 2×10^10^ cells with 18 µg of the plasmid library in 65 mL of a solution of lithium acetate/PEG/denatured salmon sperm carrier DNA (0.1M/33%/0.28mg mL^−1^) in TE pH=7.5 for 45 min at 42°C. Cells were recovered for 5 hours at 30°C in YP medium with 2% maltose and then spread onto 50 petri dishes (150×20 mm, Sarstedt) with YNB + CSM Ura^−^ with 2% maltose supplemented with G418 at 0.2 mg mL^−1^. After 72 hours of incubation more than 200,000 clones were scrapped from the plates and pooled.

### Sorting of cell subpopulations based on GFP/mCherry ratio by fluorescence-assisted cell sorting

Prior to the sorting each strain had grown for 20 generations in minimal medium without achieving a density higher than 2.5x 10^7^ cells mL^−1^ and displayed a constant and reproducible growth rate. Cell density was determined by flow cytometry using an Accuri™ C6 (Becton Dickinson). For each sorting experiment, a double negative (non-transformed strain), and two strains harboring single-color plasmids (GFP, pCS1585) and (mCherry, pCS1749) (34) were taken along as compensation controls. In addition, strains harboring the dual-color plasmids with the HHRz versions showing different GFP/mCherry values (HHRz, CTL, and M1 to M7) were also grown in parallel as calibration standards and used to set the sorting gates. All controls were generated in the both KOY wild-type and *TM6\** backgrounds. Libraries LibI and LibII were pooled at a proportion of 1:63. Cell cultures were spun down and resuspended to a density of of 5x 10^7^ mL^−1^in fresh medium, filtered through a 35 µm cell strainer (BD Falcon), and kept at 4°C until sorted. Cell sorting was carried out with a MoFlo Astrios (Beckman Coulter) using a 488 nm laser and a 513/26 bandpass emission filter to detect GFP, and a 561 laser and a 614/20 filter to detect mCherry. Live cells were first gated using forward-(FSC) and side-scattered light (SSC) height parameters, then the duplets discarded using SSC width and FSC height parameters, and finally only cells with intermediate mCherry levels within the main cloud were gated. For the pre-selection of C45-HHRz RNA devices with a functional ribozyme we sorted the 3-4% cells with lowest GFP values that did not overlap with the HHRz mutant in the active site (CTL version). For the high-throughput *in vivo* screening of RNA devices the top and bottom sorting gates corresponded to the inactive CTL mutant and the wild-type versions of HHRz, respectively, and the remaining four gates were evenly distributed throughout GFP/mCherry space in between. Sample data were acquired with Summit software (Beckman Coulter). We routinely collected 150,000 cells per sorted sub-populations. For minority subpopulations representing 0.01 to 0.2 % of the input population we collected 2,000 and 20,000 cells, respectively. Sorted cells were collected in 0.5 mL of minimal medium containing either 2% glucose or maltose and 30 µg mL^−1^ chloramphenicol at 4°C, and then diluted in 7 mL of YNB+CSM Ura^−^ +2% glucose or maltose and the culture let to grow and passed 1/50 twice.

### Preparation of DNA library for NGS

NGS library was prepared following a three-step PCR amplification scheme: (i) the *outer* PCR, where the region of the plasmid containing the RNA device was amplified, (ii) the *Nextera* PCR, a nested PCR that incorporated universal 5’ and 3’ tails for Illumina sequencing, and (iii) an indexing PCR to include sample-specific indexes as well as the Illumina adaptors. Plasmids were extracted from a 10 mL culture (5 OD) of sorted cells subpopulations using the Zymoprep™ Yeast Plasmid Miniprep II (Zymo Research) following manufacturer’s instructions. Outer PCR was performed with 10^6^ template DNA molecules and 0.5 μM of each ADO47-48 primers for 10 cycles. For nextera PCR, 10^6^ molecules of the amplicons resulting from outer PCR were used as template in a 21 cycle-amplification using 0.5 μM of each ADO50-51 primers. For absolute DNA quantification we used real-time PCR (qPCR). qPCR mix included 0.5 μM of each ADO47-48 (for plasmids) or ADO50-51 primers (for out PCR amplicons), 4 μL of a dilution of the DNA template, and 5 μL of 2x Power SYBR Green PCR Master Mix (Thermo Fisher), and the reaction run for 30 cycles in a CFX Connect™ Real-Time PCR Detection System (Bio-Rad). For absolute quantification, a calibration curve was generated with purified amplicons from pCS1748-CTL HHRz.

A diversity control (35) was spiked into each plasmid preparation (1 molecule of the diversity control per 2,000 of the plasmid) at the point of the outer PCR. It was generated using ADO049 primer as template in an outer PCR amplification, and contained a 17 nucleotide-stretch with a randomized sequence. This kind of control has been previously used to compute the typical number of reads resulting from any single molecule that existed in every sample at that point (35). Here, each outer PCR reaction contained a total of 500 molecules of the diversity control, all of them most likely with a different unique sequence. In addition, we included an additional sample with the plasmid pCS1748-CTL HHRz as a cross-contamination control. All PCR amplifications were carried out with the Q5® High-Fidelity DNA Polymerase (New England Biolabs) following the protocol recommended by the manufacturer, the T_m_ optimized empirically for each primer pair, and the PCR products purified using the NucleoSpin Gel and PCR Clean-up (Macherey-Nagel). Indexing PCR and NGS were performed by Microsynth (Balgach). In brief, purified products from indexing PCR were quantified using PicoGreen (Thermo Scientific) and pooled together (pooling factor for input samples was 7x, while for the rest of the samples was 1x). The paired-end sequencing was done on Illumina NextSeq 500 mid-output run 150v2 with a capacity of 120 million reads (passed filters), and the reads trimmed and de-replicated prior to our specific analysis.

### Analysis of high-throughput *in vivo* screening (FACS-Seq) of RNA devices

All analysis was performed in Python with the help of the modules *pandas, scipy, regex, matplotlib, matplotlib_venn* and *seaborn*. The sequences of the cross-contamination control, diversity control and potential C45-HHRz clones were identified in the NGS data via regular expressions, allowing a small number of matching errors (Supplementary table S3). Sequences failing to match these three patterns represented a small proportion of unique sequences (median 1.5e-02) as well as of total coverage (median 1.3e-03), did not show biases to specific samples (Supplementary table S4), and were ignored in further analysis. The cross-contamination rate was estimated in all samples and had the low median value of 4.2e-06 (Supplementary table S5). Detecting the sequences of diversity control, we verified that a single molecule at the point of the outer PCR was typically supported by two reads in the majority of the samples (Supplementary table S6). From the sequences identified as potential C45-HHRz clones (Supplementary table S3), the middle part between the primer-binding sites for *Nextera* PCR was extracted and put in a table with the number of corresponding reads (coverage) in every sample. Identical sequences in the table were collapsed, with their coverages summed up. We removed from the table those sequences that did not comply with any of the six RNA-device architectures (Supplementary table S3), which amounted to 403 out of the clones covered by at least four reads in both *TM6\** replicates on both carbon sources (the fraction of 0.026), 417 out of the clones covered by at least four reads in both wild-type replicates on both carbon sources (the fraction of 0.029) and 742 out of the clones covered by at least two reads in both wild-type replicates on both carbon sources (the fraction of 0.024). Calculation of the measures used for the hit selection (Supplementary table S2) was performed in this table, parts of which corresponding to different strains and coverage thresholds are provided in *Supplementary files*. Supplementary notes 2 and 3 motivate the choice of coverage thresholds, describe quality controls and present the hit selection in more detail.

### Validation of FBP-sensor candidates

Construction of individual clones identified as sensor candidates in the screening was carried out following the same strategies and experimental conditions as for the preparation of C45-HHRz library (see above). For hits 2_12 and 3_7, we proceeded as for sh libraries using a unique ssDNA primer as template, while for the rest of the constructs we proceeded by SOE PCR with a forward and a reverse primer pair just as we did for C45 or inv libraries. Sequences were verified by Sanger sequencing using ADO77 primer. Plasmids were transformed into KOY wild type, *HXT7, TM3, TM4* and *TM6\** strains using the Lithium acetate/PEG method as described above.

Individual clones were grown in minimal medium as described for the sorting of cell subpopulations (see above), 2 mL in 14-mL round-bottom tubes (Greiner) (Figure 6A-C) or 10 mL in 100-mL flasks (Figure 6D-E). To determine GFP/mCherry ratio, we measured GFP and mCherry by flow cytometry (BD Accuri™ C6, Becton Dickinson) as described above. For each biological replicate experiment (Figure 6A), the *response* of a sensor to the FBP-level difference was calculated as the mean value of the change in the median of single-cell GFP/mCherry ratio values across three time points (see the caption of Figure 6A-B). Using the average across the three time points of normal growth is intended for obtaining a robust estimate.

To uncover the dependency of the GFP/mCherry ratio of the sensor 2_6 and mutant 2_6m upon the intracellular FBP concentration in five strains, namely wild type, *HXT7, TM3, TM4* and *TM6\**, we cultivated each of these strains in three replicate cultures provided with 2% glucose as the carbon source. Each of the replicate cultures were propagated in three consecutive exponential cultures, with the data of the latest ones used to make Figure 6D-E.

### *In vivo* RNA structural mapping

#### In vivo DMS treatment

50 mM HEPES-KOH, pH 7.5, was added to a culture sample. Afterwards, we treated cells in the sample by the addition of 100 mM DMS (Sigma Aldrich, cat. D186309), and incubated the sample at 30°C with moderate shaking for 2 minutes. Next, DTT was added to the final concentration of 0.5 M to quench the reaction, after which the cells were centrifuged at 3,000 g for 2 minutes (4°C), and the medium discarded. We washed the cells once in ice-cold PBS supplemented with 0.5 M DTT. Pellets were snap-frozen in liquid nitrogen and stored at −80°C.

#### Targeted DMS-MaPseq analysis

Total RNA was extracted by the addition of TRIzol reagent (ThermoFisher Scientific, cat. 15596018) to cell pellets, followed by the incubation at 60°C for 30 minutes. RNA purification was performed following manufacturer instructions with minor changes. Briefly, after the addition of 0.2 volumes of chloroform and centrifugation, the upper aqueous phase was added to 2 volumes of 100% ethanol, and then loaded onto an RNA Clean & Concentrator-5 column (Zymo Research, cat. R1013). Before proceeding to the reverse transcription, samples were treated for 30 minutes at 37°C with 1 U TURBO DNase (ThermoFisher Scientific, cat. AM2238) to remove traces of genomic DNA. Targeted reverse transcription was performed using 100U TGIRT-III (InGex, cat. TGIRT50) in FSB buffer (50 mM Tris-HCl pH 8.3, 75 mM KCl, 3 mM MgCl_2_]) supplemented with 5 mM DTT, 1 mM dNTPs, 1 μM gene-specific RT primer ADO79, 10 U SuperaseIN RNase Inhibitor (ThermoFisher Scientific, cat. AM2694). Briefly, after mixing RNA with the primer and dNTPs, the sample was heated at 70°C for 5 minutes and then immediately transferred to ice for 1 minute. After the addition of the remaining components (but TGIRT-III), reactions were further incubated at room temperature for 5 minutes. TGIRT-III was added, and samples were incubated for 2 hours at 57°C, followed by the addition of 1 μl NaOH 5N and the incubation at 95°C for 3 minutes to release the cDNA. Samples were then purified on RNA Clean & Concentrator-5 columns and 20% of the eluate was used for targeted PCR amplification (ADO78, ADO79).

PCR amplicons were randomly fragmented to an average size of 150-200 bp using NEBNext dsDNA Fragmentase (New England Biolabs, cat. M0348S), following manufacturer instructions. Samples were then purified using 1 volume of Agencourt AMPure XP beads (Beckman Coulter, cat. A63880) and used as input for library preparation using NEBNext Ultra II DNA Library Prep Kit (New England Biolabs, cat. E7645S), according to manufacturer instructions.

#### Data analysis

Reads were mapped to the respective reference sequences using the rt-map tool of RNA Framework (72) and Bowtie v2 (73), after trimming low quality bases (Phred < 20). Per-base mutations and coverage, as well as mutation map (MM) files for spectral deconvolution, were calculated using the rf-count tool (parameters: -na -ni -md 3). Normalized reactivity profiles were computed using the rf-norm tool, by applying box-plot normalization (parameters: -rb AC -n 1000 -sm 4 -nm 3). MM files were instead used as input for DRACO to perform spectral deconvolution of alternative conformations (Moranti *et al*., manuscript in preparation).

### Microscopy and image analysis

A cover glass with one (Figure 8C-D) or two (Figure 8A-B) microfluidic devices was mounted to a Nikon Eclipse Ti-E inverted wide-field fluorescence microscope, where time-lapse imaging of cells was performed. The microscope was equipped with an Andor DU-897 EX camera, a 40x (Nikon CFI Super Fluor 40X Oil; NA = 1.3) and 100x (Nikon CFI Super Fluor 100XS Oil; NA = 0.50-1.30) objectives, as well as a Lumencor AURA excitation system. Imaging was done in four channels. Specifically, in NAD(P)H channel, we excited cells at 360 nm, employing a 350/50-nm band-pass filter, a 409-nm beam-splitter and a 435/40-nm emission filter, while having 2% of the maximal light intensity and 150 ms exposure (the corresponding fluorescence data was not used in the analysis). For GFP measurements, we excited cells at 485 nm, using a 470/40-nm band-pass filter, a 495-nm beam-splitter and a 525/50-nm emission filter, while having 2% of the maximal light intensity and 100 ms exposure (GFP channel). For mCherry measurements, we excited cells at 560 nm, using a 560/40-nm band-pass filter, a 585-nm beam splitter and a 630/75-nm emission filter, while having 2% of the maximal light intensity and 100 ms exposure (mCherry channel). For bright-field imaging, a halogen lamp produced light that was filtered with a 420-nm beam-splitter to exclude UV before illuminating cells (BF channel). The microscopes were operated using NIS-Elements software. We set the Readout Mode to 1 MHz to minimize the camera readout noise and fixed the baseline level of the cameras to 500 at −75 °C. The Nikon Perfect Focus System (PFS) was used in time-lapse imaging to prevent the loss of focus set in the beginning of the experiment. Microscopy imaging of WT strain grown on glucose and *TM6\** strain grown on maltose was done with the time steps of 5 minutes, whereas, in case of *TM6\** strain grown on glucose, we used the time step of 15 minutes. For Figures 8A-B, the 40x objective was used, whereas for Figures 8C-D we employed the 100x objective.

In every microscopy experiment, multiple non-overlapping regions in the XY-plane of the microfluidic device(s) were imaged, which resulted in a set of Nikon NIS-Elements ND2 files each containing a multi-channel movie for one XY-region. Every ND2 file was imported into ImageJ (v.1.52n, Java 1.8.0_66) (74) where images in the bright-field channel were sharpened and contrast-enhanced, after which the movie was saved as a TIFF file. Cells were tracked throughout the movie, and their mother compartments segmented by fitting an ellipse in the bright-field image at each time point via the semi-automated plugin BudJ (69) used with ImageJ (v.1.49v, Java 1.8.0_144). Simultaneously, by visual inspection and with the help of a custom macro, we recorded for each segmented cell the time points of budding events (appearance of a dark-pixel cluster from which a daughter cell would later grow) and death (abrupt shrinking and darkening of the cell, cessation of cytoplasmic movement, after which the data from the cell was not used). To analyze cellular fluorescence data, we loaded the movie-containing TIFF file into a *NumPy* (v.1.15.4) multidimensional array via Python’s module *scikit-image* (v.0.13.1) (75). For each time point and XY-region, we implemented background-correction by subtracting from every pixel value the mean of 2.5% dimmest pixels in the field of view. We extracted the pixels corresponding to a cell of interest by overlapping the NumPy multidimensional array with the segmentation ellipses provided by BudJ. The fluorescent signal of a cell was defined as the mean value of its background-corrected pixels. To implement cell-autofluorescence correction, for every time point in both GFP and mCherry channels, we calculated the median signal of cells lacking fluorescent proteins, smoothed it using LOWESS regression with 70 data points to fit a local line, and, at the same time point, subtracted the resulting value from the signals of cells expressing the sensor or the mutant. Prior to computing the GFP/mCherry ratio, we smoothed the cell-autofluorescence-corrected GFP and mCherry signals, using LOWESS regression with 8 data points to fit a local line.

## AUTHOR CONTRIBUTIONS

YL selected the aptamer and performed binding experiments with immobilized FBP mentored by FE and supervised by LFO and GM. ADO performed and analysed binding experiments by microscale thermophoresis. NM-F performed and analysed SHAPE experiments. SB-V quantified intracellular FBP. ADO and SB-V characterised the *in vivo* reporter system, prepared C45-HHRz libraries in *E. coli* and yeast, and performed the pre-selection and sorting of yeast libraries. ADO prepared DNA library for NGS. VT designed and implemented the data analysis for the high-throughput screening with a conceptual contribution from ADO. ADO, SB-V and VT validated sensor candidates. VT investigated the sensor’s dynamic range across multiple strains. DI implemented and analysed *in vivo* RNA structural probing. VT performed and analysed microscopy experiments. MH conceived the project and secured funding for it. ADO designed and directed the overall course of the study. ADO, VT and MH wrote the manuscript.

## ACKNOWLEDGEMENTS

The research leading to these results has received funding (MH) from the European Union Seventh Framework Programme (FP7-KBBE-2013-7-single-stage) under grant agreement no. 613745, Promys, from the ITN project MetaRNA under grant agreement no. 642738, and from the ITN project ISOLATE under grant agreement no. 289995. The authors would like to thank Prof. Arnold Driessen for kindly sharing the Monolith for MST analyses, Prof. Christina Smolke and Dr. Brent Townshend for plasmids to implement the *in vivo* reporter system and useful discussions, Dr. Alicia Barroso for capillary electrophoresis analysis of SHAPE samples, Dr. Athanasios Litsios for initial single-cell microscopy analyses and critical discussions, and Dr. Andreas Milias-Argeitis, Prof. Antonio Murciano, Dr. Jakub Radzikowski and Joana Saldida for input on data analysis.

## DECLARATION OF INTERESTS

The authors declare no competing interests.

## DATA AVAILABILITY STATEMENT

The datasets generated during and/or analyzed during the current study are available from the corresponding author on reasonable request. Strains and plasmids are available from the corresponding author on reasonable request.

## CODE AVAILABILITY STATEMENT

Computer code used to analyze the data are available from the corresponding author on reasonable request.

## Notes

### Competing Interest Statement

The authors have declared no competing interest.

## REFERENCES

1. Kim, J. and DeBerardinis, R.J. (2019) Mechanisms and Implications of Metabolic Heterogeneity in Cancer. Cell Metab., 30, 434–446.

2. Schreiber, F. and Ackermann, M. (2020) Environmental drivers of metabolic heterogeneity in clonal microbial populations. Curr. Opin. Biotechnol., 62, 202–211.

3. Erickson, D.W., Schink, S.J., Patsalo, V., Williamson, J.R., Gerland, U. and Hwa, T. (2017) A global resource allocation strategy governs growth transition kinetics of Escherichia coli. Nature, 551, 119–123.

4. Rugbjerg, P. and Sommer, M.O.A. (2019) Overcoming genetic heterogeneity in industrial fermentations. Nat. Biotechnol., 37, 869–876.

5. Okano, H., Hermsen, R., Kochanowski, K. and Hwa, T. (2020) Regulation underlying hierarchical and simultaneous utilization of carbon substrates by flux sensors in Escherichia coli. Nat. Microbiol., 5, 206–215.

6. Delvigne, F., Zune, Q., Lara, A.R., Al-Soud, W. and Sørensen, S.J. (2014) Metabolic variability in bioprocessing: Implications of microbial phenotypic heterogeneity. Trends Biotechnol., 32, 608–616.

7. Tasdogan, A., Faubert, B., Ramesh, V., Ubellacker, J.M., Shen, B., Solmonson, A., Murphy, M.M., Gu, Z., Gu, W., Martin, M., et al. (2020) Metabolic heterogeneity confers differences in melanoma metastatic potential. Nature, 577, 115–120.

8. Takhaveev, V. and Heinemann, M. (2018) Metabolic heterogeneity in clonal microbial populations. Curr. Opin. Microbiol., 45, 30–38.

9. Huberts, D.H.E.W., Niebel, B. and Heinemann, M. (2012) A flux-sensing mechanism could regulate the switch between respiration and fermentation. FEMS Yeast Res., 12, 118–28.

10. Litsios, A., Ortega, A.D., Wit, E.C. and Heinemann, M. (2017) Metabolic-flux dependent regulation of microbial physiology. Curr. Opin. Microbiol., 42, 71–78.

11. Deutscher, J., Küster, E., Bergstedt, U., Charrier, V. and Hillen, W. (1995) Protein kinase-dependent HPr/CcpA interaction links glycolytic activity to carbon catabolite repression in Grapositive bacteria. Mol. Microbiol., 15, 1049–1053.

12. Doan, T. and Aymerich, S. (2003) Regulation of the central glycolytic genes in Bacillus subtilis: Binding of the repressor CggR to its single DNA target sequence is modulated by fructose-1,6-bisphosphate. Mol. Microbiol., 47, 1709–1721.

13. Peeters, K., Van Leemputte, F., Fischer, B., Bonini, B.M., Quezada, H., Tsytlonok, M., Haesen, D., Vanthienen, W., Bernardes, N., Gonzalez-Blas, C.B., et al. (2017) Fructose-1,6-bisphosphate couples glycolytic flux to activation of Ras. Nat. Commun., 8.

14. Zhang, C.-S., Hawley, S.A., Zong, Y., Li, M., Wang, Z., Gray, A., Ma, T., Cui, J., Feng, J.-W., Zhu, M., et al. (2017) Fructose-1,6-bisphosphate and aldolase mediate glucose sensing by AMPK. Nature, 548, 112–116.

15. Hallberg, Z.F., Su, Y., Kitto, R.Z. and Hammond, M.C. (2017) Engineering and In Vivo Applications of Riboswitches. Annu. Rev. Biochem., 86, 515–539.

16. Schmidt, C.M. and Smolke, C.D. (2019) RNA Switches for Synthetic Biology. Cold Spring Harb. Perspect. Biol., 11.

17. Suess, B., Hanson, S., Berens, C., Fink, B., Schroeder, R. and Hillen, W. (2003) Conditional gene expression by controlling translation with tetracycline-binding aptamers. Nucleic Acids Res., 31, 1853–8.

18. Weigand, J.E., Sanchez, M., Gunnesch, E., Zeiher, S., Schroeder, R. and Suess, B. (2008) Screening for engineered neomycin riboswitches that control translation initiation. RNA, 10.1261/rna.772408.4.

19. Berschneider, B., Wieland, M., Rubini, M. and Hartig, J.S. (2009) Small-molecule-dependent regulation of transfer RNA in bacteria. Angew. Chemie – Int. Ed., 48, 7564–7567.

20. Babiskin, A.H. and Smolke, C.D. (2011) Engineering ligand-responsive RNA controllers in yeast through the assembly of RNase III tuning modules. Nucleic Acids Res., 39, 5299–5311.

21. Win, M.N. and Smolke, C.D. (2007) A modular and extensible RNA-based gene-regulatory platform for engineering cellular function. Proc. Natl. Acad. Sci. U. S. A., 104, 14283–14288.

22. Wong, R.S., Chen, Y.Y. and Smolke, C.D. (2018) Regulation of T cell proliferation with drug-responsive microRNA switches. Nucleic Acids Res., 46, 1541–1552.

23. Qi, L., Lucks, J.B., Liu, C.C., Mutalik, V.K. and Arkin, A.P. (2012) Engineering naturally occurring trans-acting non-coding RNAs to sense molecular signals. Nucleic Acids Res., 40, 5775–86.

24. Paige, J.S., Nguyen-Duc, T., Song, W. and Jaffrey, S.R. (2012) Fluorescence imaging of cellular metabolites with RNA. Science, 335, 1194.

25. Kellenberger, C.A., Wilson, S.C., Sales-Lee, J. and Hammond, M.C. (2013) RNA-Based Fluorescent Biosensors for Live Cell Imaging of Second Messengers Cyclic di-GMP and Cyclic AMP-GMP. J. Am. Chem. Soc., 135, 4906–4909.

26. Bose, D., Su, Y., Marcus, A., Raulet, D.H. and Hammond, M.C. (2016) An RNA-Based Fluorescent Biosensor for High-Throughput Analysis of the cGAS-cGAMP-STING Pathway. Cell Chem. Biol., 23, 1539–1549.

27. Breaker, R.R. (2004) Natural and engineered nucleic acids as tools to explore biology. Nature, 432, 838–845.

28. McKeague, M., Wong, R.S. and Smolke, C.D. (2016) Opportunities in the design and application of RNA for gene expression control. Nucleic Acids Res., 10.1093/nar/gkw151.

29. Groher, F., Bofill-bosch, C., Schneider, C., Braun, J., Jager, S., Geißler, K., Hamacher, K. and Suess, B. (2018) Riboswitching with ciprofloxacin –– development and characterization of a novel RNA regulator. 46, 2121–2132.

30. Porter, E.B., Polaski, J.T., Morck, M.M. and Batey, R.T. (2017) Recurrent RNA motifs as scaffolds for genetically encodable small-molecule biosensors. Nat. Chem. Biol., 13, 295–301.

31. Desai, S.K. and Gallivan, J.P. (2004) Genetic screens and selections for small molecules based on a synthetic riboswitch that activates protein translation. J. Am. Chem. Soc., 126, 13247–54.

32. Lynch, S.A., Desai, S.K., Sajja, H.K. and Gallivan, J.P. (2007) A high-throughput screen for synthetic riboswitches reveals mechanistic insights into their function. Chem. Biol., 14, 173–84.

33. Fowler, C.C., Brown, E.D. and Li, Y. (2008) A FACS-Based Approach to Engineering Artificial Riboswitches. ChemBioChem, 9, 1906–1911.

34. Liang, J.C., Chang, A.L., Kennedy, A.B. and Smolke, C.D. (2012) A high-throughput, quantitative cell-based screen for efficient tailoring of RNA device activity. Nucleic Acids Res., 40, e154.

35. Townshend, B., Kennedy, A.B., Xiang, J.S. and Smolke, C.D. (2015) High-throughput cellular RNA device engineering. Nat. Methods, 12, 989–994.

36. Klauser, B., Atanasov, J., Siewert, L.K. and Hartig, J.S. (2015) Ribozyme-Based Aminoglycoside Switches of Gene Expression Engineered by Genetic Selection in S. cerevisiae. ACS Synth. Biol., 4, 516–525.

37. Xiang, J.S., Kaplan, M., Dykstra, P., Hinks, M., McKeague, M. and Smolke, C.D. (2019) Massively parallel RNA device engineering in mammalian cells with RNA-Seq. Nat. Commun., 10, 4327.

38. Bennett, B.D., Kimball, E.H., Gao, M., Osterhout, R., Van Dien, S.J. and Rabinowitz, J.D. (2009) Absolute metabolite concentrations and implied enzyme active site occupancy in Escherichia coli. Nat. Chem. Biol., 5, 593–599.

39. Ozalp, V.C., Pedersen, T.R., Nielsen, L.J. and Olsen, L.F. (2010) Time-resolved measurements of intracellular ATP in the yeast Saccharomyces cerevisiae using a new type of nanobiosensor. J. Biol. Chem., 285, 37579–88.

40. Long, Y., Pfeiffer, F., Mayer, G., Schrøder, T.D., Özalp, V.C. and Olsen, L.F. (2016) Selection of Aptamers for Metabolite Sensing and Construction of Optical Nanosensors. Methods Mol. Biol., 1380, 3–19.

41. Bley Folly, B., Ortega, A.D., Hubmann, G., Bonsing-Vedelaar, S., Wijma, H.J., van der Meulen, P., Milias-Argeitis, A. and Heinemann, M. (2018) Assessment of the interaction between the flux-signaling metabolite fructose-1,6-bisphosphate and the bacterial transcription factors CggR and Cra. Mol. Microbiol., 109, 278–290.

42. Wilkinson, K.A., Merino, E.J. and Weeks, K.M. (2006) Selective 2’-hydroxyl acylation analyzed by primer extension (SHAPE): quantitative RNA structure analysis at single nucleotide resolution. Nat. Protoc., 1, 1610–6.

43. Reuter, J.S., Mathews, D.H., Miranti, C.K., Beck, A., Polyak, K., Toker, A., Sethi, T., Bourget, L., Lamoureux, L. and Lo, R. (2010) RNAstructure: software for RNA secondary structure prediction and analysis. BMC Bioinformatics, 11, 129.

44. Litsios, A., Huberts, D.H.E.W., Guerra, P., Schmidt, A., Buczak, K., Papagiannakis, A., Rovetta, M., Hekelaar, J., Hubmann, G., Exterkate, M., et al. (2019) Differential scaling between G1 protein production and cell size dynamics promotes commitment to the cell division cycle in budding yeast. Nat. Cell Biol., In Press.

45. Elbing, K., Larsson, C., Bill, R.M., Albers, E., Snoep, J.L., Boles, E., Hohmann, S. and Gustafsson, L. (2004) Role of hexose transport in control of glycolytic flux in Saccharomyces cerevisiae. Appl. Environ. Microbiol., 70, 5323–30.

46. Kochanowski, K, Volkmer, B., Gerosa, L, Haverkorn van Rijsewijk, B.R., Schmidt, A, Heinemann, M., Kochanowski, K., Volkmer, B., Gerosa, L., Haverkorn van Rijsewijk, B.R., Schmidt, A., Heinemann, M., Kochanowski, K, Volkmer, B., et al. (2013) Functioning of a metabolic flux sensor in Escherichia coli. Proc. Natl. Acad. Sci. U. S. A., 110, 1130–1135.

47. Hackett, S.R., Zanotelli, V.R.T., Xu, W., Goya, J., Park, J.O., Perlman, D.H., Gibney, P.A., Botstein, D., Storey, J.D. and Rabinowitz, J.D. (2016) Systems-level analysis of mechanisms regulating yeast metabolic flux. Science (80-.)., 354, aaf2786–aaf2786.

48. Schmidt, A.M. (2014) Flux-signaling and flux-dependent regulation in Saccharomyces cerevisae. 10.3929/ethz-a-010160040.

49. Zubradt, M., Gupta, P., Persad, S., Lambowitz, A.M., Weissman, J.S. and Rouskin, S. (2017) DMS-MaPseq for genome-wide or targeted RNA structure probing in vivo. Nat. Methods, 14, 75–82.

50. Homan, P.J., Favorov, O. V., Lavender, C.A., Kursun, O., Ge, X., Busan, S., Dokholyan, N. V. and Weeks, K.M. (2014) Single-molecule correlated chemical probing of RNA. Proc. Natl. Acad. Sci. U. S. A., 111, 13858–13863.

51. Lee, S.S., Avalos Vizcarra, I., Huberts, D.H.E.W., Lee, L.P. and Heinemann, M. (2012) Whole lifespan microscopic observation of budding yeast aging through a microfluidic dissection platform. Proc. Natl. Acad. Sci. U. S. A., 109, 4916–20.

52. Huberts, D.H.E.W., Sik Lee, S., Gonzáles, J., Janssens, G.E., Vizcarra, I.A. and Heinemann, M. (2013) Construction and use of a microfluidic dissection platform for long-term imaging of cellular processes in budding yeast. Nat. Protoc., 8, 1019–27.

53. Canelas, A.B.A.B., Ras, C., ten Pierick, A., van Gulik, W.M. and Heijnen, J.J. (2011) An in vivo data-driven framework for classification and quantification of enzyme kinetics and determination of apparent thermodynamic data. Metab. Eng., 13, 294–306.

54. Ames, T.D. and Breaker, R.R. (2011) Bacterial aptamers that selectively bind glutamine. RNA Biol., 8, 82–89.

55. Ren, A., Xue, Y., Peselis, A., Serganov, A., Al-Hashimi, H.M. and Patel, D.J. (2015) Structural and Dynamic Basis for Low-Affinity, High-Selectivity Binding of L-Glutamine by the Glutamine Riboswitch. Cell Rep., 13, 1800–1813.

56. Monteiro, F., Hubmann, G., Takhaveev, V., Vedelaar, S.R., Norder, J., Hekelaar, J., Saldida, J., Litsios, A., Wijma, H.J., Schmidt, A., et al. (2019) Measuring glycolytic flux in single yeast cells with an orthogonal synthetic biosensor. Mol. Syst. Biol., 15, 1–20.

57. Wieland, M. and Hartig, J.S. (2008) Improved aptazyme design and in vivo screening enable riboswitching in bacteria. Angew. Chemie – Int. Ed., 47, 2604–2607.

58. Ausländer, S., Ketzer, P. and Hartig, J.S. (2010) Aligand-dependent hammerhead ribozyme switch for controlling mammalian gene expression. Mol. Biosyst., 6, 807–814.

59. Kellenberger, C.A., Chen, C., Whiteley, A.T., Portnoy, D.A. and Hammond, M.C. (2015) RNA-Based Fluorescent Biosensors for Live Cell Imaging of Second Messenger Cyclic di-AMP. J. Am. Chem. Soc., 137, 6432–6435.

60. Yang, J., Seo, S.W., Jang, S., Shin, S.-I.I., Lim, C.H., Roh, T.-Y.Y. and Jung, G.Y. (2013) Synthetic RNA devices to expedite the evolution of metabolite-producing microbes. Nat. Commun., 4, 1413.

61. Klauser, B. and Hartig, J.S. (2013) An engineered small RNA-mediated genetic switch based on a ribozyme expression platform. Nucleic Acids Res., 41, 5542–52.

62. Bayer, T.S. and Smolke, C.D. (2005) Programmable ligand-controlled riboregulators of eukaryotic gene expression. Nat. Biotechnol., 23, 337–343.

63. Win, M.N. and Smolke, C.D. (2008) Higher-order cellular information processing with synthetic RNA Devices. Science (80-.)., 322, 456–460.

64. Boussebayle, A., Torka, D., Ollivaud, S., Braun, J., Bofill-Bosch, C., Dombrowski, M., Groher, F., Hamacher, K. and Suess, B. (2019) Next-level riboswitch development—implementation of Capture-SELEX facilitates identification of a new synthetic riboswitch. Nucleic Acids Res., 47, 4883–4895.

65. McGinnis, J.L., Duncan, C.D.S. and Weeks, K.M. (2009) High-throughput SHAPE and hydroxyl radical analysis of RNA structure and ribonucleoprotein assembly. 1st ed. Elsevier Inc.

66. Karabiber, F., McGinnis, J.L., Favorov, O. V and Weeks, K.M. (2013) QuShape: rapid, accurate, and best-practices quantification of nucleic acid probing information, resolved by capillary electrophoresis. RNA, 19, 63–73.

67. Verduyn, C., Postma, E., Scheffers, W.A. and Van Dijken, J.P. (1992) Effect of benzoic acid on metabolic fluxes in yeasts: a continuous-culture study on the regulation of respiration and alcoholic fermentation. Yeast, 8, 501–17.

68. Radzikowski, J.L., Vedelaar, S., Siegel, D., Ortega, Á.D., Schmidt, A. and Heinemann, M. (2016) Bacterial persistence is an active σS stress response to metabolic flux limitation. Mol. Syst. Biol., 12, 882.

69. Ferrezuelo, F., Colomina, N., Palmisano, A., Garí, E., Gallego, C., Csikász-Nagy, A. and Aldea, M. (2012) The critical size is set at a single-cell level by growth rate to attain homeostasis and adaptation. Nat. Commun., 3.

70. Inoue, H., Nojima, H. and Okayama, H. (1990) High efficiency transformation of Escherichia coli with plasmids. Gene, 96, 23–8.

71. Gietz, R.D. and Schiestl, R.H. (2007) Large-scale high-efficiency yeast transformation using the LiAc/SS carrier DNA/PEG method. Nat. Protoc., 2, 38–41.

72. Incarnato, D., Morandi, E., Simon, L.M. and Oliviero, S. (2018) RNA framework: An all-in-one toolkit for the analysis of RNA structures and post-transcriptional modifications. Nucleic Acids Res., 46, 1–13.

73. Langmead, B. and Salzberg, S.L. (2012) Fast gapped-read alignment with Bowtie 2. Nat. Methods, 9, 357–359.

74. Schindelin, J., Arganda-Carrera, I., Frise, E., Verena, K., Mark, L., Tobias, P., Stephan, P., Curtis, R., Stephan, S., Benjamin, S., et al. (2009) Fiji – an Open platform for biological image analysis. Nat. Methods, 9, 241.

75. van der Walt, S., Schönberger, J.L., Nunez-Iglesias, J., Boulogne, F., Warner, J.D., Yager, N., Gouillart, E. and Yu, T. (2014) scikit-image: image processing in Python. PeerJ, 2, e453.

