## Supplementary Files for "A synthetic RNA-based biosensor for fructose-1,6-bisphosphate that reports glycolytic flux"

### CONTENTS

#### SUPPLEMENTARY NOTES


Supplementary Note 1 | Analysis of the GFP/mCherry-readout span and resolution in the HHRz-controlled *in vivo* reporter system
Supplementary Note 2 | Defining the screening sequence space within the C45-HHRz input library
Supplementary Note 3 | Hit selection in the high-throughput screening of C45-HHRz clones reporting intracellular FBP
Supplementary Note 4 | Comparison between the novel RNA-based biosensor and the previously reported transcription-factor-based one


#### SUPPLEMENTARY FIGURES


Supplementary Figure S1 | Affinity and specificity of different clones obtained by SELEX
Supplementary Figure S2 | Secondary structure models of RNA constructs used in binding analyses
Supplementary Figure S3 | Deletion of U23 bulge in C45 does not alter the aptamer's selectivity
Supplementary Figure S4 | An *in vivo* reporter system that couples RNA conformational changes with a measurable quantitative fluorescent readout
Supplementary Figure S5 | Design of C45-HHRz RNA device libraries
Supplementary Figure S6 | Molecular exploration of the diversity and composition of the pre-selected C45-HHRz RNA device library with a functional ribozyme moiety
Supplementary Figure S7 | Criteria to set the gate bounds for sorting C45-HHRz RNA device library
Supplementary Figure S8 | A robust assignment of GFP/mCherry ratio to each clone in C45-HHRz library is based on its relative frequency in sorted cell populations
Supplementary Figure S9 | The sensor 2_6 has two conformations inside the cell whereas the mutant 2_6m folds in only one conformation


#### SUPPLEMENTARY TABLES


Supplementary Table S1 | The sequences of the clones retrieved from SELEX round 13
Supplementary Table S2 | Formal concepts and measures used in the analysis of cell sorting and NGS data
Supplementary Table S3 | Regular expressions (search patterns) to classify the sequences obtained in NGS
Supplementary Table S4 | The frequency of unique occurrences and the coverage fraction of undefined sequences
Supplementary Table S5 | Cross-contamination rate
Supplementary Table S6 | NGS reads per unique plasmid molecule
Supplementary Table S7 | Oligonucleotides used in this study

**SUPPLEMENTARY NOTES**

**Supplementary Note 1. Analysis of the GFP/mCherry-readout span and resolution in the HHRz-controlled *in vivo* reporter system**

For any sensor or reporter system, the span corresponds to the total range of the readout values it can deliver (*i.e*. difference between the maximum and minimum readout values), whereas the resolution is the minimal change that can be detected. To define the GFP/mCherry-readout span of the HHRz-regulated reporter system, we analyzed a series of mutants in the HHRz sequence. First, we took the wild-type sequence of the tobacco ringspot virus (sTRSV) HHRz and an inactive mutant (a substitution of the active site) as references of the minimum and maximum readout values (corresponding to the maximal and no self-cleaving activity, respectively) (HHRz and CTL in Supplementary Fig. S4B-C). Here, we observed that the system exhibits a full-scale GFP/mCherry output that spans over two-logs (Supp. Fig. S4D). Then, since the activity of HHRz has been shown to depend on tertiary structure-stabilizing interactions between the loops of stem-loops I and II, modifications in the sequence of these loops provided a set of mutants displaying intermediate GFP/mCherry levels, thereby representing a linear gradation of the readout values yielded by this reporter system (M1-M7, Supplementary Fig. S4B, D) [1]. Considering the variable dispersion of single-cell distributions of the GFP/mCherry readout along the full scale (different sizes of the boxes in Supplementary Fig. S4D), we concluded that the HHRz-controlled *in vivo* reporter system would allow the separation of maximally eight clonal populations on the basis of their GFP/mCherry ratios, without significant overlap.

**Supplementary Note 2. Defining the screening sequence space within the C45-HHRz input library**

To get a deep insight into the clonal composition of the C45-HHRz RNA device library, we explored the diversity and frequency of the clones by targeted next generation sequencing (NGS). We overlapped the sets of unique sequences found in every sample and determined that the number of unique sequences appearing in both biological replicates of the same strain and in both strains was 57,985 (Set IX, Supplementary Fig. S6A). This number is higher than the maximal number of unique sequences that could theoretically exist by design (that is, $\sum_i 4^{n_i}$, with $i$ being one of the six strategies to graft C45 into HHRz and *n* being the number of nucleotides within the randomized region: $3 \cdot 4^4 + 3 \cdot 4^7 = 49{,}920$). In addition, the C45-HHRz library is a pre-selected subpopulation enriched in C45-HHRz RNA devices with a functional HHRz moiety, which represented less than 10% of the original Lib-sI and sII collections (Fig. 3). Therefore, the diversity of the C45-HHRz library would be expected much lower.

To establish a quality control that allowed us to filter out sequences that do not correspond to actual cell clones present in the library, we determined a coverage threshold below which sequences would be considered noise and thus not suitable for screening. To this aim, we sought to identify artefactual sequences and check their coverage. Artifacts could have arisen from sequencing errors and spurious PCR products introduced during NGS library preparation. Sequencing and polymerase error rate is a consistent and repetitive source of noise while polymerase errors in initial rounds could lead to excessive amplification of particular sequences. However, the probability that the same mutation appeared in different samples is very low and, consequently, overlapping the sequence repertoire in biological replicates would eliminate both sources of noise.

We transformed our original libraries Lib-sI and sII in the isogenic *TM6** and wild-type strains, pre-selected RNA devices with a functional ribozyme, pooled the I and II libraries to get a unique C45-HHRz input sample in *TM6** and wild-type backgrounds that was eventually analyzed by targeted NGS in two independent experiments. The overlap of sequences among multiple samples (between biological replicates of the C45-HHRz library in *TM6** and wild-type backgrounds, and between common sets from each strain) resulted in different sets of unique sequences (Supplementary Fig. S6A). Sequence coverage in non-overlapped sets of sequences was notably skewed to minimally represented sequences, where more than 80% of the sequences were covered only by two reads (Sets I, II, IV and V; Supplementary Fig. S6B). Conversely, the sequences found in all the four samples (2 replicates × 2 strains) are supported by two reads in approximately 20% of cases (set IX; Supplementary Fig. S6B).

The sequences covered by four or more reads in sets VII and VIII correspond to real clones present in C45-HHRz libraries exclusively in one strain (Supplementary Fig. S6B). To generate the cell libraries, we used the same DNA libraries and collected the same number of transformants for both wild-type and *TM6** strains, suggesting that slight differences in GFP/mCherry ratio might have introduced variability in the enrichment of functional C45-HHRz devices (Fig. 3E). All in all, this analysis indicates that sequences covered by 2 reads mostly correspond to artefacts and, thus, we established the coverage threshold of more than two reads in our analysis pipeline.

By applying the coverage threshold of more than 2 reads in the overlapped set of sequences (set IX), we obtained the final set (set X) with the diversity of 28,708 clones (Supplementary Fig. S6C), which is likely a fair reflection of the actual clones present in C45-HHRz RNA device library cell population and represents the sequence space to be screened (Fig. 4). Having applied two criteria to select the sequences for analysis (that is, multiple sample overlap and sequence coverage threshold), we explain the diversity beyond theoretically possible combinations (*e.g.* sequences with differences in the HHRz constant regions) by variations introduced during DNA library preparation prior to transformation or by spontaneous mutations in yeast cells.

**Supplementary Note 3. Hit selection in the high-throughput screening of C45-HHRz clones reporting intracellular FBP**

To maximize the likelihood that a sequence observed in NGS represents a clone that existed in the cell populations, we consider the sequence in the analysis only if it is supported by more than 2 reads in each of the four sorting experiments of *TM6** strain (2 replicates × 2 carbon sources). As a sorting experiment was followed by NGS in six samples corresponding to the bins of GFP/mCherry ratio, we calculated the total coverage of the sequence in these samples and checked whether it is bigger than 2 reads. After this pre-selection of sequences in the *TM6** strain, we ended up with 15,031 clones.

To assess the technical quality of the sorting experiments, we evaluated the preservation of the clonal content during sorting and the consistency of the clone response between the replicates. We observed a high degree of correlation between the clone frequency in the input population and the reconstructed frequency after sorting (Supplementary Figure S8A). Similarly, there is a strong correlation between the replicates regarding the clone readout $\mu_i$ potentially reflecting intracellular FBP (Supplementary Figure S8B). Thus, the clonal content in the sorted populations largely reflects that of the input populations, and the clone readout reproduces robustly in replicate sorting.

To identify clones that differentially responded in *TM6** cells growing on maltose and on glucose, we used the measure $D_i$ (Supplementary Table S2). We selected clones with the extreme values of the measure $D_i$ that are located beyond the segment $[Q1 - 5 \times IQR, Q3 + 5 \times IQR]$, where $Q1$ and $Q3$ are the lower and upper quartiles, and $IQR$ is the interquartile range (Figure 5A). To exclude clones that end up in the selection due to technical noise, we further shrank the selection to those clones that have the extreme values of $D_i$ also in the other replicate sorting experiment. After this screening, we arrive to 63 clones distinctly responding in *TM6** cells consuming maltose or glucose.

To check whether the selected clones have similar readouts in the conditions of comparable intracellular FBP, we studied the $D_i$ values of the clones expressed in the wild-type cells growing on maltose and glucose. We considered a clone not distinguishing the carbon sources if its $D_i$ value is located within the $IQR$ of the clones supported by more than 2 reads in each of the four sorting experiments (Figure 5B). To find in the wild-type strain the maximal number of clones that had the extreme responses in *TM6**, we decrease the required support of these clones to at least 2 reads in each of the four sorting experiments. To exclude the technical noise, we look for the clones that are within the $IQR$ in both replicate sorting experiments of the wild-type strain. Thus, we have checked that 11 clones out of the 63 responding to

FBP difference have identical readouts in the conditions of similar FBP. These 11 clones represent the hits of the high-throughput screening.

**Supplementary note 4. Comparison between the novel RNA-based biosensor and the previously reported transcription-factor-based one**

Here, we compare the developed RNA-based biosensor (2_6) with the previously reported transcription-factor- (TF-) based biosensor [2], which comes in two versions with different response curves (denoted as 'Wildtype' and 'R250A').

**Sensor resolution**

We determined the sensor resolution as the smallest difference between the intracellular FBP concentrations under two studied metabolic conditions, in which the respective readouts of the sensor do not overlap:

$$min_{a,b:a\neq b}\{\,|FBP_a - FBP_b|\}\ such\ that\ M(R)_a \pm SD(R)_a \cap M(R)_b \pm SD(R)_b\ =\ \emptyset,$$

where $a$ and $b$ are two of the studied metabolic conditions $\boldsymbol{C}$, $FBP_a$ is the intracellular FBP concentration in the condition $a$, $M(R)_a$ and $SD(R)_a$ are the mean and standard deviation of median readouts across replicate cellular cultivations under the condition $a$. As the studied metabolic conditions do not fully overlap between the two works, we considered different sets of metabolic conditions to estimate the biosensor resolutions.

First, we considered the metabolic conditions that were studied in both works, namely, the strain TM6 on Glucose, TM6 on Maltose and the wildtype (WT) on Glucose. Given these metabolic conditions, the resolution of both biosensors is calculated as the difference between the intracellular FBP concentrations in TM6 on Glucose and TM6 on Maltose, yielding for both biosensors the same resolution value in the range from 3.11±1.13 to 5.33±1.01 mM, according to the FBP measurements performed in two works (Table SN4.1).

Second, we considered all metabolic conditions studied in each work. Here, we found that the RNA-based biosensor can even have a resolution of 0.25±0.06 mM, distinguishing the metabolic conditions of TM4 on Glucose and TM6 on Glucose (Table SN4.1).

**Table SN4.1. The resolution of the RNA- and TF-based biosensors is estimated not to differ if the same metabolic conditions studied together in both works are considered. However, if the set of compared metabolic conditions is extended to all studied ones, we find that the RNA-based biosensor can have a resolution of at least 0.25 mM.** A resolution value is the difference between mean FBP concentrations under two indicated conditions, with the error being the square root of the sum of variances in these conditions.

| Data considered | Biosensor type | Name | Two distinguished conditions with the most similar FBP, readout mean ± SD | Source of sensor readout data | Resolution | Source of FBP data |
|---|---|---|---|---|---|---|
| Metabolic conditions present in both works | RNA-based | 2_6 | *TM6 Glucose*, 1.839±0.112 vs *TM6 Maltose*, 1.558±0.137 | 3 replicates contributed to Figure 6A, 2_6 [this study] | 3.11±1.13 mM ***** | Figure 3C [this study] |
| | TF-based | R250A | *TM6 Glucose*, 0.727±0.027 vs *TM6 Maltose*, 2.034±0.389 | Figure 6D [2] | 5.33±1.01 mM ***** | Figure 6D [2] |
| | | Wild-type | *TM6 Glucose*, 1.177±0.017 vs *TM6 Maltose*, 2.752±0.067 | Figure 6D [2] | 5.33±1.01 mM ***** | Figure 6D [2] |
| All metabolic conditions | RNA-based | 2_6 | TM4 Glucose, 1.463±0.013 vs TM6 Glucose, 1.531±0.018 | Figure 6D [this study] | 0.25±0.06 mM | Figure 6D [this study] |
| | TF-based | R250A | *WT Galactose*, 0.743±0.074 vs *TM6 Maltose*, 2.034±0.389 | Figure 6D [2] | 4.21±1.04 mM | Figure 6D [2] |
| | | Wild-type | *TM6 Glucose*, 1.177±0.017 vs *WT Galactose*, 1.9±0.125 | Figure 6D [2] | 1.12±0.31 mM | Figure 6D [2] |

*****Although the values are different due to experimental variability of FBP measurements, the same two metabolic conditions are compared for both biosensors, therefore, their resolution should be considered the same.

#### Full-scale output (maximal fold change)

Full-scale output (maximal fold change) is calculated as the difference between the maximal and minimal sensor readouts related to the minimal readout:

$$100\% \cdot (M(R)_b - M(R)_a)/M(R)_a \ such\ that\ argmin_x\{M(R)_x, x \in \boldsymbol{C}\} = a,\ argmax_x\{M(R)_x, x \in \boldsymbol{C}\} = b.$$

**Table SN4.2. The full-scale output of the TF-based biosensor is higher than that of the RNA-based one.**

| Biosensor type | Name | Full-scale output | Conditions with minimal and maximal readouts | Source |
|---|---|---|---|---|
| RNA-based | 2_6 | 53% | WT Glucose vs TM6 Glucose | Figure 6D [this study] |
| TF-based | R250A | 252% | TM6 Glucose vs WT Glucose | Figure 6D [2] |
| | Wildtype | 149% | TM6 Glucose vs WT Glucose | Figure 6D [2] |

Thus, the TF-based biosensor outperforms the RNA-based one in terms of maximal fold change.

**Bibliography**

1. Townshend, B., Kennedy, A. B., Xiang, J. S. & Smolke, C. D. High-throughput cellular RNA device engineering. *Nat. Methods* **12**, 989–994 (2015).

2. Monteiro, F. *et al.* Measuring glycolytic flux in single yeast cells with an orthogonal synthetic biosensor. *Mol. Syst. Biol.* **15**, 1–20 (2019).

**SUPPLEMENTARY FIGURES**

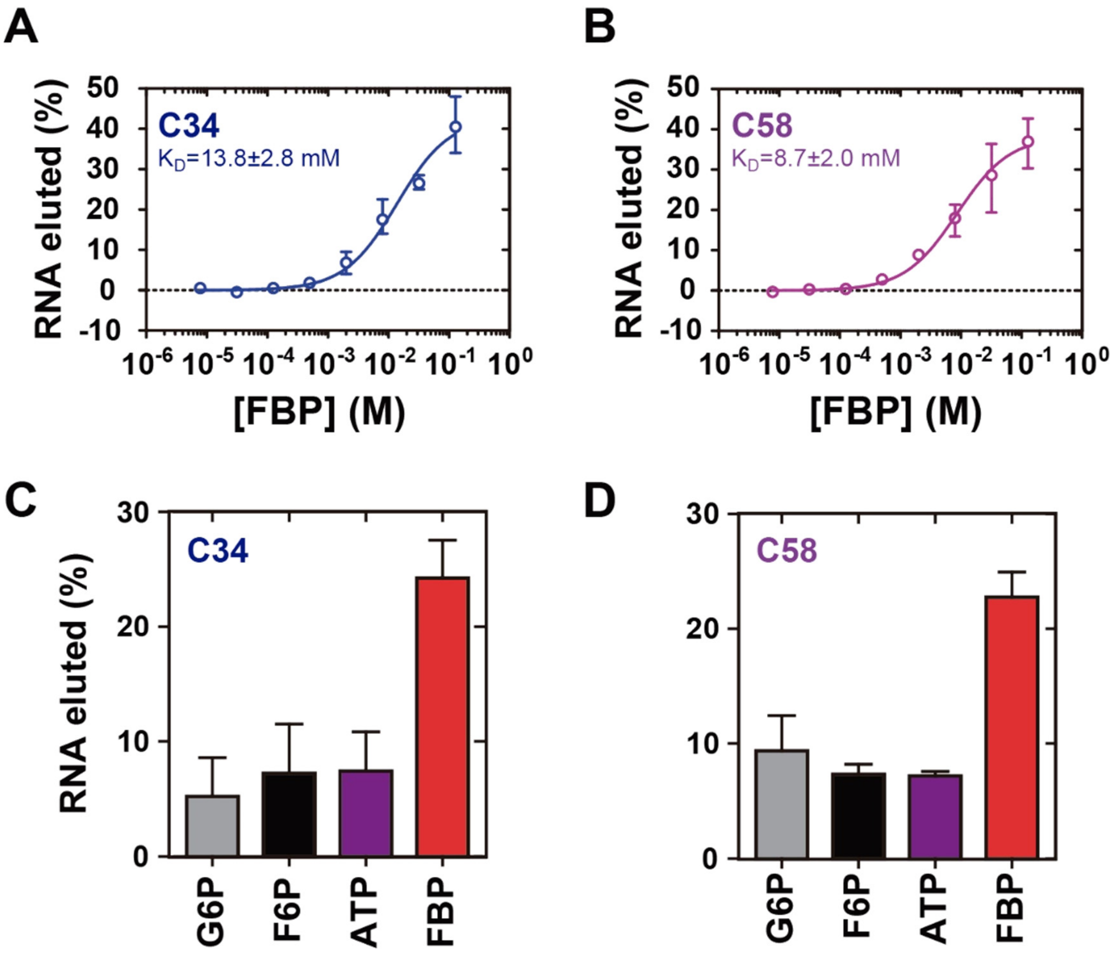


**Supplementary Figure S1. Affinity and specificity of different clones obtained by SELEX. A-B.** Binding assays of fluorescently-labeled (CAL Fluor Red 610) RNA clones C34 **(A)** and C58 **(B)** with FBP immobilized to sepharose. Labeled RNA was first incubated with FBP immobilized to sepharose, using as many samples as concentration points that were to be tested, the resin samples were washed, and then the RNA was eluted from the column with different concentrations of soluble FBP and quantified. Apparent dissociation constants ($K_D$) were estimated using a non-linear regression model assuming one binding site and specific binding with Hill slope. **C-D.** Specificity assay developed as in A but where a unique 10 mM solution of glucose-6-phosphate (G6P), fructose-6-phosphate (F6P), adenosine-triphosphate (ATP), or fructose-1,6-bisphosphate was used to release RNA from FBP-column. RNA clones C34 (**C**) and C58 (**D**) used in the specificity assay are indicated in the upper-left corner of the plots.

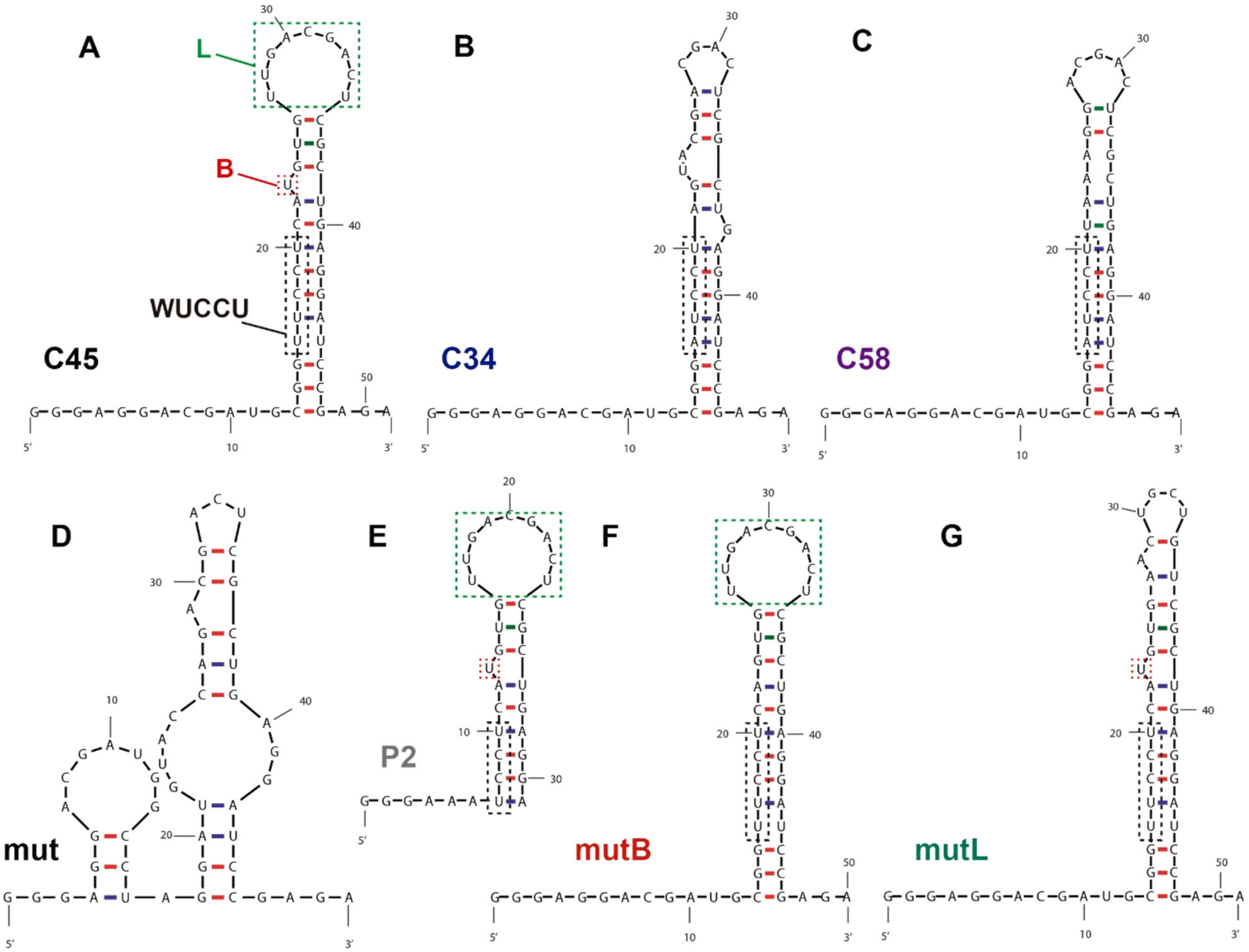


**Supplementary Figure S2. Secondary structure models of RNA constructs used in binding analyses**. **A-C.** Clones selected from the SELEX experiment. WUCCU, the enriched motif in the variable region of the clones sequenced in the SELEX experiment (W = A or U). L, loop. B, base bulge from stem. **D-E.** Mutants targeting the base of the conserved stem of C34 (**D**) and of C45 (**E**). In mutant *mut* C13-A21 was substituted by the complementary sequence; in mutant *P2* the 5' (G1 to G12) and 3' (A49 to A51) tails, and the base of the conserved stem (C13 to U16 and U45 to G48) was deleted. **F-G.** Mutants targeting the regions that underwent changes in SHAPE reactivity upon FBP binding: the U-bulge (**F**, *mutB*: U23 was deleted) and C45 loop (**G**, *mutL*: nucleotides U27 to U34 of the loop were substituted by the complementary sequence). RNA secondary structure models were retrieved using the energy minimization algorithm Mfold (M. Zuker, 2003, Nucleic Acids Res. 31 (13), 3406-15). Red and blue lines indicate *Watson and Crick* base pairs; green lines indicate Wobble or Hoogsteen base pairs.

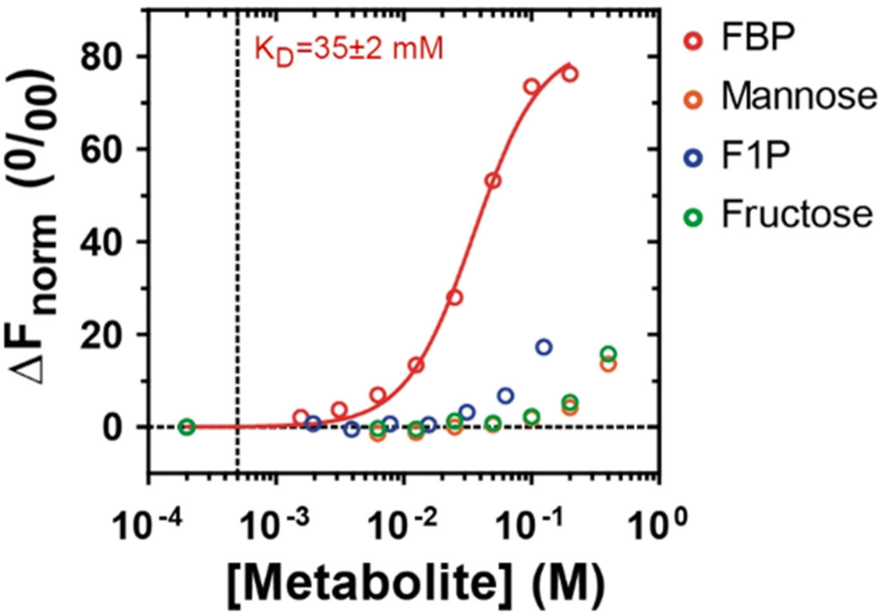


**Supplementary Figure S3. Deletion of U23 bulge in C45 does not alter the aptamer's selectivity**. MST analysis of binding of mutB to monosaccharides (fructose, mannose) and phosphorylated metabolites (F1P, fructose-1-phosphate; FBP, fructose-1,6-bisphosphate). Apparent dissociation constant ($K_D$) was estimated using a non-linear regression model assuming one binding-site and specific binding with Hill slope.

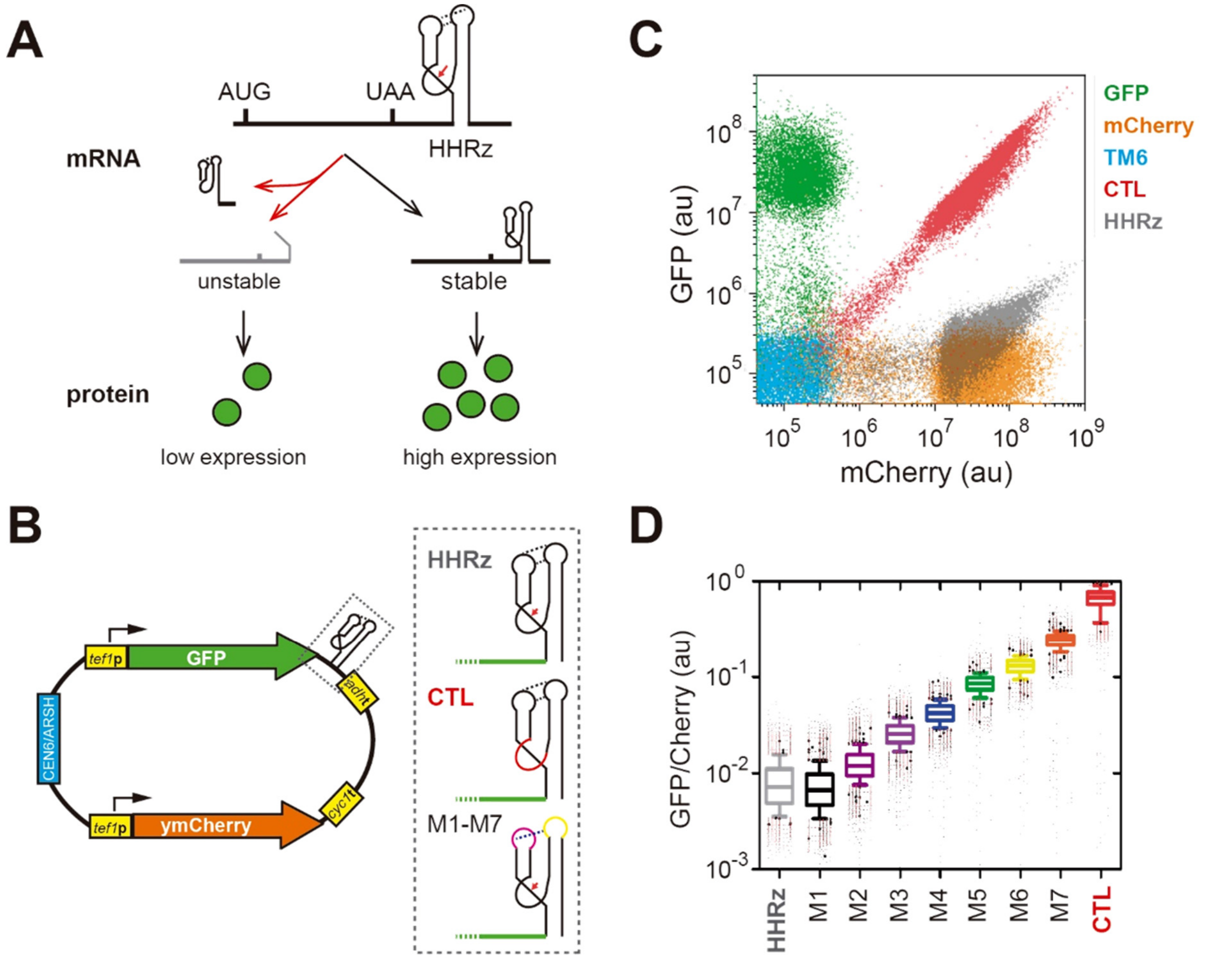


**Supplementary Figure S4. An *in vivo* reporter system that couples RNA conformational changes with a measurable quantitative fluorescent readout**. **A.** The tobacco ringspot virus (sTRSV) hammerhead ribozyme (HHRz) is inserted downstream the stop codon (UAA) of a reporter gene. Self-cleaving activity yields transcripts with shorter 3'UTR, which are less stable than transcripts with uncut 3'UTR. The red arrow denotes the cutting point. **B.** Schematic drawing of the map of the plasmid backbone (pCS1748, (Liang *et al.*, 2012)) used to clone the C45-HHRz library in the *in vivo* screening approach. *HHRz*, the tobacco ringspot virus (sTRSV) HHRz wild-type sequence; *CTL*, the inactive control ribozyme with a scrambled active site (red); *M1-M7*, a set of graded ribozymes with distinct sequences in loops I (magenta) and II (yellow) (Townshend *et al.*, 2015), which are predicted to display different long-distance tertiary structure interactions between loops I and II (dotted line), hence different self-cleaving activities that would differently affect GFP mRNA stability and protein expression. The green lines next to the HHRz schematic structures represent the rest of the GFP mRNA including the coding region. **C.** Overlay FACS dot plot showing GFP-mCherry expression pattern in controls used in this study. The dynamic range of this reporter system is covered by the space spanning from (and including) CTL (red cloud) to HHRz (grey cloud). *TM6*, the untransformed double-negative host strain; *GFP*, the GFP-only control plasmid pCS1585; *mCherry*, the mCherry-only control plasmid pCS1749. **D.** The box plot showing the distribution of the GFP/mCherry ratio in single cells determined by FACS. The boxes represent the interquartile range, and the whiskers are the 1 and 99 percentile bounds. Dots represent events lying outside the 1-99 percentile bounds of the distribution.

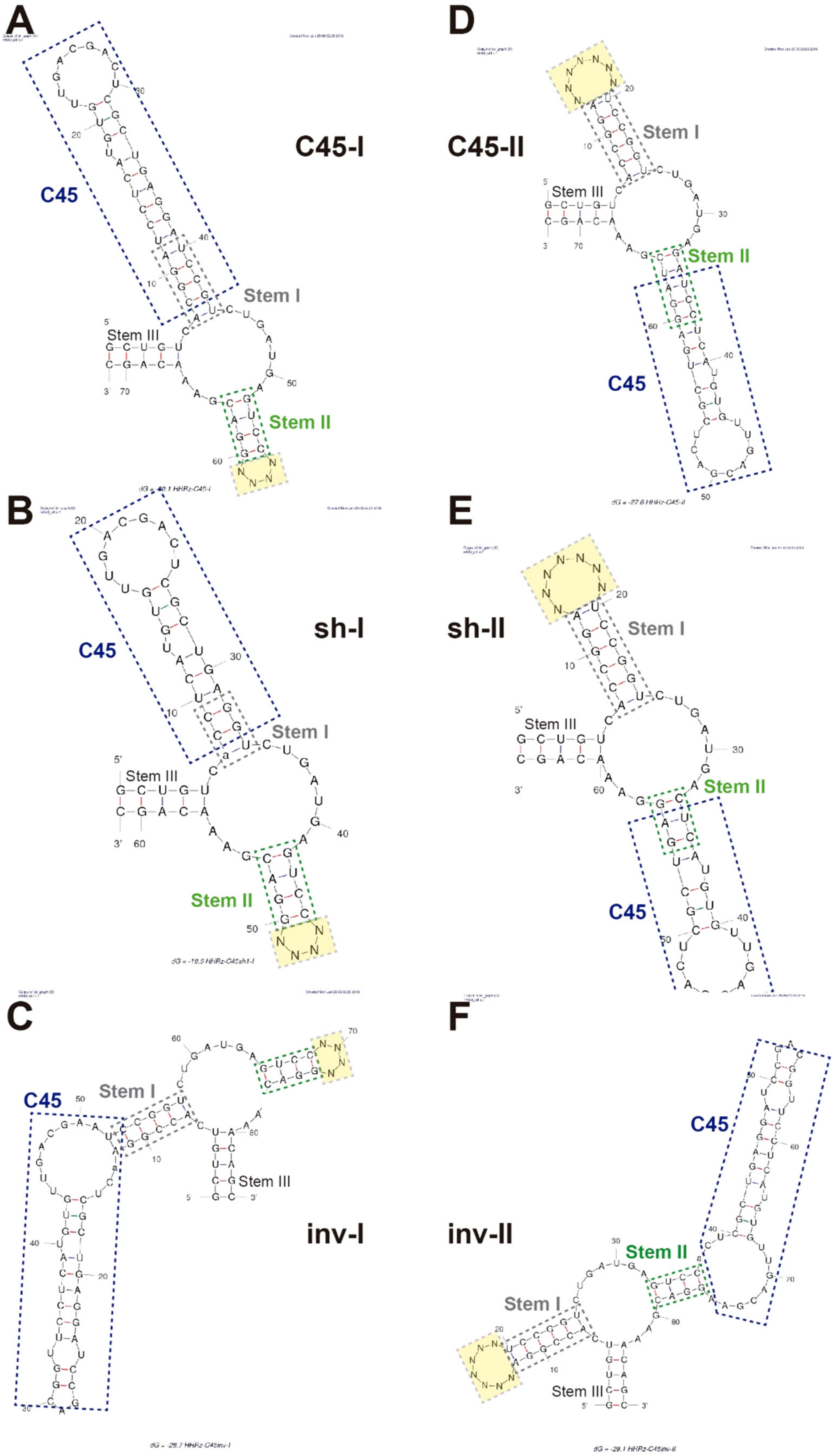
A
C45-I
C45
Stem I
Stem III
Stem II
B
sh-I
C45
Stem I
Stem III
Stem II
C
C45
Stem I
Stem III
inv-I
D
C45-II
Stem I
Stem III
Stem II
C45
E
sh-II
Stem I
Stem III
Stem II
C45
F
inv-II
C45
Stem II
Stem I
Stem III

**Supplementary Figure S5. Design of C45-HHRz RNA device libraries.** C45 was grafted into either of the HHRz stem-loops, I or II, using three different strategies to prevent biases caused by particular ways of combining the FBP aptamer and the ribozyme. For instance, the full-length C45 aptamer appended to the stem I could make the new C45-HHRz hybrid stem I too long, and hence the C45 loop would lye too far from the loop in the opposite stem-loop to establish tertiary structure interactions. Here, we show Mfold predictions of secondary structures adopted by C45-HHRz RNA devices in six libraries. C45 aptamer is grafted into either of the two stem-loops HHRz has (stem I, **A-C**; stem II, **D-F**) by removing the HHRz's corresponding loop and inserting C45 aptamer using three strategies. (i) C45 is placed next to the HHRz's remaining stem in the same (**A, D**) or (ii) inverted orientation (**C, F**); furthermore, (iii) a shortened version of C45 is merged with HHRz stem (**B, E**). C45, HHRz stems I and II in each representation are delimited with dotted-line boxes (blue, grey, green). Note that C45 stem and the HHRz's remainder stem sequences overlap in some constructs (**A-B, D-E**), and that the sequence of both stems is merged (might be slightly modified) at their edges in order to maintain the stem secondary structure and avoid increasing the length of the hybrid stem. Note that an A in C45 in inv libraries (**C**, **F**) lies out of C45 box because this nucleotide was added to close C45 stem and allow grafting C45 in the inverted orientation. The sequence of the opposite HHRz loop (*i.e.* loop I when C45 is grafted into stem II, and vice versa) is randomized (indicated with yellow boxes) to generate a library. From it, we select those clones that establish interactions between C45 and the opposite HHRz stem-loop, and therefore sustain HHRz activity.

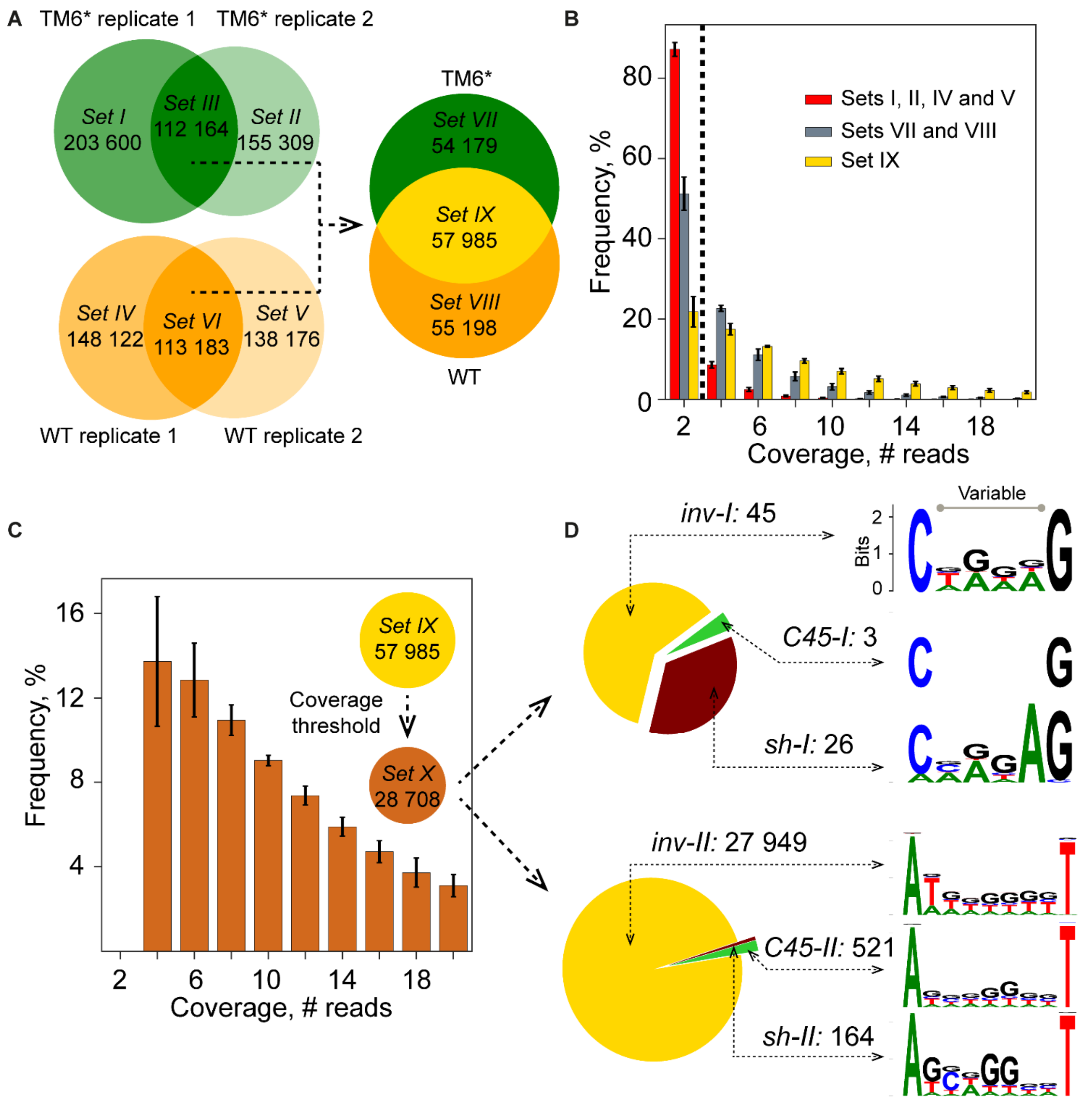


**Supplementary Figure S6. Molecular exploration of the diversity and composition of the pre-selected C45-HHRz RNA device library with a functional ribozyme moiety**. **A.** Venn diagrams showing the distribution of unique sequences in different C45-HHRz library input samples used in this study. The total number of unique sequences found in each sample is distributed among different sets. The unique sequences found in two biological replicates of the same strain (*TM6** or wild type) correspond to sets III and VI. The unique sequences found in both strains (sets III and VI) correspond to the set IX. The set sizes are presented on the segments of the Venn diagrams. **B.** The distribution of coverage among the sequences in the sets (I, II, IV and V) found within individual biological replicates only, in the sets (VII and VIII) found in two replicates of the same strain only and in the set (IX) found in all the four populations.

The bar length and the error whiskers shows the mean and two standard deviations of the frequency for a group of sequences (three groups are shown with different colors) in the four populations (samples). See Supplementary Note 2 discussing the diversity of the screening space and the introduction of the coverage threshold. **C.** The distribution of the coverage in the set (X) of the sequences that are supported by more than 2 reads in each of the four populations. In B-C, due to the paired-end sequencing, the number of reads is divisible by 2, with the smallest value being 2. **D.** Distribution of the three different C45 grafting strategies among the unique clones (Supplementary Fig. S5). The identifier of the grafting strategy (C45, inv, sh) and the receiving HHRz stem (I, II), and the number of corresponding clones are given above the arrows. The sequence logo shows the enrichment of nucleotides in the variable region and two adjacent positions. The information content shown on the y-axis is calculated using the alphabet size 5 (ATGC-).

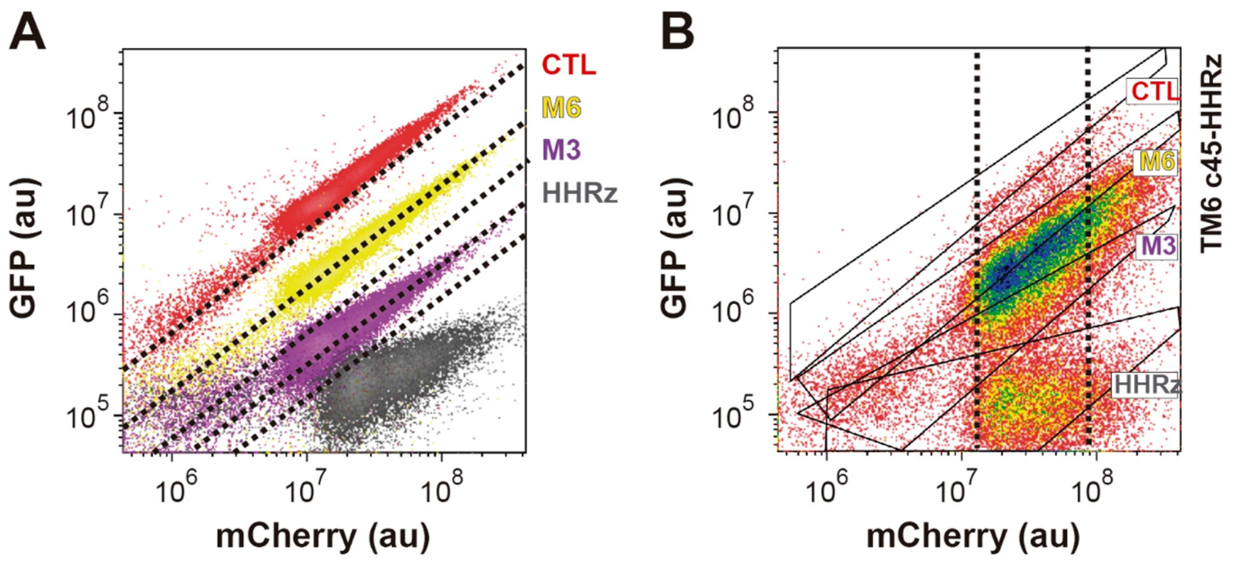


**Supplementary Figure S7. Criteria to set the gate bounds for sorting C45-HHRz RNA device library. A**. Setting gates based of GFP/mCherry ratio. Overlay GFP-mCherry dot plot of different HHRz clones used to delimit the full-scale output of the reporter system used in the screening (Supplementary Fig. S4). *HHRz*, tobacco ringspot virus (sTRSV) HHRz wild-type sequence; *CTL*, inactive control ribozyme with scrambled active site; *M3* and *M6*, graded ribozymes with distinct sequences in loops I and II displaying different self-cleaving activity. The upper and lower bounds (dotted lines) of the dynamic range of our reporter system correspond to HHRz (maximal activity) and CTL (no activity). The GFP-mCherry space spanning between these bounds is segmented into four bins whose size (width) is set according to the size that an individual clone has (and which represents roughly half of the intrinsic noise of GFP-mCherry expression). Note that bounds between bins are set so that they split the cell cloud of clones with intermediate GFP/mCherry ratio values (M3, M6) by half. **B**. Gating cells based on their mCherry value to reduce noise. GFP-mCherry dot plot of C45-HHRz RNA device library. Polygons depict GFP-mCherry cell clouds of clones used to set sorting gates based on GFP/mCherry ratio as shown in A. Note that, in case of the mCherry expression values lower than about $10^7$ but still considered positive for mCherry (Supplementary Fig. S4), cell populations displaying different GFP/mCherry ratios (poligons) get closer to each other and overlap. In contrast, cells with high mCherry values diverge, and differences in the ratio become less predictable. Gating cells with an intermediate mCherry expression value (between dotted lines) generates a more robust ratio output for cells in the populations.

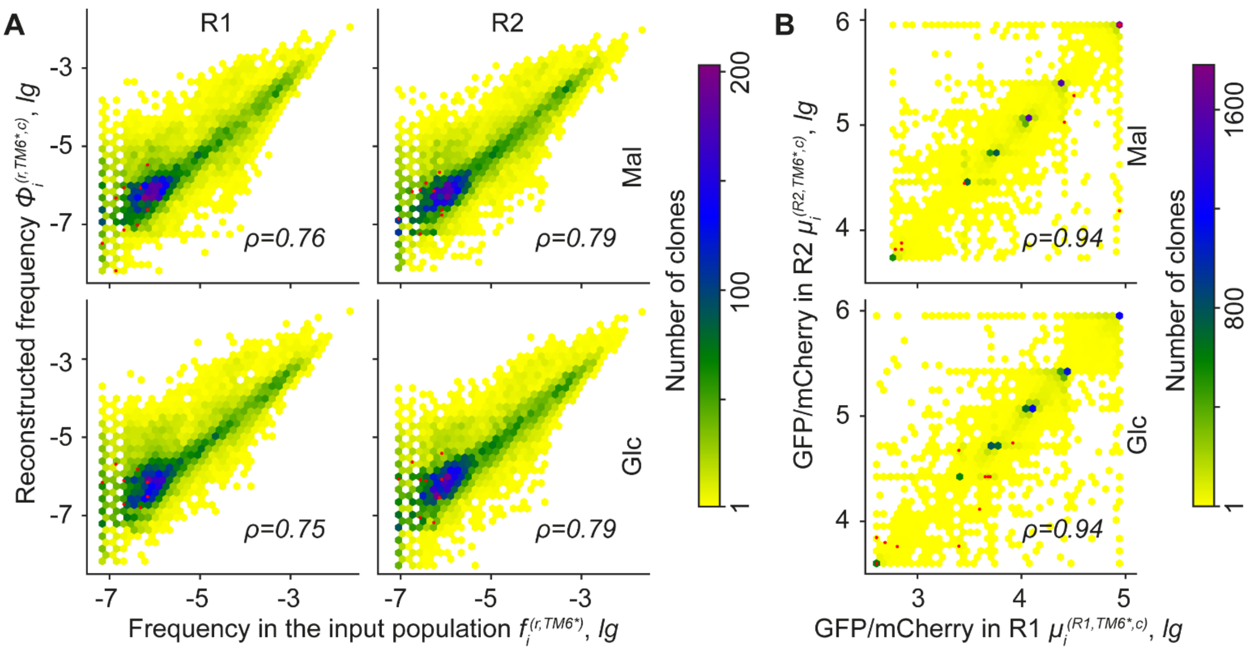


**Supplementary Figure S8. A robust assignment of GFP/mCherry ratio to each clone in C45-HHRz library is based on its relative frequency in sorted cell populations. A.** Correlation between the absolute frequency of a clone in the C45-HHRz library input population *before* sorting $f_i$ (x-axis) and its reconstructed frequency *after* sorting $\Phi_i$ (y-axis), which results from summing up its relative frequencies calculated for each sorted cell population. See Supplementary Notes 2-3 and Supplemetary Table S2 for details. The correlation is estimated with the Spearman's coefficient $\rho$. The presented data correspond to the clones that are supported by more than 2 reads in each of the four sorted *TM6** populations (2 replicates × 2 carbon sources) *and* that are found in two input *TM6** populations (2 replicates), therefore, 12,348 clones in R1 and 12,413 in R2. **B.** Correlation between GFP/mCherry ratios assigned to each clone $\mu_i$ in two independent sorting experiments (R1, R2) of C45-HHRz library in *TM6** background cultured either on maltose or glucose. The presented data correspond to 15,031 clones that are supported by 4 reads or more in each of the four sorted TM6 populations (2 replicates × 2 carbon sources). In correlation plots in **A** and **B**, the x-y space has been binned into discrete hexagons, and the color of a hexagon denotes the number of clones lying within. A calibrated color-scale reference is indicated on the right of each panel. The red points indicate the 11 hits identified by the high-throughput screening. Note that 9 out of 11 hits were identified in input populations, and that on the top plot in B some red points overlap.

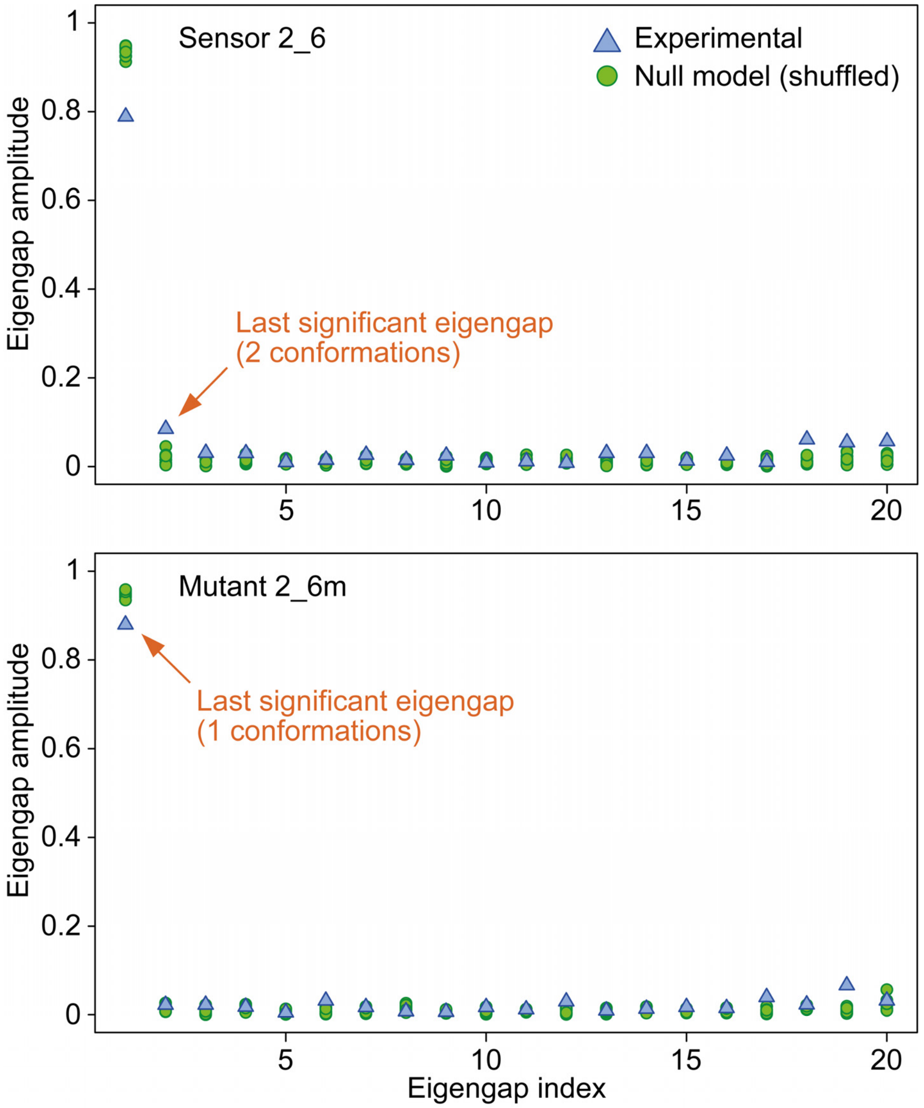


**Supplementary Figure S9. The sensor 2_6 has two conformations inside the cell whereas the mutant 2_6m folds in only one conformation.** Eigengap plots of the differences in magnitude between consecutive eigenvalues. Spectral analysis has been performed using the set of reads covering positions 4-70 (relative to the C45-HHRz RNA device). Eigenvalues are taken from the normalized graph Laplacian matrix constructed using 20 reactive adenosine and cytosine residues.

**SUPPLEMENTARY TABLES**

**Supplementary Table S1.** The sequences of the clones retrieved from SELEX round 13. Boxes split clones with WUCCU motif (in bold) into 3 sets according to sequence similarity in the downstream sequence (highlighted in red, blue and green). The affinity of individual clones representative of each set (highlighted with the same color) was tested by *in vitro* binding assays (Fig. 1 and Supplementary Fig. S1).

| Clone | Constant 5' | Variable | Constant 3' |
|---|---|---|---|
| **C58, C18** | GGGAGGACGAUGCGG | **AUCCU**UAAAG | GACGACUCGCUGAGGAUCCGAGA |
| **C25** | GGGAGGACGAUGCGG | **AUCCU**UAAA | GACGACUCGCUGAGGAUCCGAGA |
| **C77** | GGGAGGACGAUGCGG | **AUCCU**UA | GACGACUCGCUGAGGAUCCGAGA |
| **C1, C2, C50** | GGGAGGACGAUGCGG | **AUCCU** | GACGACUCGCUGAGGAUCCGAGA |
| **C34** | GGGAGGACGAUGCGG | **AUCCU**AGUAC | GACGACUCGCUGAGGAUCCGAGA |
| **C73** | GGGAGGACGAUGCGG | **AUCCU**AGUA | GACGACUCGCUGAGGAUCCGAGA |
| **C65** | GGGAGGACGAUGCGG | **AUCCU**ACU | GACGACUCGCUGAGGAUCCGAGA |
| **C45, C20** | GGGAGGACGAUGCGG | **UUCCU**CAUGUGUU | GACGACUCGCUGAGGAUCCGAGA |
| **C43** | GGGAGGACGAUGCGG | GAAUUAAUGAGG | GACGACUCGCUGAGGAUCCGAGA |
| **C3, C31, C46** | GGGAGGACGAUGCGG | AAU | GACGACUCGCUGAGGAUCCGAGA |

**Supplementary Table S2.** Formal concepts and measures used in the analysis of cell sorting and NGS data. We present them in the order of deriving each clone's response to the change between carbon sources. The values of the measures of the clones studied in the high-throughput screening are stored in *Supplementary files*.

| ***Concepts used in the analysis of the input and sorted populations*** | |
|---|---|
| $\boldsymbol{s}$ | Sample analyzed by sequencing. In case of the input data, each sample corresponds to an experimental replicate and a strain: $s = (r, x)$, where $r \in \{R1, R2\}$ and $x \in \{TM6^*, WT\}$. In case of the sorted data, each sample is also defined by a carbon source and a bin from which the sample is taken in sorting: $s = (r, x, c, b)$, where $c \in \{Mal, Glc\}$ and $b \in \{b1, b2, b3, b4, b5, b6\}$. |
| $\boldsymbol{n_i^s}$ | Coverage (number of reads) of the clone $i$ in the sample $s$ |
| $\boldsymbol{N^s = \sum_i n_i^s}$ | Total coverage in the sample $s$ |
| $\boldsymbol{f_i^s = \frac{n_i^s}{N^s}}$ | Frequency of the clone $i$ in the sample $s$ |

| ***Concepts used in the analysis of the sorted populations only*** Further, we focus on a population $(\boldsymbol{r}, \boldsymbol{x}, \boldsymbol{c})$, from which cells were taken in sorting, and use $\boldsymbol{b}$ instead of $\boldsymbol{s} = (\boldsymbol{r}, \boldsymbol{x}, \boldsymbol{c}, \boldsymbol{b})$. | |
|---|---|
| $\boldsymbol{p^b}$ | Fraction of cells taken from the bin $b$ in sorting |
| $\boldsymbol{\varphi_i^b = f_i^b \cdot p^b}$ | Probability of the cell both expressing the clone $i$ and being taken from the bin $b$ in sorting |
| $\boldsymbol{\Phi_i = \sum_b \varphi_i^b}$ | Probability of the cell expressing the clone $i$ to be sorted, that is, reconstructed frequency of the clone $i$ in the sorted population |
| $\boldsymbol{F_i^b = \frac{\varphi_i^b}{\Phi_i}}$ | Probability of the cell expressing the clone $i$ to be sorted from the bin $b$ rather than from any other bin |
| $\boldsymbol{m^b}$ | Median GFP/mCherry ratio in the bin $b$ |
| $\boldsymbol{\mu_i^{(r,x,c)} = \sum_b m^b \cdot F_i^b}$ | Estimated mean value of the GFP/mCherry ratio of the cells expressing the clone $i$, that is, the readout of the clone $i$ potentially reflecting FBP concentration (in the condition $(r, x, c)$) |
| $\boldsymbol{D_i^{(r,x)} = lg\frac{\mu_i^{(r,x,Glc)}}{\mu_i^{(r,x,Mal)}}}$ | Response of the clone $i$ to the change between the carbon sources |

**Supplementary Table S3.** Regular expressions (search patterns) to classify the sequences obtained in NGS. The regular expressions were applied using Python's module *regex*. Sequence matches were allowed to have up to a particular number of errors including insertions, deletions and substitutions. The number of allowed errors is specified either for the whole regular expression or for its parts in the form of {e<=*i*}.

| Sequence type | | Regular expression | Allowed errors |
|---|---|---|---|
| **Cross-contamination control** | | TCCGGTACGTGAGGTCC | 1 nucleotide (5%) |
| **Diversity control** | | ACAAACAAAGCTGTCAC**[ATGC]{17}**GAAACAGCAAAAAGAAAAATAAAA | 6 nucleotides (10%) |
| **Potential C45-HHRz clone** | | (ACAAACAAAGCTGTCAC){e<=*i*}(**[ATGC]{23,}**)(GAAACAGCAAAAAGAAAAATAAAA){e<=*i*}. If multiple matches are found, the longest is chosen. The middle element **[ATGC]{23,}** is used for further analysis, including the identification of the RNA device architecture (below). | Iterating across *i*∈{0,1,2,3,4,5,6} until a match is found. |
| **RNA device architecture** | **inv-II** | CGGA**[ATGC]{7}**TCCGGTCTGATGAGTCCACTCGCTGAGGATCCGACGGTTCCTCATGTGTTGACGAAGGAC | 5 nucleotides (7-11%) |
| | **C45-II** | CGGA**[ATGC]{7}**TCCGGTCTGATGAGATCCTCATGTGTTGACGACTCGCTGAGGATC | |
| | **sh-II** | CGGA**[ATGC]{7}**TCCGGTCTGATGACTCATGTGTTGACGACTCGCTGA | |
| | **inv-I** | CGGAACTCGCTGAGGATCCGACGGTTCCTCATGTGTTGACGAATCCGGTCTGATGAGTCC**[ATGC]{4}**GGAC | |
| | **C45-I** | GGATCCTCATGTGTTGACGACTCGCTGAGGATCCGTCTTGATGAGTCC**[ATGC]{4}**GGAC | |
| | **sh-I** | CTCATGTGTTGACGACTCGCTGAGGTCTGATGAGTCC**[ATGC]{4}**GGAC | |

**Supplementary Table S4.** The frequency of unique occurrences and the fraction regarding the total coverage for the sequences failing to match the patterns of the cross-contamination control, diversity control and potential C45-HHRz clone as defined in Supplementary Table S3. The samples are defined in Supplementary Table S2.

| *Statistic* | **All 52 samples** |
|---|---|
| **Median** | 1.5e-02, 1.3e-03 |
| **Mean** | 1.5e-02, 1.5e-03 |
| **Q1** | 1.3e-02, 1.3e-03 |
| **Q3** | 1.8e-02, 1.5e-03 |

| *Input sample* | |
|---|---|
| **R1,TM6** | 1.9e-02, 1.3e-03 |
| **R2,TM6** | 1.8e-02, 1.3e-03 |
| **R1,WT** | 2.1e-02, 1.2e-03 |
| **R2,WT** | 2.1e-02, 1.3e-03 |

| *Sorted sample* | **b1** | **b2** | **b3** | **b4** | **b5** | **b6** |
|---|---|---|---|---|---|---|
| **R1,TM6,Glc** | 1.7e-02, 3.4e-03 | 1.1e-02, 1.4e-03 | 1.3e-02, 1.4e-03 | 1.4e-02, 1.3e-03 | 1.9e-02, 1.3e-03 | 1.7e-02, 3.4e-03 |
| **R1,TM6,Mal** | 1.4e-02, 2.7e-03 | 1.0e-02, 1.6e-03 | 1.2e-02, 1.2e-03 | 1.3e-02, 1.3e-03 | 1.5e-02, 1.2e-03 | 1.4e-02, 2.7e-03 |
| **R1,WT,Glc** | 1.6e-02, 1.5e-03 | 1.0e-02, 1.3e-03 | 1.2e-02, 1.2e-03 | 1.4e-02, 1.3e-03 | 1.7e-02, 1.2e-03 | 1.6e-02, 1.5e-03 |
| **R1,WT,Mal** | 1.8e-02, 1.6e-03 | 1.0e-02, 1.1e-03 | 1.3e-02, 1.3e-03 | 1.4e-02, 1.2e-03 | 1.8e-02, 1.2e-03 | 1.8e-02, 1.6e-03 |
| **R2,TM6,Glc** | 1.4e-02, 4.7e-03 | 9.8e-03, 1.3e-03 | 1.3e-02, 1.3e-03 | 1.4e-02, 1.3e-03 | 2.0e-02, 1.3e-03 | 1.4e-02, 4.7e-03 |
| **R2,TM6,Mal** | 1.3e-02, 2.4e-03 | 1.0e-02, 1.4e-03 | 1.2e-02, 1.3e-03 | 1.4e-02, 1.3e-03 | 1.7e-02, 1.3e-03 | 1.3e-02, 2.4e-03 |
| **R2,WT,Glc** | 2.0e-02, 1.8e-03 | 1.0e-02, 1.3e-03 | 1.2e-02, 1.2e-03 | 1.4e-02, 1.4e-03 | 1.9e-02, 1.3e-03 | 2.0e-02, 1.8e-03 |
| **R2,WT,Mal** | 2.1e-02, 2.7e-03 | 1.2e-02, 1.3e-03 | 1.3e-02, 1.2e-03 | 1.5e-02, 1.2e-03 | 1.8e-02, 1.1e-03 | 2.1e-02, 2.7e-03 |

**Supplementary Table S5.** Cross-contamination rate measured as the coverage of the sequences containing the pattern of cross-contamination control (Supplementary Table S3) divided by the total coverage in a sample. The samples are defined in Supplementary Table S2.

| ***Statistic*** | **All 52 samples** |
|---|---|
| **Median** | 4.2e-06 |
| **Mean** | 5.0e-05 |
| **Q1** | 1.1e-06 |
| **Q3** | 1.0e-05 |

| ***Input sample*** | |
|---|---|
| **R1,TM6** | 9.9e-06 |
| **R2,TM6** | 2.7e-05 |
| **R1,WT** | 5.5e-06 |
| **R2,WT** | 7.1e-06 |

| ***Sorted sample*** | **b1** | **b2** | **b3** | **b4** | **b5** | **b6** |
|---|---|---|---|---|---|---|
| **R1,TM6,Glc** | 1.4e-04 | 0 | 6.6e-06 | 9.0e-07 | 8.2e-07 | 5.3e-05 |
| **R1,TM6,Mal** | 0 | 4.5e-06 | 7.2e-06 | 3.8e-06 | 0 | 3.4e-05 |
| **R1,WT,Glc** | 0 | 2.0e-06 | 2.6e-06 | 0 | 3.5e-06 | 5.5e-05 |
| **R1,WT,Mal** | 6.3e-06 | 2.8e-06 | 4.0e-06 | 0 | 2.7e-06 | 1.1e-05 |
| **R2,TM6,Glc** | 1.9e-03 | 6.9e-06 | 2.4e-06 | 3.6e-06 | 1.5e-06 | 3.2e-05 |
| **R2,TM6,Mal** | 0 | 1.1e-06 | 5.4e-06 | 0 | 2.5e-06 | 9.9e-06 |
| **R2,WT,Glc** | 1.0e-06 | 1.5e-05 | 6.0e-06 | 5.3e-05 | 1.4e-05 | 2.5e-05 |
| **R2,WT,Mal** | 0 | 9.2e-07 | 6.1e-06 | 2.7e-06 | 9.2e-06 | 1.1e-04 |

**Supplementary Table S6.** NGS reads per unique plasmid molecule. The values are derived using the statistics of the diversity-control sequences spiked into each sample at the point of the outer PCR before sequencing. The diversity-control sequences are identified via the pattern presented in Supplementary Table S3, and the median coverage across these sequences is calculated in each sample. Due to the paired-end sequencing, the number of reads is divisible by 2, with the smallest value being 2. The samples are defined in Supplementary Table S2.

| ***Statistic*** | **All 52 samples** |
|---|---|
| **Median** | 2 |
| **Mean** | 2.4 |
| **Q1** | 2 |
| **Q3** | 2 |

| ***Input sample*** | |
|---|---|
| **R1,TM6** | 2 |
| **R2,TM6** | 2 |
| **R1,WT** | 2 |
| **R2,WT** | 2 |

| ***Sorted sample*** | **b1** | **b2** | **b3** | **b4** | **b5** | **b6** |
|---|---|---|---|---|---|---|
| **R1,TM6,Glc** | 4 | 2 | 2 | 2 | 2 | 4 |
| **R1,TM6,Mal** | 2 | 2 | 2 | 2 | 2 | 2 |
| **R1,WT,Glc** | 2 | 2 | 2 | 2 | 2 | 4 |
| **R1,WT,Mal** | 2 | 2 | 2 | 2 | 2 | 4 |
| **R2,TM6,Glc** | 4 | 2 | 2 | 2 | 2 | 4 |
| **R2,TM6,Mal** | 2 | 2 | 2 | 2 | 2 | 4 |
| **R2,WT,Glc** | 2 | 2 | 2 | 2 | 4 | 4 |
| **R2,WT,Mal** | 2 | 2 | 2 | 2 | 2 | 4 |

**Supplementary Table S7.** Oligonucleotides used in this study

| Code | Primer ID | Sequence (5´- 3') | Remarks |
|---|---|---|---|
| **ADO01** | T7-C34-F | ccagaTAATACGACTCACTATAGGGAGGACGATGCGGATCCTAGTACGACGACTCGCTGAGGATCCGAGA | Template for in vitro transcription (binding assays) |
| **ADO02** | T7-C34-R | TCTCGGATCCTCAGCGAGTCGTCGTACTAGGATCCGCATCGTCCTCCCTATAGTGAGTCGTATTAtctgg | Template for in vitro transcription (binding assays) |
| **ADO03** | T7-C34mut1-F | ccagaTAATACGACTCACTATAGGGAGGACGATGGCCTAGGATGTACCAGACGACTCGCTGAGGATCCGAGA | Template for in vitro transcription (binding assays) |
| **ADO04** | T7-C34mut1-R | TCTCGGATCCTCAGCGAGTCGTCTGGTACATCCTAGGCCATCGTCCTCCCTATAGTGAGTCGTATTAtctgg | Template for in vitro transcription (binding assays) |
| **ADO05** | C45-F | ccagaTAATACGACTCACTATAGGGAAAAGGACGATGCGGTTCCTCATGTGTTGACGACTCGCTGAGGATCCGAGA | Template for in vitro transcription (binding assays) |
| **ADO06** | C45-R | TCTCGGATCCTCAGCGAGTCGTCAACACATGAGGAACCGCATCGTCCTTTTCCCTATAGTGAGTCGTATTAtctgg | Template for in vitro transcription (binding assays) |
| **ADO07** | C45P2-F | ccagaTAATACGACTCACTATAGGGAAATCCTCATGTGTTGACGACTCGCTGAGGA | Template for in vitro transcription (binding assays) |
| **ADO08** | C45P2-R | TCCTCAGCGAGTCGTCAACACATGAGGATTTCCCTATAGTGAGTCGTATTAtctgg | Template for in vitro transcription (binding assays) |
| **ADO09** | mutL-2-F | ccagaTAATACGACTCACTATAGGGAGGACGATGCGGTTCCTCATGTGAACTGCTGTCGCTGAGGATCCGAGA | Template for in vitro transcription (binding assays) |
| **ADO10** | mutL-2-R | TCTCGGATCCTCAGCGACAGCAGTTCACATGAGGAACCGCATCGTCCTCCCTATAGTGAGTCGTATTAtctgg | Template for in vitro transcription (binding assays) |
| **ADO11** | mutB-F | ccagaTAATACGACTCACTATAGGGAGGACGATGCGGTTCCTCAGTGTTGACGACTCGCTGAGGATCCGAGA | Template for in vitro transcription (binding assays) |
| **ADO12** | mutB-R | TCTCGGATCCTCAGCGAGTCGTCAACACTGAGGAACCGCATCGTCCTCCCTATAGTGAGTCGTATTAtctgg | Template for in vitro transcription (binding assays) |
| **ADO13** | APT-pWHE-F | tccaagctagatctcttaagGGGAGGACGATGCGG | Cloning C45 into pWHE601 |
| **ADO14** | APT-pWHE-R | tcttctcctttgctagccatTTTTTCTCGGATCCTCAGCGAGT | Cloning C45 into pWHE602 |
| **ADO15** | pWHE601-H2-F | atggctagcaaaggagaaga | Cloning C45 into pWHE603 |
| **ADO16** | pWHE601-H1-R | cttaagagatctagcttggagttga | Cloning C45 into pWHE604 |
| **ADO17** | SHAPE-1cas-F | GGCCTTCGGGCAAGGGAGGACGATGCGGTTCC | Template for in vitro transcription (SHAPE) |
| **ADO18** | GFP300-R | cacgcgtcttgtagttcccg | Template for in vitro transcription (SHAPE) |
| **ADO19** | SHAPE-T7-2cas-F | CCAGATAATACGACTCACTATAGGGCCTTCGGGCAAAAGGCCTTCGGGCCAAA | Template for in vitro transcription (SHAPE) |
| **ADO20** | GFP200-R | ggaaaagcattgaacacc | Template for in vitro transcription (SHAPE) |
| **ADO21** | SHAPE-T7 | CCAGATAATACGACTCACTATAGGG | Template for in vitro transcription (SHAPE) |
| **ADO22** | GFP100-R | ccgtatgtagcatcaccttcac | Primer extension (SHAPE) |
| **ADO23** | HHRz-F | GACCTAGGAAACAAACAAAGCTGTCACCGGATGTGCTTTCCGGTCTGATGAGTCCG | Cloning sTRSV hammerhead ribozyme, positive control in *in vivo* reporter system |
| **ADO24** | HHRz-R | GGCTCGAGTTTTTATTTTTCTTTTTGCTGTTTCGTCCTCACGGACTCATCAGACCGGAAAG | Cloning sTRSV hammerhead ribozyme, positive control in *in vivo* reporter system |

| | | | |
|---|---|---|---|
| **ADO25** | HHRz-CTL | GACCTAGGAAACAAACAAAGCtGtCACCGGAtGtGCtttCCGGtACGtGAGGtCCGtGAGGACGAAACAGCAAAAAGAAAAAtAAAAACTCGAGCC | Cloning inactive hammerhead ribozyme mutant, negative control in *in vivo* reporter system |
| **ADO26** | HHRz-M1 | GACCTAGGAAACAAACAAAGCtGtCACCGGATATGtCCGGtCtGAtGAGtCCAGAAGGACGAAACAGCAAAAAGAAAAAtAAAAACTCGAGCC | Cloning hammerhead ribozyme loop mutants, calibration in *in vivo* reporter system |
| **ADO27** | HHRz-M2 | GACCTAGGAAACAAACAAAGCtGtCACCGGATGTTtCCGGtCtGAtGAGtCCACTAGGACGAAACAGCAAAAAGAAAAAtAAAAACTCGAGCC | Cloning hammerhead ribozyme loop mutants, calibration in *in vivo* reporter system |
| **ADO28** | HHRz-M3 | GACCTAGGAAACAAACAAAGCtGtCACCGGATTGTtCCGGtCtGAtGAGtCCCATAGGACGAAACAGCAAAAAGAAAAAtAAAAACTCGAGCC | Cloning hammerhead ribozyme loop mutants, calibration in *in vivo* reporter system |
| **ADO29** | HHRz-M4 | GACCTAGGAAACAAACAAAGCtGtCACCGGAGGCTtCCGGtCtGAtGAGtCCAGCTGGACGAAACAGCAAAAAGAAAAAtAAAAACTCGAGCC | Cloning hammerhead ribozyme loop mutants, calibration in *in vivo* reporter system |
| **ADO30** | HHRz-M5 | GACCTAGGAAACAAACAAAGCtGtCACCGGATGCAtCCGGtCtGAtGAGtCCCGTTGGACGAAACAGCAAAAAGAAAAAtAAAAACTCGAGCC | Cloning hammerhead ribozyme loop mutants, calibration in *in vivo* reporter system |
| **ADO31** | HHRz-M6 | GACCTAGGAAACAAACAAAGCtGtCACCGGACAGGtCCGGtCtGAtGAGtCCAGTTGGACGAAACAGCAAAAAGAAAAAtAAAAACTCGAGCC | Cloning hammerhead ribozyme loop mutants, calibration in *in vivo* reporter system |
| **ADO32** | HHRz-M7 | GACCTAGGAAACAAACAAAGCtGtCACCGGATGCAtCCGGtCtGAtGAGtCCCGCGGGACGAAACAGCAAAAAGAAAAAtAAAAACTCGAGCC | Cloning hammerhead ribozyme loop mutants, calibration in *in vivo* reporter system |
| **ADO33** | C45_I-F | GACCTAGGAAACAAACAAAgctgtcaCGGATCCTCATGTGTTGACGACTCGCTGAGGATCCGTC | Cloning C45-HHRz library. Receiving stem-loop: stem I; randomized loop: stem-loop II; aptamer version: C45 |
| **ADO34** | C45_I-R | GGCTCGAGTTTTTATTTTTCTTTTTgctgtttcgtccNNNNggactcatcaAGACGGATCCTCAGCGAGTCGTC | Cloning C45-HHRz library. Receiving stem-loop: stem I; randomized loop: stem-loop II; aptamer version: C45 |
| **ADO35** | C45_II-F | GACCTAGGAAACAAACAAAgctgtcaccggaNNNNNNNtcCGGTCTGATGAGATCCTCATGTGTTG | Cloning C45-HHRz library. Receiving stem-loop: stem II; randomized loop: stem-loop I; aptamer version: C45 |
| **ADO36** | C45_II-R | GGCTCGAGTTTTTATTTTTCTTTTTgctgtttcGATCCTCAGCGAGTCGTCAACACATGAGGATCtcatcagaccg | Cloning C45-HHRz library. Receiving stem-loop: stem II; randomized loop: stem-loop I; aptamer version: C45 |
| **ADO37** | C45inv2-I-F | GACCTAGGAAACAAACAAAgctgtcaccggaaCTCGCTGAGGATCCGACGGTTCCTCATGTGTTGACGAAtccg | Cloning C45-HHRz library. Receiving stem-loop: stem I; randomized loop: stem-loop II; aptamer version: inverted C45 |
| **ADO38** | C45inv2-I-R | GGCTCGAGTTTTTATTTTTCTTTTTgctgtttcgtccNNNNggactcatcagaccggaTTCGTCAACACATGAGGAACCGTC | Cloning C45-HHRz library. Receiving stem-loop: stem I; randomized loop: stem-loop II; aptamer version: inverted C45 |
| **ADO39** | C45inv2-II-F | GACCTAGGAAACAAACAAAgctgtcaccggaNNNNNNNtccggtctgatgagtccaCTCGCTGAGGATCCGACGGTTCC | Cloning C45-HHRz library. Receiving stem-loop: stem II; randomized loop: stem-loop I; aptamer version: inverted C45 |
| **ADO40** | C45inv2-II-R | GGCTCGAGTTTTTATTTTTCTTTTTgctgtttcgtccTTCGTCAACACATGAGGAACCGTCGGATCCTCAGCGAGtggactc | Cloning C45-HHRz library. Receiving stem-loop: stem II; randomized loop: stem-loop I; aptamer version: inverted C45 |
| **ADO41** | HHRz-C45sh1-I | GACCTAGGAAACAAACAAAGCtGtCaCCTCATGTGTTGACGACTCGCTGAGGtCtGAtGAGtCCNNNNGGACGAAACAGCAAAAAGAAAAAtAAAAACTCGAGCC | Cloning C45-HHRz library. Receiving stem-loop: stem I; randomized loop: stem-loop II; aptamer version: short C45 |
| **ADO42** | HHRz-C45sh2-II | GACCTAGGAAACAAACAAAGCtGtCACCGGANNNNNNNtCCGGtCtGAtGACTCATGTGTTGACGACTCGCTGAGGAAACAGCAAAAAGAAAAAtAAAAACTCGAGCC | Cloning C45-HHRz library. Receiving stem-loop: stem II; randomized loop: stem-loop I; aptamer version: short C45 |
| **ADO43** | Lib-HHRz-F | GACCTAGGAAACAAACAAAGCTGTCACC | Cloning C45-HHRz library and hammerhead ribozyme versions. Primers for amplification. |

| | | | |
|---|---|---|---|
| **ADO44** | Lib-HHRz-R | GGCTCGAGTTTTTATTTTTCTTTTTGC TGTTTCG | Cloning C45-HHRz library and hammerhead ribozyme versions. Primers for amplification. |
| **ADO45** | GFP_seq-F | CTGCTGCTGGTATTATCCATGGTATG | Cloning C45-HHRz library and hammerhead ribozyme versions. Primers for sequencing. |
| **ADO46** | ADHt_seq-R | CATAAGAAATTCGCTTATTTAGAAGTGGC | Cloning C45-HHRz library and hammerhead ribozyme versions. Primers for sequencing. |
| **ADO47** | Out-Nex1-HH-F | CCTAGGAAACAAACAAAGCTG | NGS library. Outer PCR. |
| **ADO48** | Out-Nex1-HH-R | CTCGAGTTTTTATTTTTCTTTTTGCTG | NGS library. Outer PCR. |
| **ADO49** | DivC-HHI-tmp | CCTAGGAAACAAACAAAgctgtcacNNNNNNN NNNNNNNNNNgaaacagcAAAAAGAAAAATA AAAACTCGAG | NGS library. Diversity control. |
| **ADO50** | Nex1-HHsh-F | TCGTCGGCAGCGTCAGATGTGTATAAGAGAC AGACAAACAAAGCTGTCAC | NGS library. Nextera PCR. |
| **ADO51** | Nex1-HHsh-R | GTCTCGTGGGCTCGGAGATGTGTATAAGAGA CAGTTTTATTTTTCTTTTTGCTGTTTC | NGS library. Nextera PCR. |
| **ADO52** | Control-1_10-F | GACCTAGGAAACAAACAAAGCTGTCACCGGA TTTCAGTTCCGGTCTGATGAGTCCACTCGCTG AGGATCCGACGGTTCC | Cloning sensor candidates for validation. |
| **ADO53** | Control-1_10-R | GGCTCGAGTTTTTATTTTTCTTTTTGCTGTTTC GTCCTTCGTCAACACATGAGGAACCGTCGGAT CCTCAGCGAGTGGACTC | Cloning sensor candidates for validation. |
| **ADO54** | Hit-1_8-F | GACCTAGGAAACAAACAAAGCTGTCACCAGA TTTGTGATCCGGTCTGATGAGTCCACTCGCTG AGGATCCGACGGTTCC | Cloning sensor candidates for validation. |
| **ADO55** | Hit-1_8-R | GGCTCGAGTTTTTATTTTTCTTTTTGCTGTTTC GTCCTTCGTCAACACATGAGGAACCGTCGGAT CCTCAGCGAGTGGACTC | Cloning sensor candidates for validation. |
| **ADO56** | Hit-2_10-F | GACCTAGGAAACAAACAAAGCTGTCACCGGA TTCGTGGTCCGGTCTGATGAGTCCACTCGCTG AGGATCCGACGGTTCC | Cloning sensor candidates for validation. |
| **ADO57** | Hit-2_10-R | GGCTCGAGTTTTTATTTTTCTTTTTGCTGTTTC GTCCTTCATCAACACATGAGGAACCGTCGGAT CCTCAGCGAGTGGACTC | Cloning sensor candidates for validation. |
| **ADO58** | Hit-2_12 | GACCTAGGAAACAAACAAAGCTGTACCGGAG GAGGGGTCCGGTCTGATGACTCCTGTGTTGA CGACTCGCTGAGGAAACAGCAAAAAGAAAAA TAAAAACTCGAGCC | Cloning sensor candidates for validation. |
| **ADO59** | Hit-2_2-F | GACCTAGGAAACAAACAAAGCTGTCACGGAG TAAGGCTCCGGTCTGATGAGATCCTCATGTGT TG | Cloning sensor candidates for validation. |
| **ADO60** | Hit-2_2-R | GGCTCGAGTTTTTATTTTTCTTTTTGCTGTTTC GATCCTCAGCGAGTCGTCAACACATGAGGAT CTCATCAGACCG | Cloning sensor candidates for validation. |
| **ADO61** | Hit-2_3-F | GACCTAGGAAACAAACAAAGCTGTCACGGAG TATGGCTCCGGTCTGATGAGATCCTCATGTGT TG | Cloning sensor candidates for validation. |
| **ADO62** | Hit-2_3-R | GGCTCGAGTTTTTATTTTTCTTTTTGCTGTTTC GATCCTCAGCGAGTCGTCAACACATGAGGAT CTCATCAGACCG | Cloning sensor candidates for validation. |
| **ADO63** | Hit-2_6-F | GACCTAGGAAACAAACAAAGCTGTCACCGGA ATAGGAGTCCGGTCTGATGAGTCCACTTGCT GAGGATCCGACGGTTCC | Cloning sensor candidates for validation. |
| **ADO64** | Hit-2_6-R | GGCTCGAGTTTTTATTTTTCTTTTTGCTGTTTC GTCCTTCGTCAACACATGAGGAACCGTCGGAT CCTCAGCAAGTGGACTC | Cloning sensor candidates for validation. |

| | | | |
|---|---|---|---|
| **ADO65** | Hit-2_7-F | GACCTAGGAAACAAACAAAGCTGTCACCGGAATGGGGGTCCGGTCTGATGAGTCCACTCGCTGAGGATCCGACGGTTCC | Cloning sensor candidates for validation. |
| **ADO66** | Hit-2_7-R | GGCTCGAGTTTTTATTTTTCTTTTTGCTGTTTCAGTCCTTCGTCAACACATGAGGAACCGTCGGATCCTCAGCGAGTGGACTC | Cloning sensor candidates for validation. |
| **ADO67** | Hit-2_8-F | GACCTAGGAAACAAACAAAGCTGTCACCGGAATGGGGGTCCGGTCTGATGAGTCCACTCGCTGAGGATCCGACGGTTCC | Cloning sensor candidates for validation. |
| **ADO68** | Hit-2_8-R | GGCTCGAGTTTTTATTTTTCTTTTTGCTGTTTCGTACTTCGTCAACACATGAGGAACCGTCGGATCCTCAGCGAGTGGACTC | Cloning sensor candidates for validation. |
| **ADO69** | Hit-3_7 | GACCTAGGAAACAAACAAAGCTGTACCGGAGGAGGGGTCCGGTCTGAGGACTCATGTGTTGACGACTCGCTGAGGAAACAGCAAAAAGAAAAATAAAAACTCGAGCC | Cloning sensor candidates for validation. |
| **ADO70** | Hit-4_1-F | GACCTAGGAAACAAACAAAGCTGTCACGGAGTGGGGTTCCGGTCTGATGAGATCCTCATGTGTTG | Cloning sensor candidates for validation. |
| **ADO71** | Hit-4_1-R | GGCTCGAGTTTTTATTTTTCTTTTTGCTGTTTCGATCCTCAGCGAGTCGTCAACACATGAGGATCTCATCAGACCG | Cloning sensor candidates for validation. |
| **ADO72** | Mutant-2_6m-F | GACCTAGGAAACAAACAAAGCTGTCACCGGAATAGGAGTCCGGTCTGATGAGTCCTGTTGCTGAGGATCCGACGGTTCC | Cloning sensor candidates for validation. |
| **ADO73** | Mutant-2_6m-R | GGCTCGAGTTTTTATTTTTCTTTTTGCTGTTTCGTCCAAGCAGTTCACATGAGGAACCGTCGGATCCTCAGCAACAGGACTC | Cloning sensor candidates for validation. |
| **ADO74** | Mutant 3_7m | GACCTAGGAAACAAACAAAGCTGTACCGGAGGAGGGGTCCGGTCTGAGGACTCATGTGAACTGCTGTCGCTGAGGAAACAGCAAAAAGAAAAATAAAAACTCGAGCC | Cloning sensor candidates for validation. |
| **ADO75** | Mutant-4_1m-F | GACCTAGGAAACAAACAAAGCTGTCACGGAGTGGGGTTCCGGTCTGATGAGATCCTCATGTGAAC | Cloning sensor candidates for validation. |
| **ADO76** | Mutant-4_1m-R | GGCTCGAGTTTTTATTTTTCTTTTTGCTGTTTCGATCCTCAGCGACAGCAGTTCACATGAGGATCTCATCAGACCG | Cloning sensor candidates for validation. |
| **ADO77** | M13-R | CAGGAAACAGCTATGACCATG | Sequencing sensor candidates clones. |
| **ADO78** | DMS-MaPseq-F | TGGTGATGGTCCAGTCTTGTT | cDNA amplification for *in vivo* structural probing. |
| **ADO79** | DMS-MaPseq-R | AGAAGTGGCGCGCCCT | Reverse transcription and cDNA amplification for *in vivo* structural probing. |
